# Highly reproducible characterization of *Escherichia coli* tRNA epitranscriptome with a simple method of library preparation for deep sequencing

**DOI:** 10.1101/2020.02.21.958157

**Authors:** Ji Wang, Claire Toffano-Nioche, Florence Lorieux, Daniel Gautheret, Jean Lehmann

**Affiliations:** Institute for Integrative Biology of the Cell (I2BC), CEA, CNRS, Université Paris-Sud, Campus Paris-Saclay, 91198 Gif-sur-Yvette, France

## Abstract

In conventional RNA high-throughput sequencing, modified bases prevent a large fraction of tRNA transcripts to be converted into cDNA libraries. Recent proposals aiming at resolving this issue take advantage of the interference of base modifications with RT enzymes to detect and identify them by establishing signals from aborted cDNA transcripts. Because some modifications, such as methyl groups, do almost not allow RT bypassing, demethylation and highly processive RT enzymes have been used to overcome these obstacles. Working with *Escherichia coli* as a model system, we show that with a conventional (albeit still engineered) RT enzyme and key optimizations in library preparation, all RT-impairing modifications can be highlighted along the entire tRNA length without a demethylation procedure. This is achieved by combining deep-sequencing samples, which allows to establish aborted transcription signal of higher accuracy and reproducibility, with the potential for differentiating tiny differences in the state of modification of all cellular tRNAs. In addition, our protocol provides estimates of the relative tRNA abundance.

## INTRODUCTION

The characterization of all cellular tRNAs from either cells or tissues, both in terms of modification state and relative frequency, has attracted much interest during the past few years (Cozen et al. 2015; Zheng et al. 2015; Hauenschield et al. 2015; Gogakos et al. 2017; Shigematsu et al. 2017; Schwartz et al. 2018). It is now well established that a large variety of diseases originate from an absence of tRNA modifications (Bohnsack and Sloan 2017; Jonkhout et al. 2018), while the modification state of tRNAs has also been shown to relate to metabolism (Richter et al. 2018; Ng et al. 2018; Pollo-Oliveira and De Crecy-Lagard 2018). It is, therefore, highly desirable to develop straightforward methods for the characterization of all tRNAs. The presence of modifications on tRNA constitutes a burden in traditional deepsequencing experiments because they prevent RT enzymes from generating full cDNA transcripts by blocking their progression along the RNA strand. Procedures to amplify aborted cDNA transcripts have been developed, which allows establishing signals at the location of RT-impending modifications, and thus acquiring information about the state of modification (Hauenschild et al. 2015, Zheng et al. 2015, Clark et al. 2016). Some modifications, such as methyl groups, do almost not allow any RT bypass when they are situated on the Watson-Crick face of bases. In order to overcome this issue, Cozen et al. (2015) and Zheng et al. (2015) combined demethylase treatments with an exceptionally processive RT enzyme to generate full-lengh cDNA transcripts at higher yields. Alternate procedures were developed by Motorin, Helm and coworkers, who established aborted transcription signals from fragmented RNA (Hauenschild et al. 2015). Although it is still used in many RNA deep-sequencing protocols (Hauenschild et al. 2015; Gogakos et al. 2017), RNA fragmentation generates a high level of noise in the analysis due to ambiguous gene mapping. This procedure was, therefore, avoided in the present study.

Searching for a method that could highlight RT-impairing base modifications with minimal benchwork, we identified a robust procedure that allows to generate signals from aborted cDNA transcripts (we call them termination signals, **ts**) without requiring demethylase treatment and/or RNA fragmentation, and that we validated with a regular (although still engineered) RT enzyme (Superscript III, Invitrogen). Working with 3 biological replicates from *Escherichia coli*, a bacterial species the tRNA modifications of which are well characterized, we first show that conventional treatments of total RNA samples (DNAse, DNAse and deacylation) have no significant effect on **ts** signals. The number of QC-passed reads obtained from single samples (10-15 million) is, however, too low to fully characterize all tRNAs. Next, we show that a significant gain in accuracy of **ts** signals is obtained by combining independently sequenced samples. Because deep-sequencing experiments require PCR amplification, in which tiny variations in the amount of initial sample translates into noticeable differences in relative fractions of cDNA transcripts, **ts** signals fluctuate around characteristic values, these fluctuations reaching amplitudes as high as 90%. Pooling reads from 3 experiments allows us to demonstrate a reduction in **ts** standard deviation by a factor > 2. Furthermore, by combining all samples from 9 independent deep-sequencing experiments, totalizing about 60 million mapped reads, we show that the coverage is high enough to fully characterize 43 out of the 48 different *E. coli* tRNAs, 5 weakly expressed tRNAs being incompletely covered with **ts** signals. *E. coli* being an extensively studied bacteria, almost all **ts** signals can be attributed to modifications listed in the *Modomics* database (Boccaletto et al. 2018). An innovative aspect of our protocol was found essential to obtaining highly consistent data. Because the A76 residue can be specifically removed by RNase T in *E. coli* (Deutscher *et al.* 1984, 1985), a subpopulation of nonfunctional tRNAs may always occur in these cells (this feature was observed in conventional deep-sequencing; Czech 2020 and our unpublished data). Conversely, an additional CCA may be added by the CCA-adding enzyme that tags these tRNAs for degradation through the combined action of poly(A) polymerase and RNase R (Li et al. 2002; Mohanty et al. 2012, Wellner *et al.* 2018). We found that the use of primers with TGG 3’ overhang in the first PCR step essentially only amplifies CCA 3’-ending RNA transcripts and drastically reduces background amplification, enabling our protocol to generate signals of unprecedented cleanliness, although it is restricted to almost only tRNAs. This feature offers an opportunity: because all transcribed cDNAs start at the 3’ end of the tRNAs, a relative quantification of mature isoacceptors can be established from read coverage at that position.

## MATERIAL & METHODS

### Samples preparation

To test the proposed method, three biological replicates (A, B and C) of total RNA extracts from *E. coli MG1655* cells grown at 37°C in standard LB medium were sampled at an OD600 of ~4.0. Each of these samples were prepared following three different procedures (Fig. 1): Total RNA was either not treated (**-**), or treated with DNase (**D**), or treated with DNase and deacylated (**Dd**) prior to 3’ adapter ligation (Table 1). After reverse transcription, chimeric RNA/cDNA products were directly loaded on gel for size selection between 35 to 200 bp by electrophoresis, to eliminate the excess of 3’ adapters and select mid-size RNA transcripts (Fig. 1). Gel-extracted products were subjected to poly-A tailing by a TdT enzyme. A first round of selective pre-amplification of the cDNA was achieved with *Illumina* TruSeq sRNA RTP with TGG overhang and *Illumina* TruSeq sRNA 5’ adapter primer with polyT(20). A second amplification with *Illumina* RP1 and *Illumina* TruSeq sRNA Index primer (RPIX) that added a specific tag to each of the 9 samples (Table 1) provided the final libraries. *Illumina* deep sequencing (libraries multiplexing; pair-end: 2×125bp PE, HiSeq High Output mode, V4 chemistry) was performed by an external contractor (Genewiz). The detailed protocol of library preparation is available in supplemental File S1.

**Figure 1.**
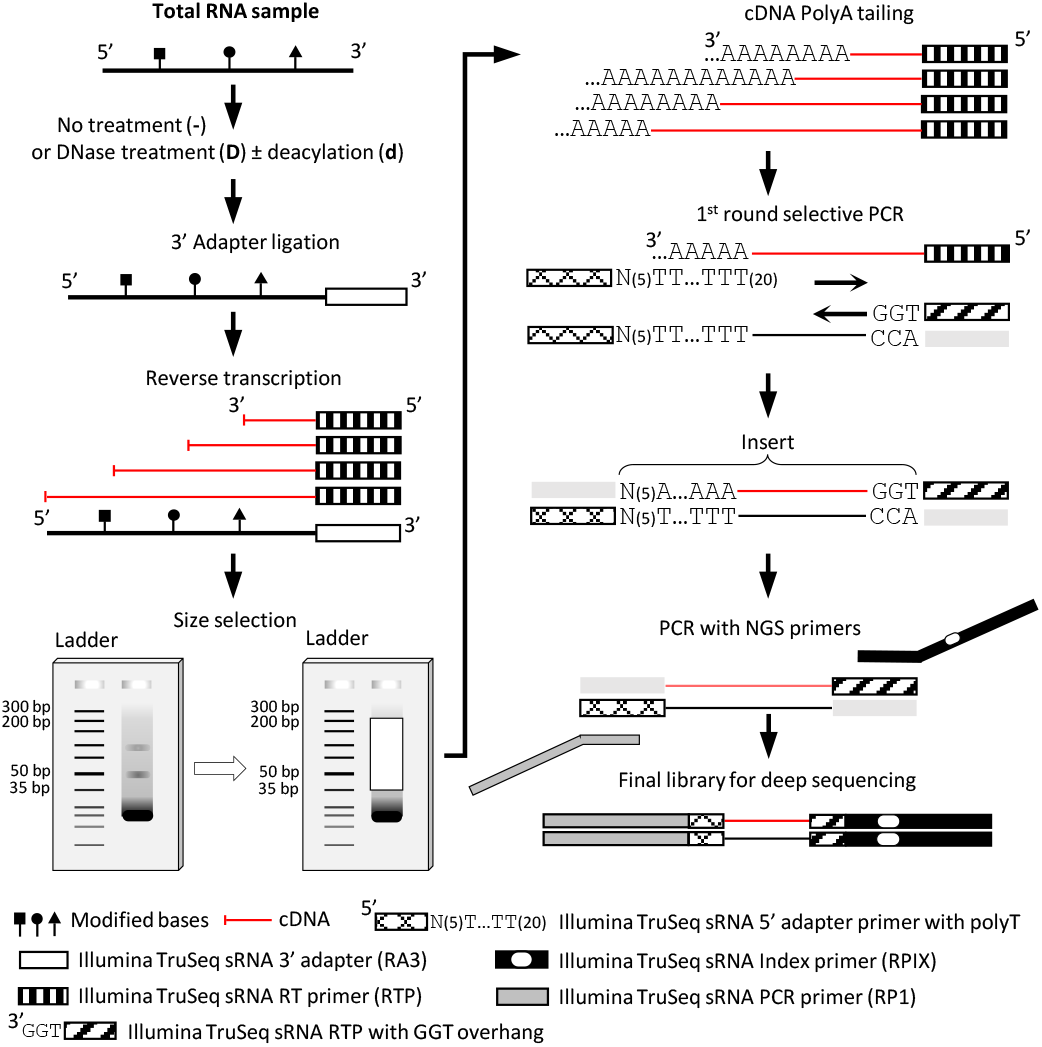
Workflow of library preparation (see text for explanations and Supplemental File S1 for detailed protocol). Poly-A tailing of cDNA was applied on gel-extracted cDNA/RNA chimeric duplexes without RNase treatment. Selective PCR was achieved with 10-18 cycles, while the PCR with NGS primers was programmed with 5-15 cycles. The stretch of 5 random positions (N5) preceding the poly(T) tail of the 5’ adapter primer is a requirement of Illumina deep-sequencing spot localization.

**Table 1:** The 9 total RNA samples analysed in this study.

| Treatment | Biological replicate |  |  |
| --- | --- | --- | --- |
|  | A | B | C |
| No treatment | A- | B- | C- |
| DNase | AD | BD | CD |
| DNase + deacylation | ADd | BDd | CDd |

### Computational treatment of deep-sequencing data

Processing of the deep-sequencing data required an optimization in order to extract the most relevant signal from read coverage. We established a protocol (Suppl. Fig. S1) that generated the sharpest termination signals (see below). In brief, only R2 reads (sequenced from the 3’ end) were processed. Both adapters were removed and poly-A tails were trimmed. Mapping on *E. coli* K-12 (substr. MG1655) genome was achieved with Bowtie 2 (Langmead and Salzberg 2012) with the *--local* option, that maximizes the stretch of correct matches at the end of a read. This option turned out to be essential because bases other than A sometimes occur in the added poly-A that prevent its complete removal at the trimming step. Finally, a CCA/TGG filter complementing the selective PCR step only kept mapped transcripts with CCA at the 3’ end. The quality and distribution of read lengths at each step, performed with *FastQC* (v0.11.5, *Babraham Bioinformatics*), are shown in supplemental Figure S1. After filtering, about 5 million to 9 million reads mapped onto the *E. coli* genome (Table 2; Suppl. Fig. S1), allowing to establishing tRNA genomic coverages.

**Table 2:**
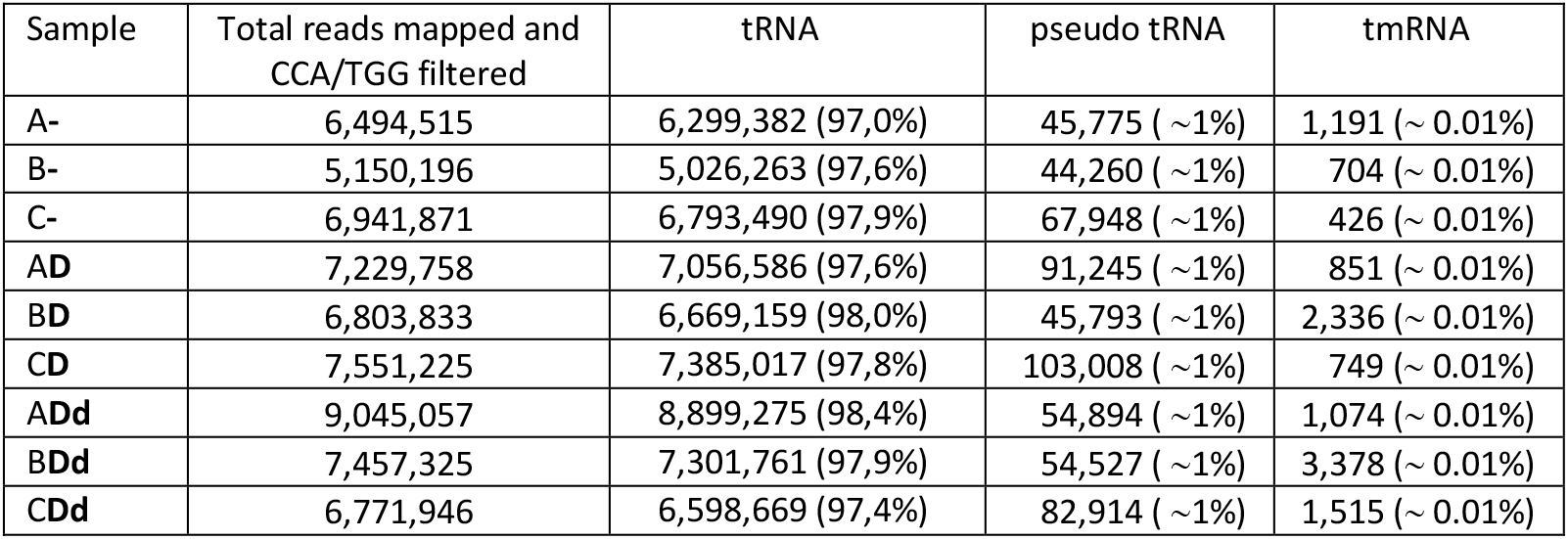
Total read numbers of samples. Reads of tRNA are from the following genes (identical copies are indicated in brackets): *(alaT alaU alaV), (alaX alaW), (argQ argV argY argZ), (argU), (argW), (argX), (asnT asnU asnV asnW), (aspT aspU aspV), (cysT), (glnU glnW), (glnV glnX), (gltT gltU gltV gltW), (glyT), (glyU), (glyV glyW glyX glyY), (hisR), (ileT ileU ileV), (ileX), (ileY), (leuP), (leuQ leuT leuV), (leuU), (leuW), (leuX), (leuZ), (lysQ lysT lysV lysW lysY lysZ), (metT metU), (metV metW metZ), (metY), (pheU pheV), (proK), (proL), (proM), (selC), (serT), (serU), (serV), (serW serX), (thrT), (thrU), (thrV), (thrW), (trpT), (tyrT tyrV), (tyrU), (valT valU valX valY valZ), (valV), (valW)*. Reads of pseudo tRNA are from the following genes: *(pauD), (pawZ), (ptwF).* Reads of tmRNA are from the following gene: *(ssrA).*

| Sample | Total reads mapped and CCA/TGG filtered | tRNA | pseudo tRNA | tmRNA |
| --- | --- | --- | --- | --- |
| A- | 6,494,515 | 6,299,382 (97,0%) | 45,775 ( ~1%) | 1,191 ( ~ 0.01%) |
| B- | 5,150,196 | 5,026,263 (97,6%) | 44,260 ( ~1%) | 704 ( ~ 0.01%) |
| C- | 6,941,871 | 6,793,490 (97,9%) | 67,948 ( ~1%) | 426 ( ~ 0.01%) |
| AD | 7,229,758 | 7,056,586 (97,6%) | 91,245 ( ~1%) | 851 ( ~ 0.01%) |
| BD | 6,803,833 | 6,669,159 (98,0%) | 45,793 ( ~1%) | 2,336 ( ~ 0.01%) |
| CD | 7,551,225 | 7,385,017 (97,8%) | 103,008 ( ~1%) | 749 ( ~ 0.01%) |
| ADd | 9,045,057 | 8,899,275 (98,4%) | 54,894 ( ~1%) | 1,074 ( ~ 0.01%) |
| BDd | 7,457,325 | 7,301,761 (97,9%) | 54,527 ( ~1%) | 3,378 ( ~ 0.01%) |
| CDd | 6,771,946 | 6,598,669 (97,4%) | 82,914 ( ~1%) | 1,515 ( ~ 0.01%) |

### Visualization

To connect **ts** signals with modified bases, all *E. coli* tRNA sequences from the *Modomics* database (Boccaletto et al. 2018) were aligned onto the *E. coli* genome with blastn (ncbi-blast-2.5.0+), where a .gff file specifying the position of 661 modifications (Suppl. File S2) allowed their visualization on IGV (Robinson et al. 2017). A separate .txt file (Suppl. File S3) was used to generate plots connecting **ts** signals with tRNA sequences and modifications (Fig. 2b). Modifications one-letter-code are from the *Modomics* database (supplemental Table S1).

**Figure 2.**
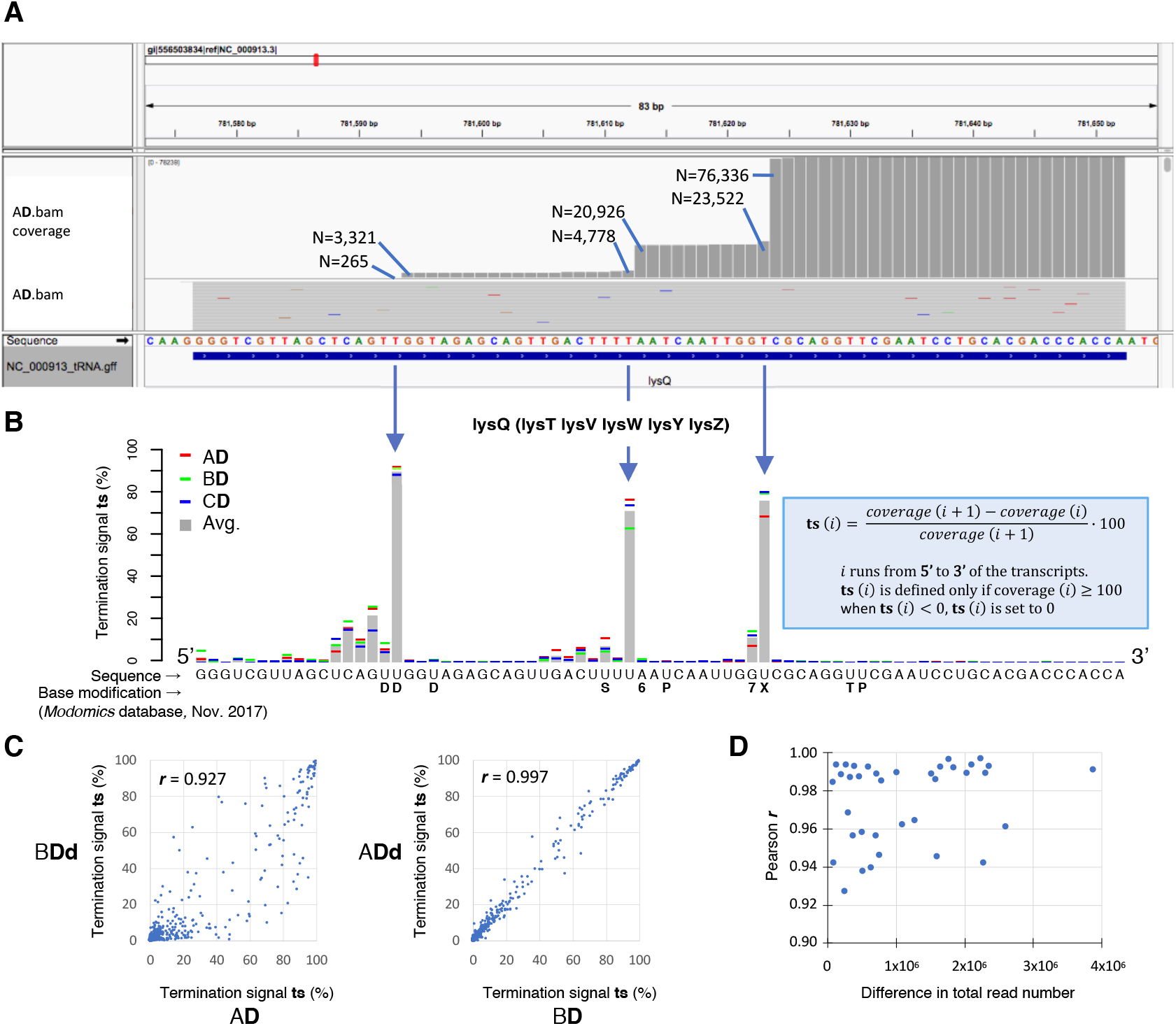
Overview of the analysis of deep-sequenced CCAЗ’-amplified RNA transcripts, a) Snapshot of an IGV window (Thorvaldsdóttir et al. 2013) centered on the lysQ tRNA gene of E. coli MG1655 genome, showing the coverage established from the AD sample. This gene is oriented 5’-3’ from left to right. Interruption of reverse transcription events (from 3’ to 5’) generate coverage jumps. b) Termination signal (ts, defined in the box) of lysQ tRNA, established from the coverage of AD, BD and CD samples. Because the lysQ sequence is identical to that of lysT, lysV, lysW, lysY and lysZ, reads are evenly distributed among these 6 genes. Base modifications symbols indicated below the sequence are from the Modomics database (Boccaletto et al. 2018) (Supplemental Table S1). c) Ts-ts Plots illustrating the lowest (left) and the highest (right) obtained Pearson correlation coefficients (r) between samples (see Table 3); n = 3146 in both plots. d) Pearson correlation coefficients as a function of the difference in total tRNA read number between samples.

## RESULTS

### Analytical treatment of tRNA transcripts coverages

Examination of tRNA coverages (Fig. 2a) revealed that they were highly specific and reproducible. The coverages decrease from the terminal CCA^3^’, with sudden jumps at locations of RT-impairing base modifications. Following analytical procedures introduced by Helm, Motorin and coworkers (Hauenschild et al. 2015) and Pan and coworkers (Zhang et al. 2015; Clark et al. 2016) these jumps were converted into termination signals (**ts**) highlighting the loss of coverage (in %) from nucleotide i+1 to nucleotide i (Fig. 2b). A **ts** signal is defined only when coverage (i) ≥ 100 reads. Furthermore, when negative values occur (reflecting very rare instances of amplified internal fragments), **ts** is set to zero. Figure 2a illustrates the analysis with the A**D** sample of *lysQ*, the coverage of which corresponds to the average expression of 6 identical genes (*lysQ, lysT, lysV, lysW, lysY, lysZ*). The average **ts** signal established from the A**D**, B**D** and C**D** samples (Fig. 2b) allows to highlight three major termination events associated with base modifications (see below for the analysis). Although the amplitude of these signals fluctuates among samples, they all correlate well (Fig. 2c and Table 3): Pearson correlation coefficients (**r**) are all above 0.92, with 22/36 coefficients higher than 0.98 (all associated *p* values are zero). Two significant observations can be made from Table 3. First, the amplitude of **ts** signals does not depend on RNA treatment: the two samples related by the lowest **r** (0.927) are A**D** and B**Dd** (Fig. 2c, left), while the highest **r** (0.997) is observed between the B**D** and A**Dd** samples (Fig. 2c, right). It suggests that the difference in **ts** signals in between samples is essentially related to PCR amplification. Remarkably, these signals are not affected by the total read number of a sample (Fig. 2d), which constitutes an unexpected robustness of the method. It can thus be concluded that **D** and **Dd** treatments do not have noticeable effects on the **ts** analysis, and, within the range of values obtained in the present study (Table 2), **ts** signals are not affected by the total number of reads provided by a deep sequencing experiment. Because these signals display fluctuations that can be higher than 90%, single deep-sequencing experiments cannot accurately determine their characteristic values (Fig. 3a), a quantitative aspect that has not been addressed in earlier analyses based on aborted cDNA signals (e.g. Hauenschild et al. 2015, Zhang et al. 2015). Since **ts** signals were found not to be affected by total RNA treatments, we established the average **ts** standard deviation by combining **-**, **D** and **Dd** samples in each replicate, and found an increase in the accuracy by a factor > 2 when three samples were combined (Fig. 3b; Table 4), the average standard deviation dropping from 6 to 3 % for **ts** signals equal or higher than ~7 % (i.e. **ts**_A+B+C_ > 20). The decrease of the average standard deviation with the number of samples being approximately logarithmic, our data suggest that 7 to 8 independent samples may provide signals ~10 times more accurate than single ones (supplemental Fig. S2). Furthermore, when samples are separately analyzed, from 11 to 23 tRNAs sequences lack full 5’ **ts** characterization due to low coverage at the 5’ end. By combining the coverages of the 9 independently sequenced samples of the present study (Table 2), only about 3% of the combined tRNA sequence length could not be characterized (Figure 4, supplemental File S4). Combining independent samples has, therefore, a double benefit: it significantly increases the precision in the value of **ts** signals, required to assess the level of modification of residues, and allows to establishing a complete (or nearly complete) **ts** analysis of all tRNA sequences.

**Figure 3.**
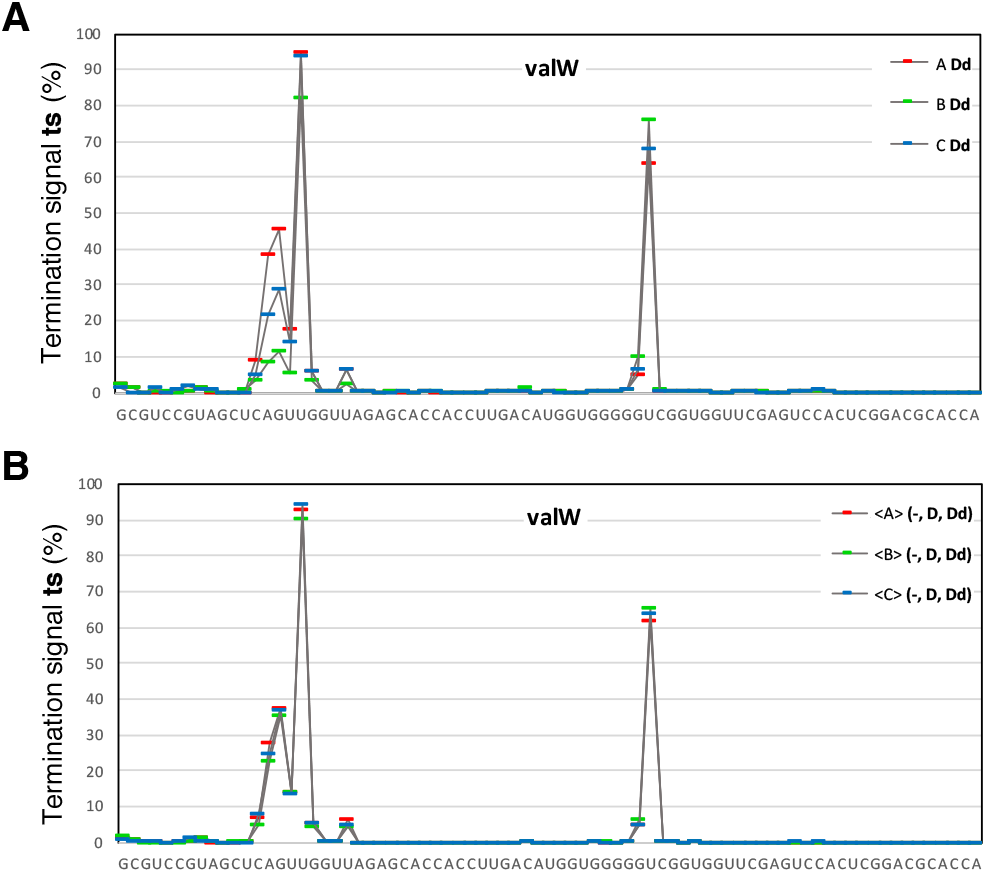
Gain in precision obtained in the ts analysis when ts signals from 3 samples are averaged, as compared to signals obtained from single samples (illustration with valW tRNA). a) Comparison of ts values between three biological replicates (A, B and C) with a single sample (Dd). b) Same comparison as in a, but with ts values of each biological replicate averaged over three samples (-, D, Dd). Because treatments of RNA samples (-, D, or Dd) were found not to noticeably alter ts signals (see Table 3 and text), they are considered as equivalent. For a global quantitative analysis, see Table 4.

**Figure 4.**
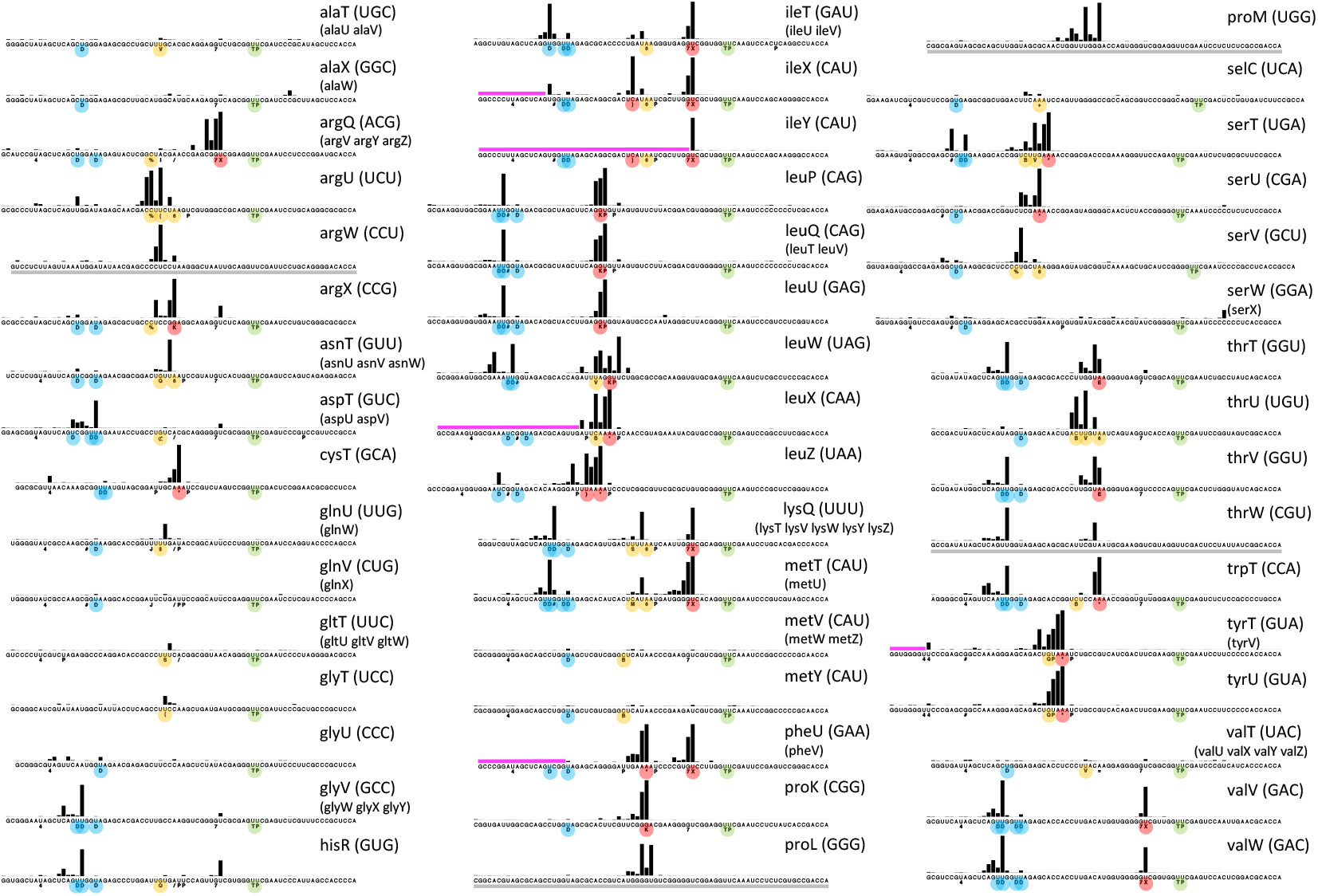
Atlas of ts signals of all E. coli tRNAs, with base modifications reported from the Modomics database (as of Nov. 2017). Modomics-uncharacterized tRNAs are underlined with gray lines. Identical tRNA genes are represented only once (copies are listed in brackets). Ts signals are established from the combined coverage of all 9 samples of the present work (Table 2). Magenta lines highlight 5’ portions of tRNAs for which the total coverage was too low to establish ts signals. Plots with ts values for all 9 samples are available in Supplemental File S4.

**Table 3:** Pearson correlation coefficients (**r**) between samples **ts** values. Correlation coefficients established from **ts** signals of tRNA, tmRNA and pseudo tRNA only when these signals are defined in all 9 samples (N = 3146 per sample); see Table 2 for the list of genes. Identical transcript copies (e.g. *metT, metU*) are included only once in the analysis. Dark green: 0.92 ≤ **r** < 0.95; medium green: 0.95 ≤ **r** < 0.98; light green: **r** ≥ 0.98. All associated *p*-values are 0.

|  | A- | B- | C- | AD | BD | CD | ADd | BDd | CDd |
| --- | --- | --- | --- | --- | --- | --- | --- | --- | --- |
| A- | 1.0000 | 0.9643 | 0.9584 | 0.9463 | 0.9565 | 0.9623 | 0.9611 | 0.9896 | 0.9686 |
| B- |  | 1.0000 | 0.9965 | 0.9892 | 0.9926 | 0.9930 | 0.9909 | 0.9424 | 0.9861 |
| C- |  |  | 1.0000 | 0.9937 | 0.9937 | 0.9926 | 0.9935 | 0.9377 | 0.9887 |
| AD |  |  |  | 1.0000 | 0.9928 | 0.9871 | 0.9923 | 0.9272 | 0.9876 |
| BD |  |  |  |  | 1.0000 | 0.9890 | 0.9968 | 0.9398 | 0.9844 |
| CD |  |  |  |  |  | 1.0000 | 0.9889 | 0.9422 | 0.9853 |
| ADd |  |  |  |  |  |  | 1.0000 | 0.9455 | 0.9892 |
| BDd |  |  |  |  |  |  |  | 1.0000 | 0.9566 |
| CDd |  |  |  |  |  |  |  |  | 1.0000 |

**Table 4:**
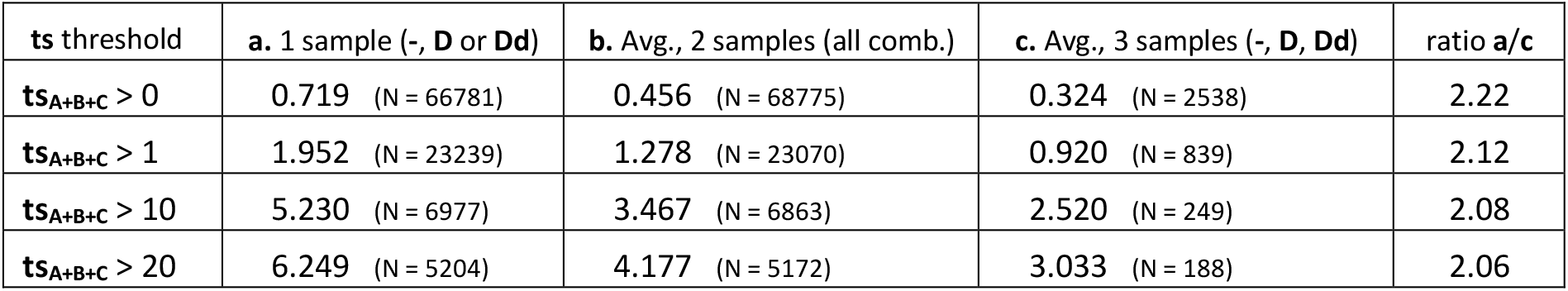
Comparison between biological replicates A, B and C: average **ts** standard deviation (%) as a function of the number of samples (**-**, **D** and/or **Dd**), established for four **ts** threshold. **ts_A+B+C_** > x implies that a particular standard deviation is computed only if the sum of **ts** signals of the three replicates at a given position is > x. About 2,500 **ts** values are simultaneously determined in all three replicates for a considered set of samples (the total number of nucleotide positions, including all tRNAs and three tRNA pseudogenes, is 3,916). Standard deviations are established for all possible combinations of samples. With one sample (**a**), there are 3^3^ = 27 combinations. With two samples (**b**), there are also 3^3^ = 27 combinations (ex: avg. (A**Dd**, A**-**); avg. (B**D**, B**-**); avg. (C**Dd**, C**D**)), whereas only one possibility occurs with three samples (**c**).

### Termination signal analysis of *E. coli* tRNA transcripts

The **ts** analysis generates signals similar to the arrest rate (Hauenschild et al. 2015) and modification index (Zhang et al. 2015; Clark et al. 2016). It provides a set of characteristic signals reflecting the modification state of a tRNA. Some modifications, such as **P** and **T**, can only be revealed by chemical treatments that make them interfering with RT enzymes (Motorin et al. 2007, Zhang et al. 2019). These were not investigated in the present study. To facilitate a rapid assessment of all tRNAs, four types of modifications are highlighted in Figure 4 (see supplemental Table S1 for one letter code). Green dots indicate the **T** and **P** modifications of the T-loop, for which no interference with the RT enzyme is noticeable. Blue dots highlight the **D** modification, with which strong **ts** signals are observed when three or more of such residues are present (see below). Orange dots signal modifications for which strong **ts** signals are observed only when two or more of such modifications are in close vicinity (**V**, **%**, **{**, **6**, **Q**, **c**, **\***, **$**, **S**, **B**, **M** and **+**) while red dots highlight modifications that are always interfering strongly with the RT enzyme (**X**, **K**, **\***, **}**, ) and **E**). Modifications that do essentially not affect the path of the RT enzyme (such as **4** and **#**) are left unhighlighted.

When modified residues are close to each other, non-linear, amplified **ts** responses are observed, a situation that can be attributed to the combined burden that these modifications impose to the core of the RT enzyme. It is thus often not possible to relate specific **ts** signals with a given modification, although the *pattern* of these signals is most often altered if a modification is missing. Also, when stretches of U residues occur on the 3’ side of modified bases, the position of **ts** signal(s) resulting from the interference with the RT enzyme is shifted to the 3’ side due to cDNA poly-A computational trimming (supplemental Figs. S1 and S3).

Following the path of the RT enzyme, **ts** signals are analyzed below from the 3’ end to 5’ end (Fig. 5a), crossing four regions highlighted in green (from the 3’ end to the 5’ side of the T-arm), salmon (variable loop), sand (anticodon stem-loop) and blue (from the D-loop to the 5’ end).

**Figure 5.**
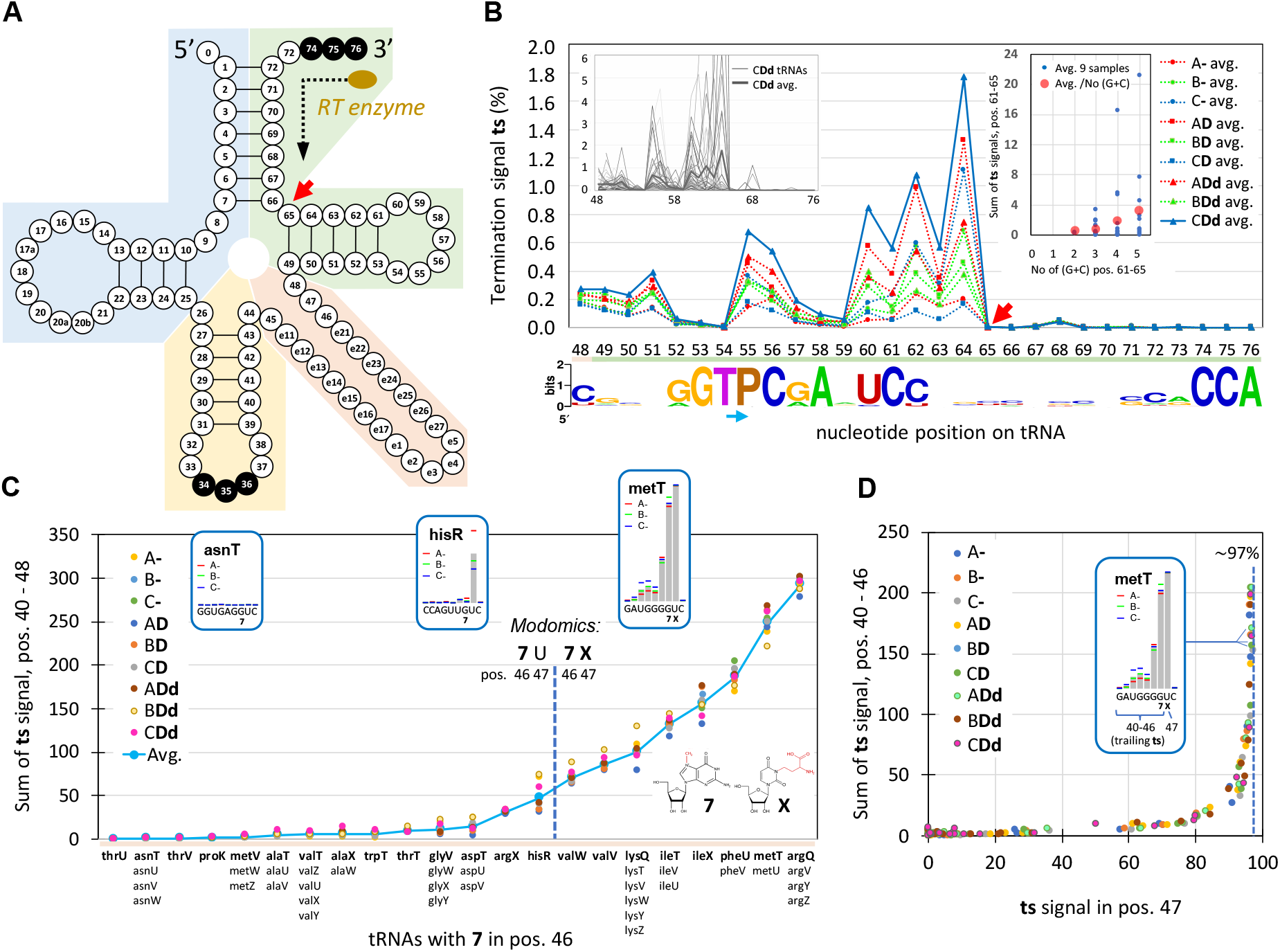
a) Standard tRNA nucleotide numbering according to the tRNA gene database curated by experts website (Abe et al., 2014). The colored regions correspond to the four successively analyzed tRNA segments, from 3’ to 5’: green, salmon, sand and blue. b) Ts analysis from pos. 76 to pos. 48 (green segment), established from an alignment of all tRNA transcripts. The sample with the highest values (CDd) is highlighted with a thick line. The left inset illustrates individual tRNA values (and the average) of the CDd sample. The right inset shows the sum of the ts signal as a function of the number of (G+C) residues in positions 61 to 65. Each blue dot corresponds to a tRNA; values averaged over all 9 samples (see Table 1). Red dots correspond to values averaged over all tRNAs. The base conservation logo reported at the bottom is based on all E. coli tRNA listed in Modomics. Because T and P modifications correspond to a stretch of two As on the cDNA, the ts signal of the two residues essentially combine into a single signal on the P residue (blue arrow) (see Material and Methods). c) Ts analysis of the variable loop (salmon segment). Sum of the ts signal from pos. 40 to 48 for all tRNA bearing the 7 modification in pos. 46, according to Modomics. For genes with multiple copies (e.g. asnT, asnU, asnV, asnW), only one set of values is indicated since the multiple mapping algorithm attributes a similar amount of reads to each gene. tRNA genes are ranked from left to right following an increasing average value of the ts signal. The vertical dashed line splits tRNAs into those with U (left) from those with X in pos. 47 according to Modomics (as of Nov. 2017). Insets illustrate three ts signals. d) Sum of ts values from pos. 40 to 46 (trailing ts signals) as a function of the ts value in pos. 47. Note that according to the Modomics database, all E. coli tRNAs with 7 and X modifications do not have any e residue in the variable loop, implying that the sequences analyzed in c and d follow standard numbering (panel a).

### Region of the T-arm

The length encompassing the 3’ end of the tRNA to the 5’ side of the T-arm being highly conserved (it comprises 28 nucleotides in *E. coli*, tRNA^sel^ being the only exception with 27 nts), we performed the **ts** analysis at once by combining signals of all tRNAs (Fig. 5b). Only **T** and **P** modifications are present, which do not constitute an obstacle to the RT enzyme (Motorin *et al.* 2007). From the 3’ end, the first region presenting a significant (albeit very low) **ts** signal starts after the entry of the T-arm (pos. 65; red arrow). It appears to be related to the RT enzyme encountering the densely folded region of the tRNA, as suggested by a correlation between the proportion of (G+C) residues in the T-arm and the summed **ts** signals along this segment (Fig. 5b, right inset). The line chart of Figure 5b reveals that the profile of the average signal is highly reproducible: essentially only the amplitude varies among the 9 samples, which may be related to PCR amplification. The very low value (note the scale of the y axis) and reproducibility of these signals highlight a very low level of noise, which is a hallmark of our protocol. The **ts** signals of pos. 54 and 55 (**T** and **P**) combine on pos. 55 (blue arrow on Fig. 5b) as a result of poly-A computational trimming (supplemental Figure S3). The variability of the **ts** signal observed at positions **T** and **P** for individual tRNAs (Fig. 5b, left inset) reveals that the average signal is not due to the modifications *per se*, but rather to the local structural context.

### Region of the variable loop

Only two modified bases are present at the start of the variable loop in some *E. coli* tRNA: 3-(3-amino-3-carboxypropyl) uridine (**X**) and 7-methylguanosine (**7**). **X** is analog to N1-Methyl-3-(3-amino-3-carboxypropyl)-pseudouridine, known to allow only minimal bypass by RT enzymes, while **7** was demonstrated not to detectably affect reverse transcription (Motorin *et al.* 2007). According to the *Modomics* database, the **7** modification is present in position 46 in slightly more than 50% of all *E. coli* tRNAs, and only when **7** is present does **X** occur in position 47 for half of these tRNAs. Our data show that a strong **ts** signal is always present when **X** is listed in *Modomics* (Fig. 5c). Furthermore, additional “trailing” **ts** signals occur on the 5’ side of the major **ts** signal when it is higher than ~80% (Fig. 5d). Two important deductions can be made from our results: Figure 5c shows that the **ts** signal associated with **X** cannot unambiguously separate **7**U from **7X**-modified tRNAs, suggesting that all tRNAs with **7**U in positions 46-47 are converted to **7X** to some extent although they may not be listed as such in *Modomics* (prominent cases are argX and hisR). This observation corroborates biochemical analyses performed by Meyer et al. (2019), who showed that the modifications of these two residues are interdependent. Furthermore, the value of the **ts** signal in position 47 converges towards ~97% when the overall signal reaches the highest values (Fig. 5d). Assuming that the fraction of **X**-modified tRNAs reaches 100% in that case implies a bypass rate of this modification by the RT enzyme of about 3%. A straightforward interpretation of trailing **ts** signals is that RT enzymes bypassing the **X** modification drop off one or a few nucleotides downstream as a result of the perturbation caused by the modification embedded in the chimeric cDNA/RNA duplex.

### Region of the anticodon loop

The anticodon loop is by far the region with the most diverse types of modifications (Boccaletto et al. 2018). Figure 6 and supplemental Figure S4 highlight **ts** signals generated by some of the most frequent *E. coli* modifications.

**Figure 6.**
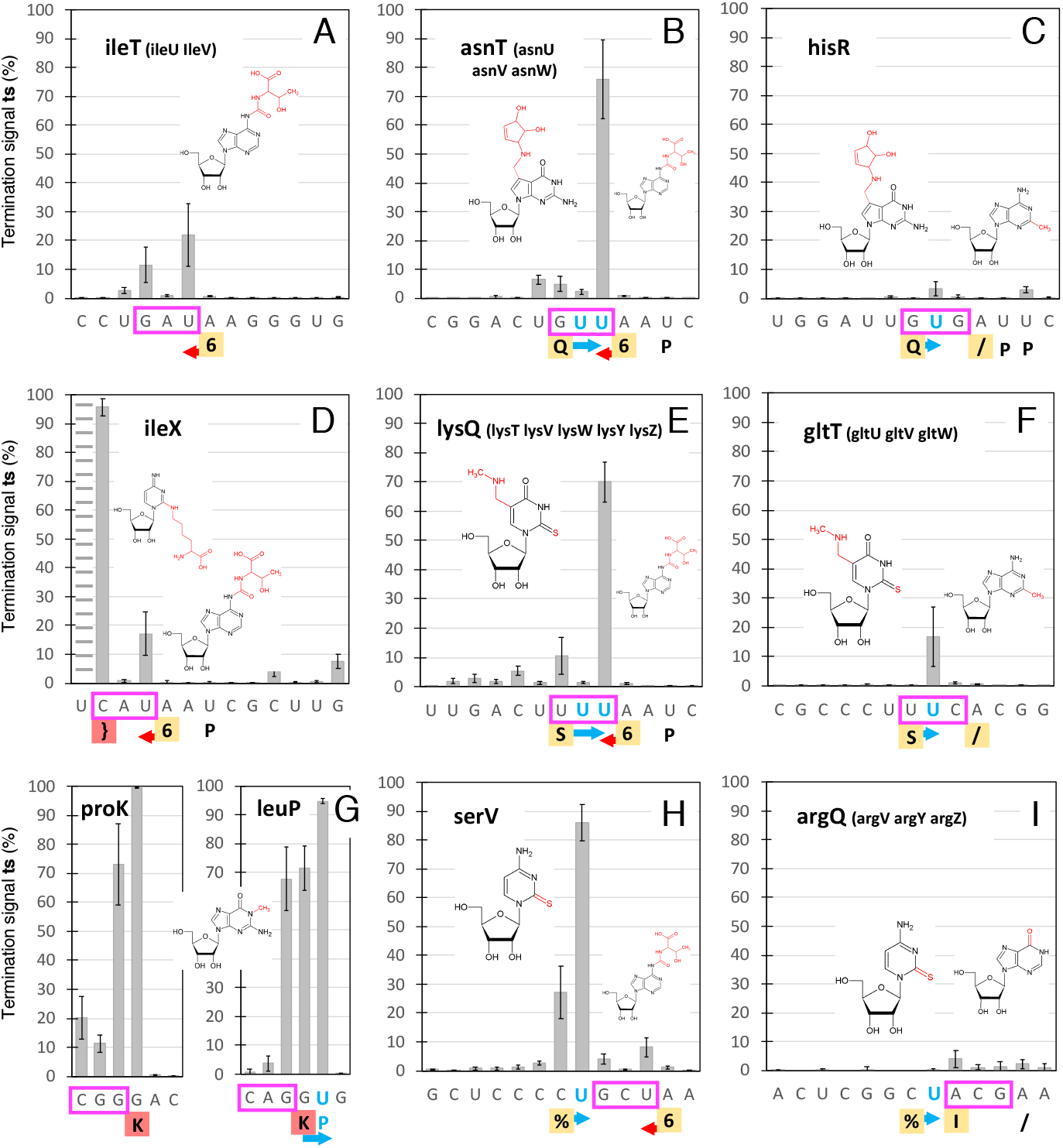
Ts analysis of the anticodon loop. Plots of the average values and standard deviations established from all 9 samples (Table 1), highlighting ts signatures of some typical E. coli combinations of modifications (panels a to i). Anticodons are boxed in magenta. Known modifications are reported from the Modomics database below the sequences (5’-3’). Highlighted modifications (in yellow and red) are illustrated in the plots following the same order (cartoons are from the Modomics website). Blue arrows indicate shifts of ts signals associated with a modification that result from the presence of U residue(s) on its 3’ side (highlighted in blue), an artifact caused by poly-A computational trimming (see Material and Methods). Red arrows highlight the shift of the ts signal to the 5’ direction observed with the 6 modification. The ts signal could not be established on the 5’ side of the } modification for IleX (panel d, dashed lines) in all samples. Although Inosine (I) is associated with a low ts signal (panel i), an examination of the read coverage shows that it leads to incorporation of 86%C and 14%U on the cDNA, a typical signal of the presence of this modification (Schwartz and Motorin, 2017).

#### N6-threonylcarbamoyladenosine (6)

This large modified base appears to constitute a moderate obstacle to the RT enzyme (Fig. 6a, b, d, e, h). Unexpectedly, a major **ts** signal is not present at the location of the modification, but one base downstream the path of the RT enzyme (small red arrow), suggesting that it generates an impediment only after being incorporated into the cDNA/RNA duplex. An examination of the read processing revealed that the observed **ts** pattern (Fig. 6a) is partly the result of incomplete 5’ poly-A trimming due to the incorporation of other bases (often a C) in the poly-A, followed by local mapping (supplemental Fig. S3). Thus, the pattern of **ts** signals associated with a modification may not always be interpreted as a direct consequence of the interaction between the modification and the RT enzyme. *Interference of **6** with other modifications:* **6**, that is only found in pos. 37, is often accompanied by other modifications in pos. 34 (**Q**, **S**) or in pos. 32 (**%**). Figure 6b, 6e and 6h show that whenever this occurs, the combined perturbations generate a **ts** response that is stronger than a linear combination of individual **ts** signals (see below).

#### Queuosine (Q)

The effect of this large modification on RT processivity was not assessed in the seminal work of Motorin et al. (2007). Surprisingly, it does not generate a strong **ts** signal (Fig. 6c), suggesting that the RT enzyme can easily cross this residue if it is not in the vicinity of other modifications (Fig. 6b).

#### 5-methylaminomethyl-2-thiouridine (S)

This modification generates a small **ts** signal (Fig. 6f) similar to **Q** (Fig. 6c). When combined with **6**, the resulting **ts** signal (Fig. 6e) is similar to that of the **6**-**Q** combination (Fig. 6b).

#### 2-thiocytidine (%)

The nearly isosteric C2-O → C2-S modification of C does not generate any **ts** signal when it is in an unperturbed environment such as the one shown in Figure 6i. In that case, the closest modified neighbor is an inosine (**I**), which is known not to interfere with the RT enzyme, although it leads to the incorporation of C and U (Motorin et al, 2007). However, in the context of a **6** modification in pos. 37 (Fig. 6h), **%** generates an unexpectedly high **ts** signal, suggesting that additional modification(s) unlisted in *Modomics* could be present on the **%** residue in that case.

#### 2-methyladenosine (/)

This modification was found to only generate RT pause at low dNTP concentration (Motorin et al. 2007). Representative examples (Fig. 6c, f, i) show that it is indeed associated with almost no **ts** signal. ***2-lysidine (}).*** Only one *E. coli* tRNA bears this large modification, that is associated with a strong **ts** signal (Fig. 6d).

#### 1-methylguanosine (K)

This small modification is known to constitute a major obstacle for the RT enzyme (Motorin et al. 2007, Zheng et al. 2015). This can clearly be seen in our data: with *proK* tRNA (Fig. 6g, left), the average **ts** signal at the position of the modification is as high as 99.65 ± 0.08 %, corresponding to a coverage drop from 283,925 to 864 reads with the A**D** sample. Assuming a rate of modification of 100% in that case implies a bypass rate of the order of 0.35%. As a result of computational trimming, the major **ts** signal is shifted when a pseudouridine (**P**) is present on its 3’ side (Fig. 6g, right).

#### 2-methylthio-N6-isopentenyladenosine (*)

This base has a complex pattern of modifications that is expected to strongly impair the RT enzyme. It consistently generates a high **ts** signal (supplemental Fig. S4a). All related tRNAs only have **7**, **T** and **P** modifications upstream the path of the RT enzyme, which do not affect its processivity. The only exception is *pheU (pheV)*, that bears the dual **7X** modifications in the variable loop. Although this early roadblock generates a strong **ts** signal in these tRNAs, our data show that the **ts** signal at the **\*** position is identical to that of other tRNAs (supplemental Fig. S4b), demonstrating that **ts** signals generated by distant modifications along transcripts may be considered as independent.

#### Uridine 5-oxyacetic acid (V)

In a way similar to the % modification, **V** essentially does not generate any **ts** signal when occurring in a sequence context without perturbing modifications (supplemental Fig. S4c, d). When other nearby modifications are present, such as in the case of *LeuW*, the effect of this base is difficult to assess because stretches of U (supplemental Fig. S4e).

#### 2’-O-methylcytidine (B)

This modification is known to generate RT pauses at low dNTP concentration (Motorin et al. 2007). In a sequence context without any other modified base, **B** does only generate a faint **ts** signal similar to **V** (supplemental Fig. S4f). With a **\*** modification 6 bases upstream the path of the RT enzyme, the associated **ts** signal is slightly more pronounced (supplemental Fig. S4g), and with an additional **V** residue in between **B** and **\***, a strong and complex **ts** pattern is observed (supplemental Fig. S4h). This last example illustrates the non-linear response of the **ts** signal in the presence of three modifications, two of which do almost not generate any signal when found isolated (supplemental Fig. S4c, f).

### Region of the D-loop

#### Dihydrouridine (D)

With 152 reported instances, **D** is the most frequent tRNA modification found in *E. coli MG1655* tRNAs according to the *Modomics* database. Because the D-loop is located near the 5’ end, there is often not much exploitable signal in each sample due to 3’ upstream termination events caused by RT-blocking modifications (Fig. 2a). However, with the coverage provided by 9 deep-sequencing samples, accurate **ts** signals can be established up to the 5’ end for all but 5 weakly expressed tRNAs (Fig. 4). **D** is known to only create RT pauses (Motorin *et al.* 2007). The presence of major **ts** signals associated with this modification therefore came as a surprise. It turns out that these signals are highly dependent on the number of dihydrouridine(s) known to occur on the D-loop, a phenomenon already noticed by Clark et al. (2016), although our quantitative analysis leads to a different conclusion with regard to the effect. Considering the sum of **ts** signals in 11-nt windows centered on each dihydrouridine, a near clear-cut threshold separates loops with one and two from those with three and four dihydrouridines when the window with highest **ts** sum is considered (Fig. 7a,b). No clear sequence context allows to predict the position of the major **ts** signal, which can be associated either with a doublet of dihydrouridines *(glyV)* or with one dihydrouridine *(ileT).* Furthermore, it can occur ‘early’ *(aspT)* or ‘late’ *(ileT)* along the path of the RT enzyme (Fig. 4). Clark et al. (2016) found a strong signal associated with a doublet of dihydrouridine in human tRNA^asn^ (GUU), and concluded that a doublet sufficiently destabilizes the RT enzyme to lead to transcription abortion. Our analysis suggests instead that all three U residues present on the D-loop of this tRNA are likely modified. Similarly, our analysis predicts the presence of an unlisted **D** modification on the D-loop of *leuW*, only known to harbor two dihydrouridines: the value of the **ts** signal (Fig. 7b) and the sequence context around an unmodified U (Fig. 7d), which is identical to five others fully modified leucine tRNAs, are both consistent with the presence of a third dihydrouridine. With a high **ts** signal and only two possible **D** modifications on the loop, the case of *serT* is less clear (Fig. 7d). Examination of the *Modomics* database reveals that whenever a uridine is present in the D-loop, it is essentially modified into **D**, with two exceptions, *glyT* and *argU* (Fig. 7c). Our predictive analysis (Fig. 7b) suggests that at least two among the four uridines present in the D-loop of *glyT* are indeed not modified (Fig. 7d), supporting the possibility of this exception (the **ts** signal associated with *argU* cannot be unambiguously interpreted).

**Figure 7.**
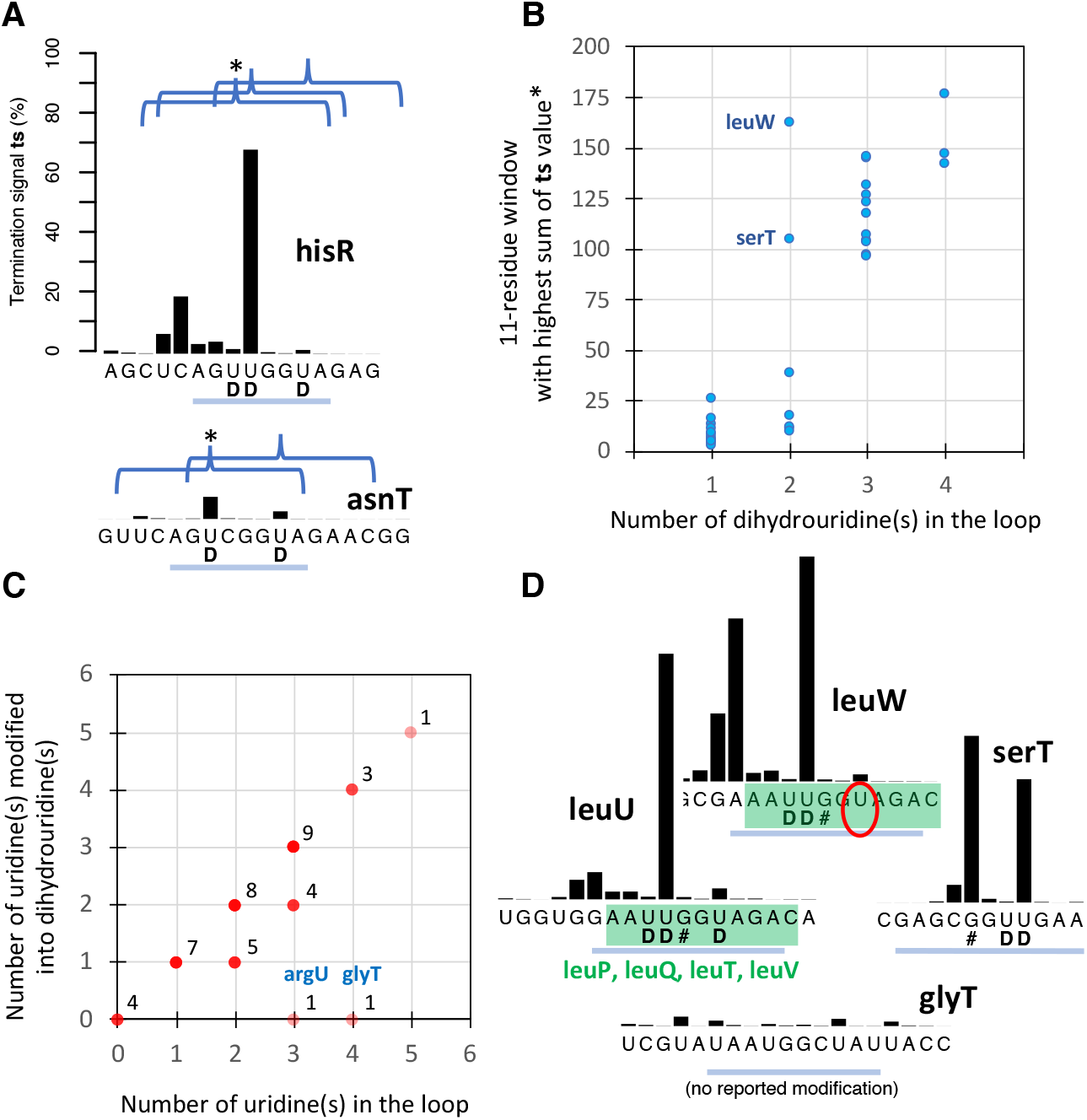
Ts analysis of the D-loop. a) IIlustrati on of the effect of the presenee of >= 3 (hisR, on top) and < 3 (asnT, bottom) dihydrouridine modifications. Curly brackets encompassing a window of 11 nts are centered on every reported dihydrouridine modification. b) Plot of the sum of ts signals in the window with the highest signal (*) as a function of the number of reported dihydrouridine modifications. c) Plots of the number of uridine (s) modified into dihydrouridine(s) in the D-loop according to the Modomics database (nov. 2017). d) Left and top: ts signals and sequence similarity between leuU, leuP, leuQ, leuT and leuV (in green, left), for which three dihydrouridines have been reported, and leuW (top), for which ts analysis (in b) and sequence similarity suggest the presence of an unlisted dihydrouridine (circled). Bottom: The low ts signal of glyT is consistent with the absence of at least two dihydrouridine modifications among the four uridines of the loop. Right: serT displays high ts signals although only two dihydrouridines are present in the loop. Ts values are established from the combined coverage of all 9 samples (Table 1). D-loop segments are highlighted with a blue line on panels a and d.

### Quantification of -CCA ending tRNA transcripts

Since amplified 3’ tRNA segments are always long enough to be uniquely mapped onto the genome (except for identical tRNA genes), a relative frequency of tRNA isoacceptors can be established by reporting the number of reads at the position of the 3’ end (Fig. 8a; see also Fig. 2a). It is, however, important to keep in mind the non-linear nature of PCR amplification, that changes RNA transcripts true relative frequencies. Figure 8a shows that the observed frequencies are comparable among all three biological replicates, and that sample treatment (**–**, **D** or **Dd**) only has a marginal effect, except for isoacceptors for which the amino acyl linkage is particularly resistant (ile, val) or prone to hydrolysis (ser) (Matthaei *et al.* 1966). In the first case, deacylation (**d**) increases these frequencies, while isoacceptors that are already substantially deacylated without any treatment become proportionally less represented (ser). Our data thus show that deacylation still occurs with (**–**) and (**D**) samples, that allowed the ligation of the RNA adapter. The mild pH of the RNA adapter ligation step (pH 7.5; overnight incubation at 10°C) is likely responsible for this phenomenon. Deacylated samples (**Dd**) provide a picture of the population of all tRNA isoacceptors regardless of their aminoacylation state. Figure 8b shows that their relative frequency mainly follows that of their corresponding codons in the *E. coli* genome, a phenomenon that is expected based on tRNA gene copy number (Fig. 8c) (Higgs and Ran 2009; Du *et al.* 2017). Transcripts of a group of isoacceptors (gln, asp, ser, glt, val gly and ala), however, appear significantly underrepresented, a feature that was not observed in a study in which quantification was achieved by probe hybridization on gel (Dong *et al.* 1996; Supplemental Fig. S5). While we could not find a clear explanation for this result, it is possibly related to the -CCA 3’ end requirement of our protocol. The 3’ terminal adenosine can be specifically removed by RNase T in *E. coli* (Deutscher *et al.* 1984, 1985), in which case these tRNAs would not be amplified. Results of conventional deep sequencing indeed show that a significant proportion of tRNA 3’ segments lack the terminal adenosine (unpublished results), suggesting that the removal of this residue, that can be re-added by the CCA-adding enzyme, may play a role in translation regulation. A tRNA quality control mechanism involving these two enzymes has also been suggested (Wellner *et al.* 2018). The reason for which the relative frequency of these seven CCA-ending isoacceptors is unexpectedly low might be related to the cellular concentration of their cognate amino acid (Fig. 8b): in *E. coli* cells grown in conditions similar to that of our cultures, five of these amino acids were found to be at least one order of magnitude more abundant than the other ones (Bennett *et al.* 2009), while the amino acid of the most underrepresented isoacceptor, glt, is even 10^3^ times more abundant in normal growth conditions. High substrate concentration enables a fast turnover of tRNA aminoacylation, which might abrogate the necessity for a large amount of functional isoacceptors. Further experiments are required to clarify this issue.

**Figure 8.**
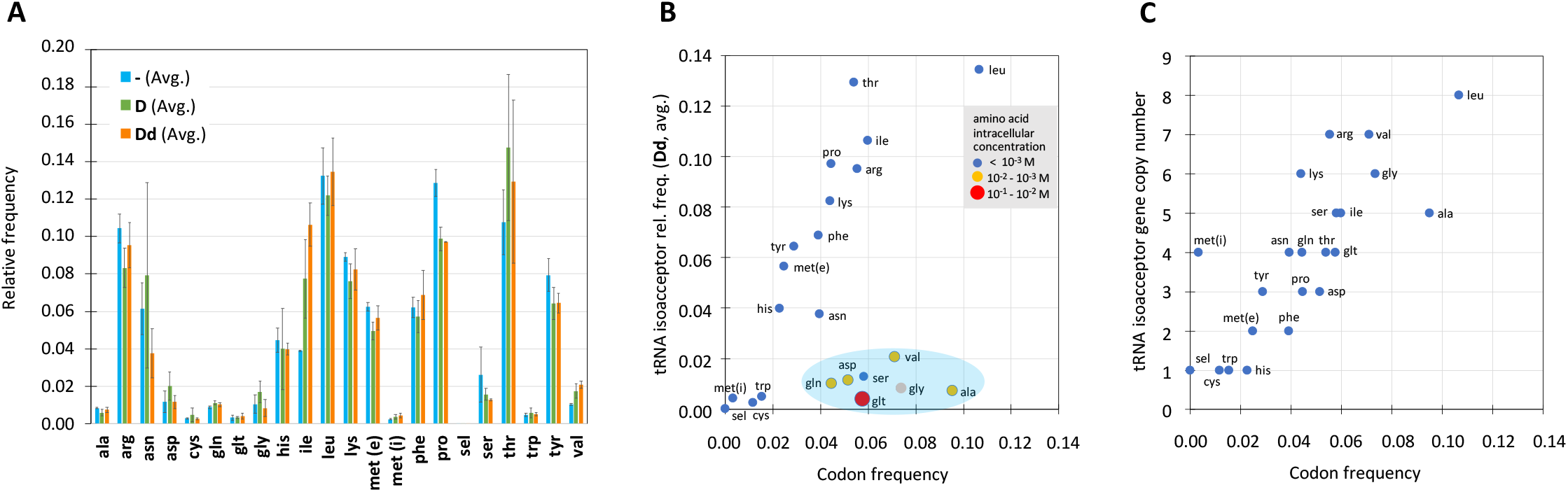
Relative quantification of E. coli CCA-ending tRNA isoacceptors (cultures grown in LB medium, sampled at OD650nm 0.4) and codon usage. a) Relative quantification of tRNA isoacceptors as established from the coverages at the 3’ position of each tRNA (see Fig. 2a; values are in Supplemental File S6). Each bar represents the average relative frequency calculated over three replicates. Avg (-): A-(N = 6,299,382), B-(N= 5,026,263), C-(N = 6,793,490). Avg (D): AD (N = 7,056,586), BD (N = 6,669,159), CD (N = 7,385,017). Avg (Dd): ADd (N = 8,899,275), BDd (N= 7,301,761), CDd (N = 6,598,669). b) Relative frequency of CCA-ending tRNA isoacceptors as a function of codon frequency in the E. coli K12 genome*. Inset: intracellular concentrations of encoded amino acids in E. coli determined by Bennett et al. (2009). Cells grown under normal conditions, with glucose as the major source of carbon; cultures sampled at OD650nm = 0.35. Glycine is the only encoded amino acid for which concentration was not determined in this study. c) E. coli MG1655 tRNA gene copy number as a function of codon frequency*. *data from https://openwetware.org/wiki/Escherichia_coli/Codon_usage

## DISCUSSION

Compared to recent tRNA sequencing proposals (Cozen et al. 2015; Zeng et al. 2015, Hauenschild et al. 2015; Schwartz and Motorin 2017; Gogakos et al. 2017), the present method is relatively easier to implement because no total RNA pre-processing (such as demethylase treatment and/or RNA fragmentation) is required, and we found that a conventional RT enzyme (Superscript III) is processive enough to generate signals far to the 5’ end of the tRNAs. Furthermore, avoiding fragmentation enables an accurate gene mapping of all reads, as they all start at the 3’ end of the tRNA, where essential identity elements are present. Two methodological aspects ensure the lowest possible bias: first, the ligation of the 3’ adapter is non-selective owning to the invariant CCA^3^’ dangling end of mature tRNAs, that are specifically amplified by GGT-ending primer during a first round of PCR. This feature of our protocol also allows to virtually remove all background noise, which is present in current protocols (Hauenschild et al. 2015, Clark et al. 2016, Gogakos et al. 2017). Second, TdT polyadenylation of cDNA after reverse transcription circumvents the issue of inefficient 5’ tRNA adapter ligation in conventional deep-sequencing protocols. Circularization of cDNA, which was used by Zheng et al. (2015) in their own protocol, is another possibility. The obtained termination signals (**ts**) provide an atlas of the modified state of all tRNAs (Fig. 4). Furthermore, we showed that two limitations inherent to deep-sequencing experiments and the use of a conventional RT enzyme are simultaneously resolved by combining several samples: a significant gain in the accuracy of **ts** signals is achieved, and these signals can be established down to the 3’ end of all tRNAs. The combined tRNA read coverage of the 9 samples reported in the present study, totalizing about 62 million mapped R2 reads, were just short of generating a full tRNA **ts** atlas: about 3% of sequence length was left uncharacterized (magenta lines in Fig. 4). However, current standards of *Illumina* sequencing experiments generate 200 million reads. Based on the results of our computational processing, in which about 60% of the initial reads were mapped, full tRNA characterization is almost certainly achievable with a single *Illumina* sequencing experiment. While some modifications do not interfere with the RT enzyme, RNA chemical treatment may alter these silent modifications and make them visible with our protocol (Motorin et al. 2007), a procedure that has recently been applied by Zhang et al. (2019) with an alternate protocol. Several silent modifications can still be detected when they occur nearby each other (highlighted in orange in Fig. 4), implying that a change of the **ts** pattern is observed if one modification is missing. Thus, while the present method may not be used as a primary tool for the identification of modified bases, its power is to provide a robust and extensive diagnostic of the modified state and relative frequencies of all cellular tRNAs with minimal benchwork.

## Acknowledgements

We thank Jinwei Zhang, Sine Lo Svenningsen and Michael A. Sorensen for helpful comments and suggestions on several aspects of our analysis, and Philippe Bouloc for providing lab space and a stimulating working environment. Ji Wang was supported by the Chinese Scholarship Council, grant No. 201206140111.

## Author contributions

JL and DG conceived of the project. JW and FL performed the experimental work. JL, DG, JW, and CT interpreted the results and performed the analysis. JL wrote the manuscript, with inputs from DG, CT and JW. CT and DG implemented the computational pipeline. All authors read and approved the manuscript.

