## Supplemental Figures and Tables for "Highly reproducible characterization of *Escherichia coli* tRNA epitranscriptome with a simple method of library preparation for deep sequencing"

### Supplemental material

**Supplemental Figure S1** : Computational processing of deep-sequencing *Illumina* experiments

**Supplemental Figure S2** : Effect of the number of samples on the gain in precision of **ts** signals

**Supplemental Figure S3** : Illustration of how **ts** signals are generated through the processing of initial cDNA reads

**Supplemental Figure S4** : **Ts** analysis of base modifications on the anticodon region (complement to Figure 6)

**Supplemental Figure S5** : *E. coli* K-12 MG1655 tRNA isoacceptors relative frequency in cells

**Supplemental Table S1** : Modified bases occurring on *E. coli* K-12 MG1655 tRNAs (*Modomics* database notation)

Additional **Supplemental Files** are separately provided:

**Supplemental File S1** : Total RNA library preparation for deep sequencing: full protocol (pdf format)

**Supplemental File S2** : bm.gff. This file specifies the positions of all modified bases reported in *Modomics* along the *Escherichia coli* str. K-12 (substr. MG1655) genome (NC_000913.3)

**Supplemental File S3** : tRNA_features.txt. This file specifies the genomic coordinates of all *Escherichia coli* str. K-12 (substr. MG1655) tRNAs, three pseudo tRNAs and one tmRNA

**Supplemental File S4** : **Ts** signal sequence analysis of all *E. coli* tRNAs and three *E. coli* pseudo tRNAs: graphs (pdf format)

**Supplemental File S5** : **Ts** signals sequence analysis of all *E. coli* tRNAs and three *E. coli* pseudo tRNAs: table (Excel format)

**Supplemental File S6** : 3' Coverage of all *E. coli* tRNA and *pauD*, *pawZ* and *ptwF* tRNA pseudogenes (Excel format)

**The computational analysis pipeline is available at** https://github.com/i2bc/dts-seq

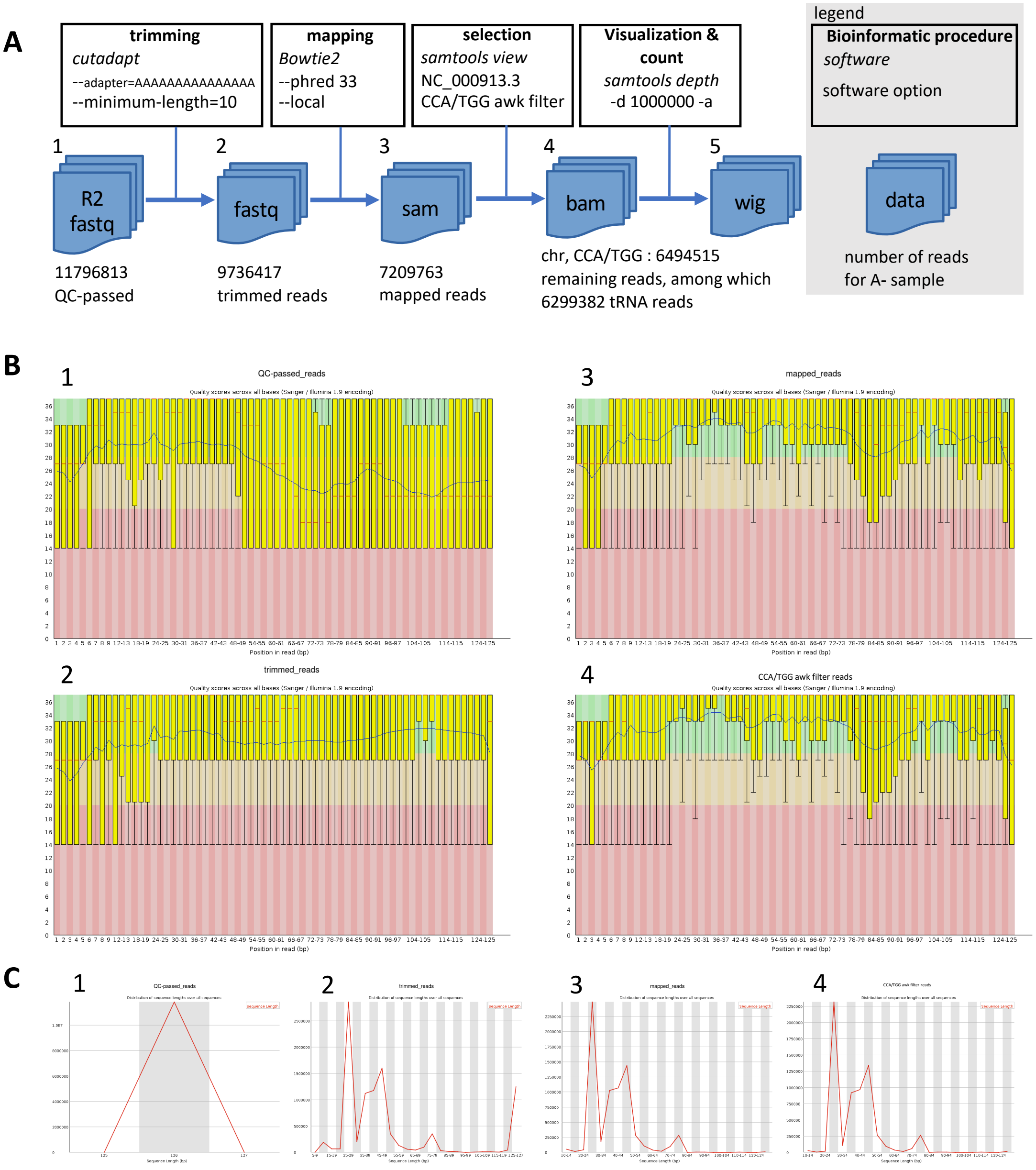


**D**

| | | samples | | | | | | | | |
|---|---|---|---|---|---|---|---|---|---|---|
| *Processing step* | | **A-** | **B-** | **C-** | **AD** | **BD** | **CD** | **ADd** | **BDd** | **CDd** |
| **1** | QC-passed | 11796813 | 13114509 | 11455875 | 11680073 | 13517533 | 10775965 | 15388902 | 12562074 | 10691188 |
| **2** | trimmed | 9736417 | 8639091 | 9522279 | 10257977 | 10586494 | 10181006 | 12595828 | 10640726 | 9146981 |
| **3** | mapped | 7209763 | 5765983 | 7528570 | 7829777 | 7574852 | 8282069 | 9771220 | 8149220 | 7276564 |
| **4** | CCA/TGG f. | 6494515 | 5150196 | 6941871 | 7229758 | 6803833 | 7551225 | 9045057 | 7457325 | 6771946 |

**Supplemental Figure S1**

Computational processing of deep-sequencing *Illumina* results (*Illumina* sequencing achieved by *Genewiz*). Shown values (**A**) and qualitative/quantitative analyses (**B, C**) are from the A- sample; all 9 samples (see panel **D**) provided similar results.

The results of each bioinformatic processing step (1 to 4) shown in **A** are analyzed in panels **B** and **C**, while panel **D** shows the number of sequences remaining at each step.

Computational workflow (**A**): among half a dozen tested strategies, the established workflow provided the sharpest termination signals. From the initial ***QC-passed reads*** (**1**), ***trimmed reads*** (**2**) were obtained with *cutadapt* by identifying and trimming all TdT-added poly-A (i.e. poly-Ts on the tRNA gene sequence) with at least 15 nt. Reads shorter than 10 nt were removed since they could not be unambiguously mapped onto the genome. See https://doi.org/10.14806/ej.17.1.200

In a second step, a local mapping of the reads onto the *Escherichia coli* genome* with *bowtie2 (bowtie2-align-s version 2.2.8, options –phred 33 –local)* was achieved, providing the ***mapped reads*** **(3)**. See https://www.nature.com/articles/nmeth.1923

The *–local* option maximizes the stretch of matching bases by removing mismatches at the end of the reads (softclipping).

**str. K-12 substr. MG1655 (GCF_000005845.2, ASM584v2).*

*Supplemental Figure S1 : end of legend*

In a third step, mapped reads were filtered with a custom *awk* script to select only the reads ending with *CCA/TGG (depending to the orientation of the tRNA on the genomic sequence),* thus complementing the selection of mature tRNA achieved during library preparation. This resulted in the ***final mapping*** **(4)**, from which the ***wig*** file **(5)** compiling the coverage of all expressed RNA genes was established with *samtools depth* using options *–d* 1000000 and *–a*. *–d* 1000000 specifies the maximal coverage (in read number) at one position, while *–a* (all) specifies that the coverage at all positions are needed, including positions with no read.

-----------------------------------------------------------------------------------------------------------------------------------------------------

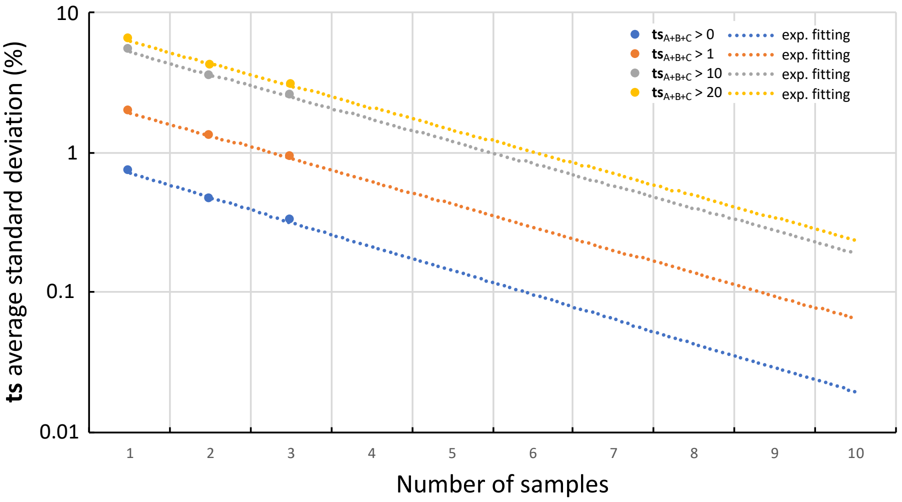


**Supplemental Figure S2**

Effect of the number of samples on the gain in precision of **ts** signals in the comparison between three biological replicates (A, B and C): **ts** average standard deviation (%) established for four different **ts** threshold. **ts**$_{\mathbf{A+B+C}}$ > x implies that a particular standard deviation is included in the computation only if the sum of **ts** signals over the three replicates at a given position is > x. Values were computed for 1, 2 and 3 samples (see Table 4 for details); exponential fitting is shown for up to 10 values.

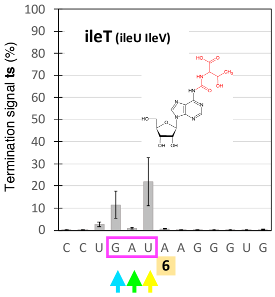


| | |
|---|---|
| >SL-HBW:622:CA37GANXX:1:1114:19391:48157 | **Read ID** |
| TTTTTTATTTTTTCTGCTACCTCCTTTTTTTTCTTTTTTTTTTTTTTTTTTTTTTTTGTTCCGCGGTTTTTTTTTTTTTTTTTTTAAGGGTGAGGTCGGTGGTTCAAGTCCACTCAGGCCTACCA | **Initial QC-passed read** |
| AAGGGTGAGGTCGGTGGTTCAAGTCCACTCAGGCCTACCA | **trimmed read** |
| AAGGGTGAGGTCGGTGGTTCAAGTCCACTCAGGCCTACCA | **–*local* alignment : removal of end mismatche(s)** |
| >SL-HBW:622:CA37GANXX:1:1306:11690:31837 | **Read ID** |
| TTTTATGTTACGGCGGTCGCCTGTTTTTTTTTTTTTTTGTTTTTTTCAGTTCGACGAGCGGGAATTTTTTTTTTTTTTTTTTTTTAAGGGTGAGGTCGGTGGTTCAAGTCCACTCAGGCCTACCA | **Initial QC-passed read** |
| AAGGGTGAGGTCGGTGGTTCAAGTCCACTCAGGCCTACCA | **trimmed read** |
| AAGGGTGAGGTCGGTGGTTCAAGTCCACTCAGGCCTACCA | **–*local* alignment : removal of end mismatche(s)** |
| >SL-HBW:622:CA37GANXX:1:2106:9948:49997 | **Read ID** |
| TTTTTTTTTTATGATTTTCCCCTGGATTTTTGTACTTTTTTGTTTTTTAGTTCCGACATTCTTTCGGTTTTTTTTTTTTTTTTTTTAAGGGTGAGGTCGGTGGTTCAAGTCCACTCAGGCCTACCA | **Initial QC-passed read** |
| AAGGGTGAGGTCGGTGGTTCAAGTCCACTCAGGCCTACCA | **trimmed read** |
| AAGGGTGAGGTCGGTGGTTCAAGTCCACTCAGGCCTACCA | **–*local* alignment : removal of end mismatche(s)** |
| >SL-HBW:622:CA37GANXX:1:2206:2834:40657 | **Read ID** |
| TTTTTTTTTATATGTTCACTATCTATTTTTATTCTTTTATGTTTTTACTGTCTAACCATCAAATTTTTTTTTTTTTTTTTTTTTTTAAGGGTGAGGTCGGTGGTTCAAGTCCACTCAGGCCTACCA | **Initial QC-passed read** |
| AAGGGTGAGGTCGGTGGTTCAAGTCCACTCAGGCCTACCA | **trimmed read** |
| AAGGGTGAGGTCGGTGGTTCAAGTCCACTCAGGCCTACCA | **–*local* alignment : removal of end mismatche(s)** |
| >SL-HBW:622:CA37GANXX:1:2315:16187:13372 | **Read ID** |
| TTTTATTTTTTGTCTTCCTCTGTGTTTTTTTTTTTTTTGTGTTTTTTTGTTCCGGCGTCCGCATTTTTTTTTTTTTTTTTTTTTCTTAAGGGTGAGGTCGGTGGTTCAAGTCCACTCAGGCCTACCA | **Initial QC-passed read** |
| CTTAAGGGTGAGGTCGGTGGTTCAAGTCCACTCAGGCCTACCA | **trimmed read** |
| TAAGGGTGAGGTCGGTGGTTCAAGTCCACTCAGGCCTACCA | **–*local* alignment : removal of end mismatche(s)** |
| >SL-HBW:622:CA37GANXX:1:2212:17551:44544 | **Read ID** |
| TTTTACGGAGACCGCCCCTCTGTAGGTTGTCTTTTTTGTTTTTTTTTATTTTTCCGATTGGCGGTTTTTTTTTTTTTTTTCTTTTTAAGGGTGAGGTCGGTGGTTCAAGTCCACTCAGGCCTACCA | **Initial QC-passed read** |
| CTTTTTAAGGGTGAGGTCGGTGGTTCAAGTCCACTCAGGCCTACCA | **trimmed read** |
| TAAGGGTGAGGTCGGTGGTTCAAGTCCACTCAGGCCTACCA | **–*local* alignment : removal of end mismatche(s)** |
| >SL-HBW:622:CA37GANXX:1:2215:15263:77287 | **Read ID** |
| TTTTTTTTTTTTTTGTGTCTACCTTGTTGTTGCCTTTTTTTTTTTTTTTTTTTGGCGGTCTAAAAGTTGTGTTGTTTTTTTTTTTTAAGGGTGAGGTCCGTGGTTAAAGTCCACTCAGGCCTACCA | **Initial QC-passed read** |
| GGCGGTCTAAAAGTTGTGTTGTTTTTTTTTTTTAAGGGTGAGGTCCGTGGTTAAAGTCCACTCAGGCCTACCA | **trimmed read** |
| TAAGGGTGAGGTCCGTGGTTAAAGTCCACTCAGGCCTACCA | **–*local* alignment : removal of end mismatche(s)** |
| >SL-HBW:622:CA37GANXX:1:1101:14850:58375 | **Read ID** |
| TTTTTATTTTACCGTACCCTCCTTTTTTTTCTTGTTTGGGTTTCTCATGTCCGACGTACAACGGTTTTTTTTTTTTTTTTTTTTATAAGGGTGAGGTCGGTGGTTCAAGTCCACTCAGGCCTACCA | **Initial QC-passed read** |
| ATAAGGGTGAGGTCGGTGGTTCAAGTCCACTCAGGCCTACCA | **trimmed read** |
| ATAAGGGTGAGGTCGGTGGTTCAAGTCCACTCAGGCCTACCA | **–*local* alignment : removal of end mismatche(s)** |
| >SL-HBW:622:CA37GANXX:1:2310:17002:50237 | **Read ID** |
| TTTTTTTTATGTTGTTACTACCTAATTTTTTTCTTTCTTTTTTTTTTTTTCCGGAGTTCAAGGTTTTTTTTTTTTTTTTTTTTTATAAGGGTGAGGTCGGTGGTTCAAGTCCACTCAGGCCTACCA | **Initial QC-passed read** |
| ATAAGGGTGAGGTCGGTGGTTCAAGTCCACTCAGGCCTACCA | **trimmed read** |
| ATAAGGGTGAGGTCGGTGGTTCAAGTCCACTCAGGCCTACCA | **–*local* alignment : removal of end mismatche(s)** |
| >SL-HBW:622:CA37GANXX:1:2102:10660:58060 | **Read ID** |
| TTTTTGTGGTGGGGGGGTGCGTGTGTTTTTTGTGTTGTGGTGTTTTGGGTCCGGGGGTTCCGGGTTTTTTTTTTTTTTTTTTTTATAAGGGTGAGGTCGGTGGTTCAAGTCCACTCAGGCCTACCA | **Initial QC-passed read** |
| ATAAGGGTGAGGTCGGTGGTTCAAGTCCACTCAGGCCTACCA | **trimmed read** |
| ATAAGGGTGAGGTCGGTGGTTCAAGTCCACTCAGGCCTACCA | **–*local* alignment : removal of end mismatche(s)** |
| >SL-HBW:622:CA37GANXX:1:2208:5830:51255 | **Read ID** |
| TTTTGGTTTGGGTGTTTTTGTTGTGTTTTTTTTTTTGTTTTTTTTTTTTGGGCGTTTTCGGGGTTTTTTTTTTTTTTTTTTTTTATAAGGGTGAGGTCGGTGGTTCGAGTCCTCTCAGGCCTACCA | **Initial QC-passed read** |
| ATAAGGGTGAGGTCGGTGGTTCGAGTCCTCTCAGGCCTACCA | **trimmed read** |
| ATAAGGGTGAGGTCGGTGGTTCGAGTCCTCTCAGGCCTACCA | **–*local* alignment : removal of end mismatche(s)** |

**Supplemental Figure S3**

Illustration of how **ts** signals are generated: from initial QC-passed reads (the longest) to the trimmed/locally aligned reads (highlighted in color) that contribute to generate **ts** signals at three positions (in yellow, green and blue) along *ileT* tRNA. Only 11 reads are shown, representing a tiny fraction of all reads. It can be noticed that C misincorporations inside poly-Ts contribute to generate green reads.

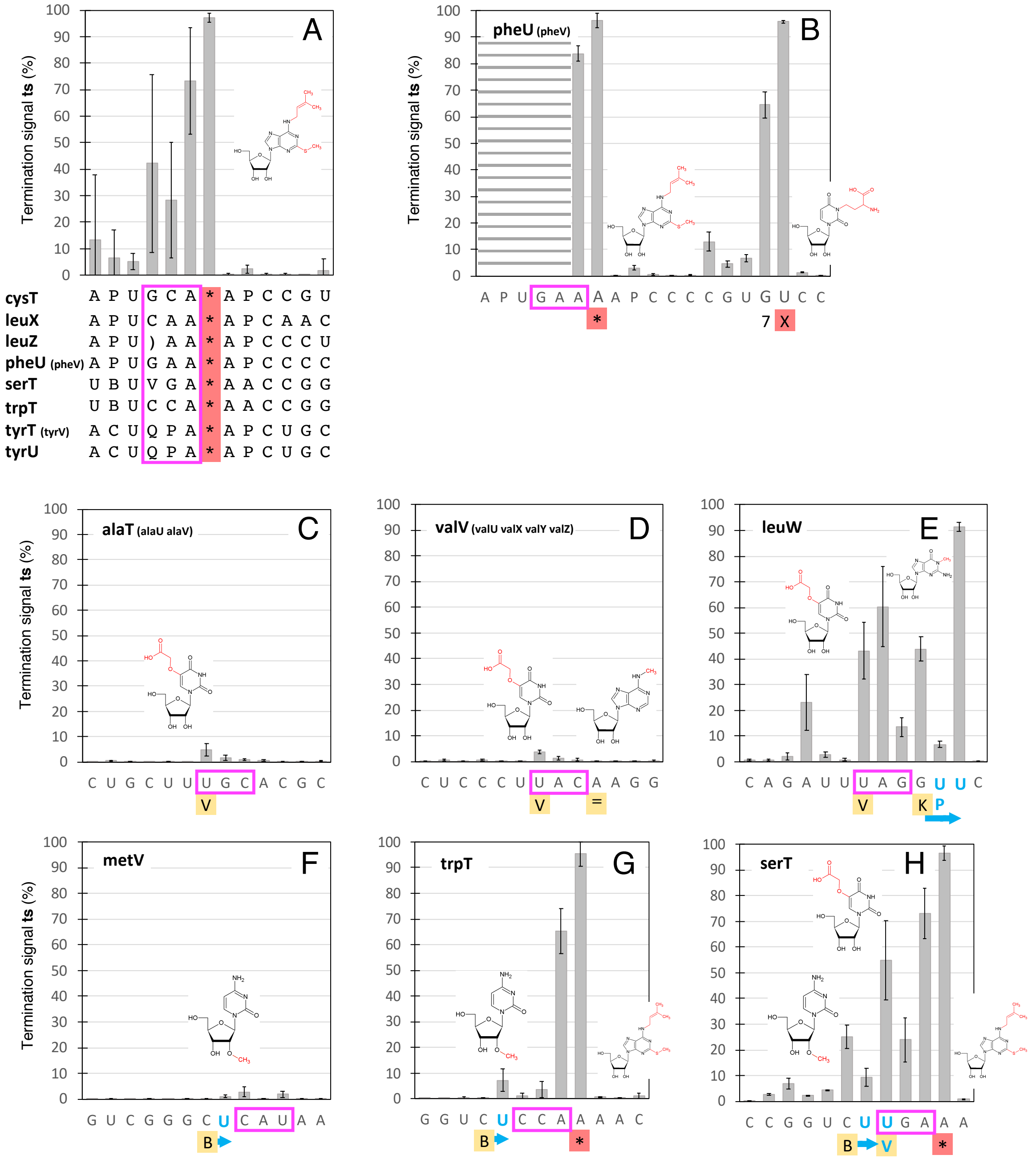


**Supplemental Figure S4**

**Ts** analysis of the anticodon loop: additional cases completing Figure 6. Except for panel **A**) (see below) the plots show the average values and standard deviations established from all 9 samples (Table 1), highlighting **ts** signatures of some typical *E. coli* combinations of modifications. Anticodons are boxed in magenta. Known modifications are reported from the *Modomics* database below the sequences (5’-3’). Highlighted modifications (in yellow and red) are illustrated above in the same order (illustrations are from the *Modomics* website). A blue arrow highlights a shift of a **ts** signal associated with a modification as a result of the presence of U residue(s) on its 3’ side, an artifact caused by poly-A computational trimming (see Material and Methods). **A**) Average values and standard deviations of the **ts** signal around the * modification from 8 different sequence contexts (in each context, average **ts** values are established from the 9 samples investigated in this study, Table 1). **B**) Sequence context with two modifications (* ad **X**) interfering strongly with the RT enzyme. The **ts** signal could not be established on the 5’ side of the * modification due to low coverage (dashed lines). **C-H**) **ts** signal associated with various combinations of modifications.

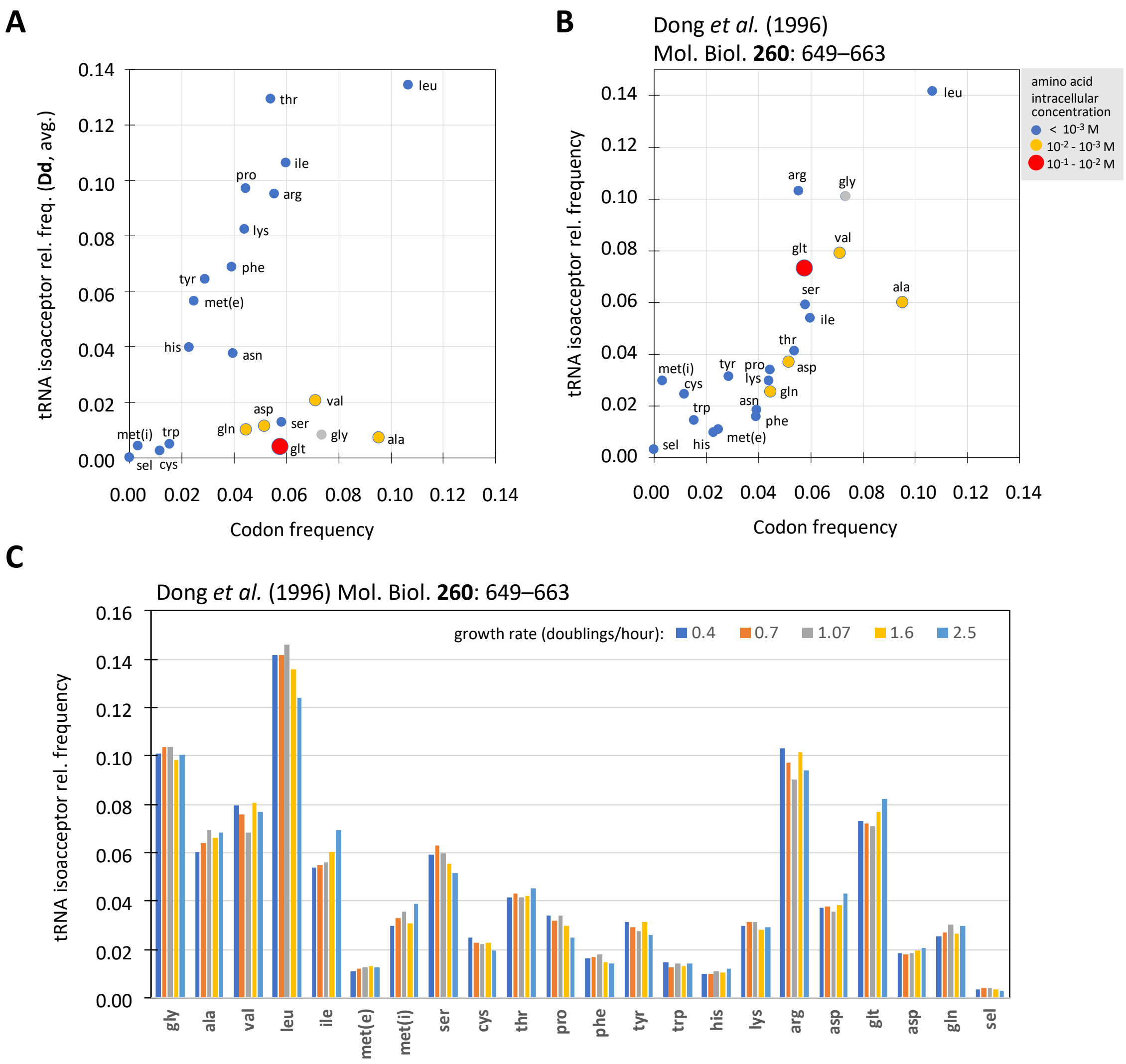


**Supplemental Figure S5**

*E. coli* tRNA isoacceptors relative frequency in cells: comparison between the relative frequencies obtained in the present study by deep-sequencing of CCA-ending tRNA (**A**) and the relative frequencies obtained through tRNA probe hybridization on gel by Dong et al. (1996) (**B**, **C**). **A**) Relative CCA-ending tRNA isoacceptor frequency by deep-sequencing as a function of codon frequency in *E. coli*. K-12 genome*; cultures samples at $OD_{600nm}$ ~ 0.4. Total RNA was DNase treated and deacylated (**Dd** samples). Values are in Supplemental File S6. **B**) Relative tRNA isoacceptor frequency at a growth rate of 0.4 (doubling/hour) as a function of codon frequency in *E. coli*. K-12 genome*. **C**) Relative tRNA isoacceptor frequency at growth rates of 0.4, 0.7, 1.07, 1.6 and 2.5 doubling/hour. Panels **B** and **C** were established from Table 5 in Dong et al. (1996). **Legend (top right):** intracellular concentrations of encoded amino acids in *E. coli* determined by Bennett *et al.* (2009). Cells grown under standard conditions, with glucose as the major source of carbon; cultures sampled at $OD_{650nm}$ = 0.35. Glycine is the only encoded amino acid for which the concentration was not determined in this study.

*Comments:*

Contrary to a statement** made in a study (Avcilar-Kucukgoze et al. 2016), panel **C** shows that the relative concentrations of tRNA species are surprisingly not depending on growth rate. Furthermore, we could not find any information in the original publication of Dong et al. (1996) that may explain the puzzling uniformity of these measurements, also considering experimental errors.

Note that exponential amplifications associated with deep sequencing experiments (panel **A**) alter true tRNA transcript frequencies, while the frequencies shown in **B** and **C** should in principle directly reflect these frequencies.

* data from https://openwetware.org/wiki/Escherichia_coli/Codon_usage

** *"Importantly, the abundance of each tRNA species changes non-uniformly with growth rate. For example, the levels of abundant tRNAs increase while minor tRNAs remain relatively unchanged when growth rate increases (Dong et al. 1996)"* (Avcilar-Kucukgoze et al. 2016).

| **Full name** | **Short name** | **One letter code** | ***Modomics* reference** |
|---|---|---|---|
| N6-methyladenosine | m6A | = | http://modomics.genesilico.pl/modifications/m6A/ |
| 2-methyladenosine | m2A | / | http://modomics.genesilico.pl/modifications/m2A/ |
| 5-carboxymethylaminomethyl-2′-O-methyluridine | cmnm5Um | ) | http://modomics.genesilico.pl/modifications/cmnm5Um/ |
| 5-methylaminomethyluridine | mnm5U | { | http://modomics.genesilico.pl/modifications/mnm5U/ |
| 2-lysidine | k2C | } | http://modomics.genesilico.pl/modifications/k2C/ |
| 5-carboxymethylaminomethyl-2-thiouridine | cmnm5s2U | $ | http://modomics.genesilico.pl/modifications/cmnm5s2U/ |
| 2-methylthio-N6-isopentenyladenosine | ms2i6A | * | http://modomics.genesilico.pl/modifications/ms2i6A/ |
| 2′-O-methylguanosine | Gm | # | http://modomics.genesilico.pl/modifications/Gm/ |
| 2-thiocytidine | s2C | % | http://modomics.genesilico.pl/modifications/s2C/ |
| N6-isopentenyladenosine | i6A | + | http://modomics.genesilico.pl/modifications/i6A/ |
| glutamyl-queuosine | gluQtRNA | ⊄ | http://modomics.genesilico.pl/modifications/gluQtRNA/ |
| 4-thiouridine | s4U | 4 | http://modomics.genesilico.pl/modifications/s4U/ |
| N6-threonylcarbamoyladenosine | t6A | 6 | http://modomics.genesilico.pl/modifications/t6A/ |
| 7-methylguanosine | m7G | 7 | http://modomics.genesilico.pl/modifications/m7G/ |
| 2′-O-methylcytidine | Cm | B | http://modomics.genesilico.pl/modifications/Cm/ |
| dihydrouridine | D | D | http://modomics.genesilico.pl/modifications/D/ |
| N6-methyl-N6-threonylcarbamoyladenosine | m6t6A | E | http://modomics.genesilico.pl/modifications/m6t6A/ |
| inosine | I | I | http://modomics.genesilico.pl/modifications/I/ |
| 2′-O-methyluridine | Um | J | http://modomics.genesilico.pl/modifications/Um/ |
| 1-methylguanosine | m1G | K | http://modomics.genesilico.pl/modifications/m1G/ |
| N4-acetylcytidine | ac4C | M | http://modomics.genesilico.pl/modifications/ac4C/ |
| pseudouridine | Y | P | http://modomics.genesilico.pl/modifications/Y/ |
| queuosine | QtRNA | Q | http://modomics.genesilico.pl/modifications/QtRNA/ |
| 5-methylaminomethyl-2-thiouridine | mnm5s2U | S | http://modomics.genesilico.pl/modifications/mnm5s2U/ |
| 5-methyluridine | m5U | T | http://modomics.genesilico.pl/modifications/m5U/ |
| uridine_5-oxyacetic_acid | cmo5U | V | http://modomics.genesilico.pl/modifications/cmo5U/ |
| 3-(3-amino-3-carboxypropyl)uridine | acp3U | X | http://modomics.genesilico.pl/modifications/acp3U/ |

**Supplemental Table S1**: Modified bases occuring on *E. coli* K-12 MG1655 tRNAs. These notations follow that of the *Modomics* database website http://modomics.genesilico.pl/ (Boccaletto et al. 2018).
