## Supplemental File S1 for "Highly reproducible characterization of *Escherichia coli* tRNA epitranscriptome with a simple method of library preparation for deep sequencing"

### Library preparation from total RNA: full protocol

**NOTES**:

- It is essential to work in an RNase-free environment.
- Shown values are actual measurements related to the 9 samples processed and analyzed in the present work: 3 biological replicates (or batches) x 3 different total RNA treatments.
- It was initially planned to investigate tRNA aminoacylation by applying oxidation and deacylation treatments to additional samples (**DOd** and **DdO** samples). This part of the analysis was finally not pursued. For the sake of presenting unaltered gel results, lanes corresponding to these samples may still be shown.

---

#### Total RNA extraction from *Escherichia coli* (*E. coli*) MG1655

**1. Pre-culture preparation**

1) Add 5 mg RNase A powder (supplied in plasmid purification kit, MN) into 250 ml sterilized LB medium and incubate at 37°C for at least 48 hours (ensures an absence of LB contamination).
2) Pick the *E. coli* MG1655 stock and streak at least two times on LB plate.
3) Pick a single clone and incubate in 6 ml LB medium at 37°C/180 rpm overnight. Prepare as many independent cultures as required by the investigation.
4) Set a blank control to test for contamination.

**2. Total RNA extraction**

1) Prepare 4 Erlenmeyer flasks, and fill them with 50 ml RNase A-treated LB medium.
2) Slightly vortex the cultures and pipet 500 µl into Erlenmeyer flasks. Measure the initial $OD_{600}$.
3) Measure the $OD_{600}$ every 15-30 min and harvest the cells at $OD_{600}$ about 0.4.

| Time \ $OD_{600}$ | Batch A | Batch B | Batch C |
|---|---|---|---|
| 0 min | 0.05 | 0.04 | 0.05 |
| 30 min | 0.12 | 0.13 | 0.13 |
| 60 min | 0.28 | 0.29 | 0.31 |
| 75 min | 0.45 | 0.45 | 0.45 |
| *Cells were harvested at 75 min* | | | |

4) Use 2 ml Eppendorf tube; harvest 2 ml of culture by centrifugation 1 min at maximum speed; discard the supernatant.
5) Immediately add 1 ml TRIZOL reagent and vortex to re-suspend the bacteria pellet.
6) Keep the homogenate on ice for 30 min.
7) Add 200 µl chloroform and mix well by turning the tube up and down for about 15 seconds.
8) Keep the tube on ice for about 10 min and centrifuge at 12,000*g* for 15 min at 4°C.
9) Carefully pipet around 400~450 µl of the upper phase into a new RNase-free tube; add a same volume of Isopropanol and vortex.

10) Keep the tube at -20°C for 1 h and centrifuge at 12,000*g* for 15 min at 4°C.
11) Carefully discard the supernatant. Add 1 µl glycogen (ThermoFisher), 5 µl NaOAc/HOAc buffer (Sigma-Aldrich), 44 µl RNase-free water and 125 µl 100% Ethanol. Incubate at -20°C for at least 45 min.
12) Centrifuge at maximum speed for 30 min at 4°C; discard the supernatant. Add 125 µl 70% Ethanol and centrifuge at the same speed for 15 min.
13) Air-dry the pellet and suspend in 20 µl water.

#### 3. Verification of the RNA quality

1) Run total RNA on a 1% agarose gel, 135 Volts, 25 min.

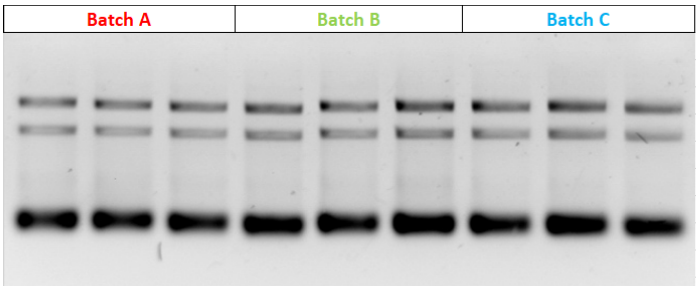


2) Measure the concentration; Store the sample at -20°C and use them the next day or as soon as possible.

| Batches | Batch A | Batch B | Batch C |
|---|---|---|---|
| Concentration (µg/µl) | 1.259 | 1.326 | 1.360 |

#### 4. DNase treatment (D and Dd samples)

| | |
|---|---|
| RNA in $H_2O$ (around 8 µg) | 87 µl |
| 10X buffer TurboDNase | 10 µl |
| TurboDNase (Ambion) | 2 µl |
| DTT (0.1 M) | 1 µl |
| Total | 100 µl |

1) Incubate 4h at 37°C then keep at 4°C overnight. Phenol/chloroform extraction achieved subsequently.
2) Run a gel to check the quality of RNA.
Run 4% agarose gel to check the quality of RNA. 135 volts for 25 min. Load 1 µl RNA solution.

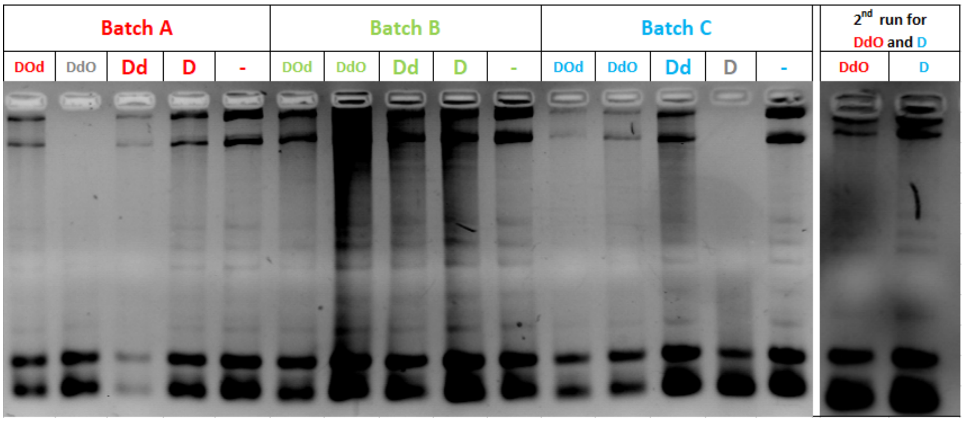

3) Establish the concentration of all DNase-treated samples. Add 1 µg in each tube ready for RNA ligation (6 tubes: **AD**, **BD**, **CD**, **ADd**, **BDd**, **CDd**; **d**eacylation of **Dd** samples is performed in a subsequent step). Meanwhile, add 1 µg of non-DNase treated RNA into the remaining 3 tubes (**A-**, **B-**, **C-**).

| Sample | Volume | Concentration | Volume for RNA ligation | Quantity for RNA ligation | Optional spike for quantification purposes (not used in our analysis) | Quantity of Spike |
|---|---|---|---|---|---|---|
| **ADd** | 20 µl | 261.19 ng/µl | 3.83 µl | 1 µg | 1 µl | 1.3 ng |
| **AD** | 20 µl | 467.36 ng/µl | 2.14 µl | 1 µg | 1 µl | 1.3 ng |
| **A-** | 20 µl | 481.92 ng/µl | 2.08 µl | 1 µg | 1 µl | 1.3 ng |
| **BDd** | 20 µl | 399.32 ng/µl | 2.50 µl | 1 µg | 1 µl | 1.3 ng |
| **BD** | 20 µl | 492.02 ng/µl | 2.03 µl | 1 µg | 1 µl | 1.3 ng |
| **B-** | 20 µl | 510.10 ng/µl | 1.96 µl | 1 µg | 1 µl | 1.3 ng |
| **CDd** | 20 µl | 454.01 ng/µl | 2.20 µl | 1 µg | 1 µl | 1.3 ng |
| **CD** | 20 µl | 470.76 ng/µl | 2.12 µl | 1 µg | 1 µl | 1.3 ng |
| **C-** | 20 µl | 475.48 ng/µl | 2.10 µl | 1 µg | 1 µl | 1.3 ng |

#### 5. PCR to verify the absence of DNA

1) Use primers tRNA-tyr5' / tRNA-tyr3' (also add a negative control)

| | |
|---|---|
| RNA sample | 1 µl |
| DreamTaq polymerase | 0.1 µl |
| 10X DreamTaq buffer | 2.5 µl |
| dNTPs (5mM) | 2 µl |
| Primer Tyr F (10 µM) | 1 µl |
| Primer Tyr R (10 µM) | 1 µl |
| $H_2O$ | 17.5 µl |
| Total | 25µl |

| PCR program | | |
|---|---|---|
| 95°C | 2' | |
| 95°C | 15'' | X 30 cycles |
| 55°C | 15'' | |
| 72°C | 15'' | |
| 72°C | 2' | |
| 12°C | ∞ | |

2) Analysis of the amplification products on 4% agarose gel, 135 Volts, 25 min.

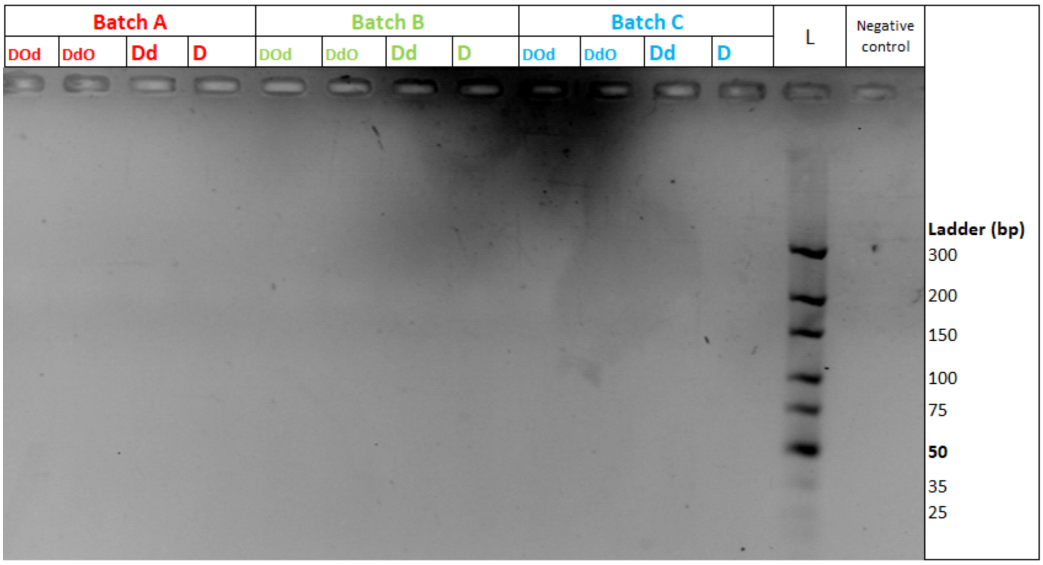

#### RNA-seq library construction protocol

##### 1. RNA ligation

| | |
|---|---|
| RNA treated (**Dd**, **D** or **-**) (1 µg per tube) | 5 µl |
| Pre-adenylated RNA 3' adapter (100 µM) | 1 µl |
| Mix well and then place on PCR machine, denature at 70°C for 2 min then chill on ice immediately. After denaturation, the tube **MUST** be kept on ice. | |
| 10x T4 RNA ligase 2 buffer | 1 µl |
| DMSO | 1 µl |
| BSA | 1 µl |
| T4 RNA ligase 2 truncated (NEB) | 1 µl |
| $H_2O$ | Fill up to 10 µl |

Incubate at 10°C overnight; Add 40 µl $H_2O$, then EtOH precipitate.
Note: After adding ethanol, add 1 µl RT primer (100 µM) in the solution. After precipitation, re-suspend the pellet into 13 µl water.

##### 2. Reverse Transcription

| | |
|---|---|
| RNA ligation pellet | 13 µl |
| dNTPs mix (10 mM) | 1 µl |
| Put the tube at 70°C for 2 min then chill on ice immediately | |
| 5X RT Buffer | 4 µl |
| DTT 0.1M | 1 µl |
| SuperScript III (200 u/µl) | 1 µl |
| Total | 20µl |

Mix well and keep at r.t. for 2 min and incubate 1h at 55°C then cool down to r.t.

##### 3. PCR the cDNA to check RNA ligation

In this step, a $tRNA^{Tyr}$ is amplified with a primer specific to its 5'end and a TruSeq-RT primer specific to the 3' adapter.

| | |
|---|---|
| RT product | 0.5 µl |
| DreamTaq polymerase | 0.1 µl |
| 10X DreamTaq buffer | 2.5 µl |
| dNTPs (5mM) | 2 µl |
| TruSeq-RT Primer (10µM) | 1 µl |
| Primer Tyr 5'R (10 µM) | 1 µl |
| $H_2O$ | 18.5 µl |
| Total | 25µl |

| PCR program | | |
|---|---|---|
| 94°C | 2' | |
| 94°C | 20'' | X 30 cycles |
| 55°C | 20'' | |
| 72°C | 20'' | |
| 72°C | 2' | |
| 12°C | ∞ | |

Run a 3% agarose gel, 135 Volts, 25 min

Expected size = 107 bp :

5'-GGTGGGGTTCCCGAGCGGCCAAAGGGAGCAGACTGTAAATCTGCCGTCACAGACTTCGAAGGTTCGA
ATCCTTCCCCCACCACCATGGAATTCTCGGGTGCCAAGGC-3'

(The adaptor sequence is in green).

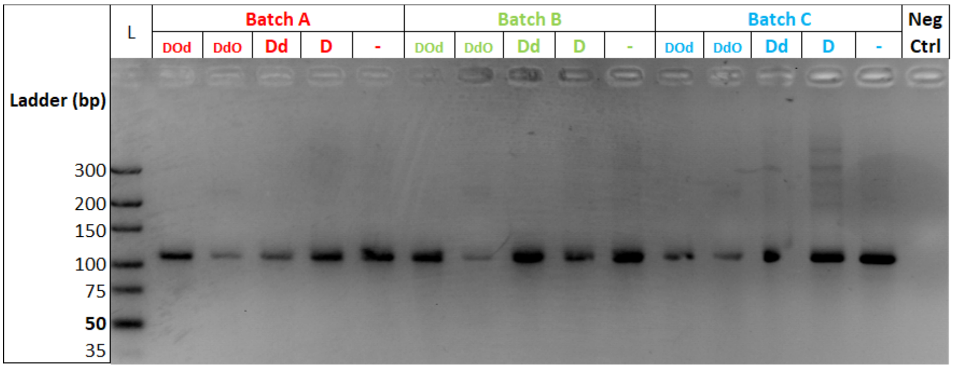


#### 4. Run a gel for RT products purification

3% agarose gel with 12 wells combo.
Use fresh 0.5X TAE running buffer.
Load the RT products and then run the gel at 135 V for 5 min. Then 100 V for 35 min.
Cut the gel from ~35bp to 200 bp (avoid the large band corresponding to the excess of adapter)

**Batch A**

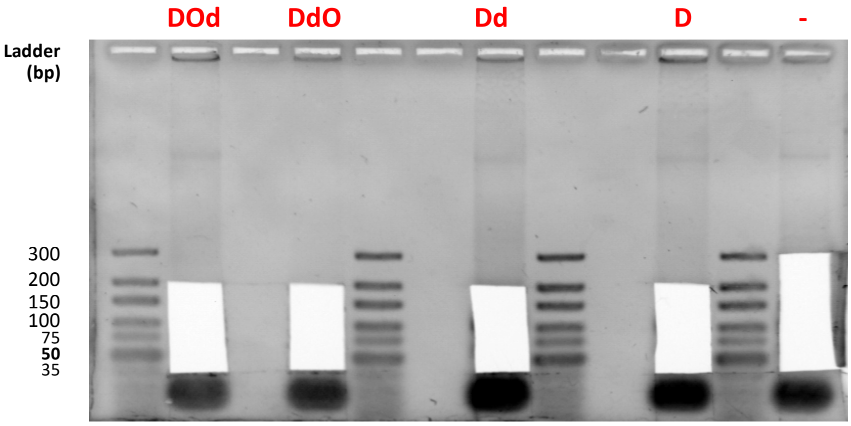

Batch B

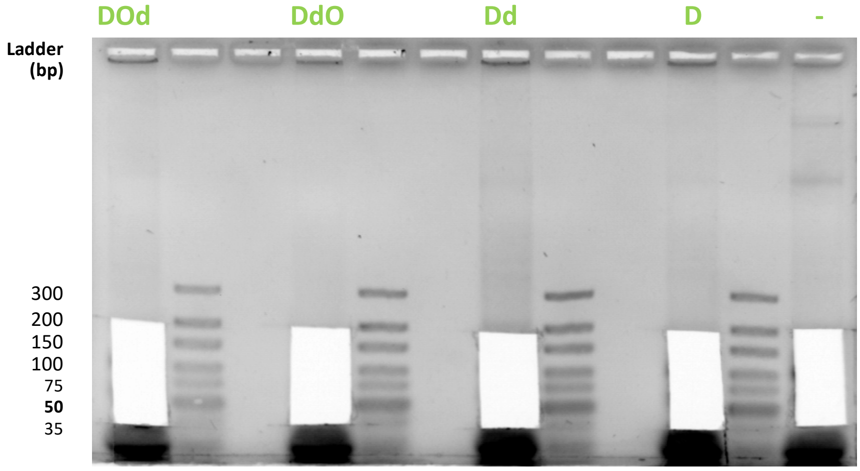


Batch C

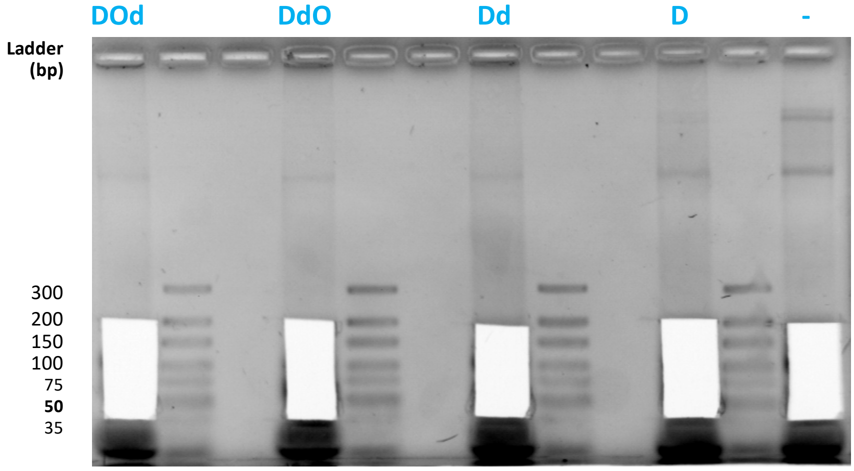


With a 0.8mm needle, punch the bottom of 0.5 ml tubes 3-4 times to make small holes. Cut the gel and put it into the 0.5ml gel breaker tube, then cover it with a 2ml tube; Centrifuge at maximum speed for 15 min. Resuspend well with 350 µl $H_2O$ + 50 µl TE buffer per tube and shake overnight at 20°C.

**5. cDNA recovery**

1) Transfer the gel solution into a SPIN-X costar filter tube, then centrifuge at maximum speed for 15 min and EtOH Precipitation.

2) After centrifugation, resuspend the pellet in 22 µl water, take 1 µl to measure the concentration.

| Sample | Volume | Concentration | Volume for TdT | Quantity for TdT |
|---|---|---|---|---|
| ADd | 20 µl | 32.25 ng/µl | 6.20 µl | 200 ng |
| AD | 20 µl | 18.33 ng/µl | 10.91 µl | 200 ng |
| A- | 20 µl | 15.64 ng/µl | 12.79 µl | 200 ng |
| BDd | 20 µl | 40.17 ng/µl | 5.00 µl | 200 ng |
| BD | 20 µl | 20.71 ng/µl | 9.66 µl | 200 ng |
| B- | 20 µl | 15.63 ng/µl | 12.80 µl | 200 ng |
| CDd | 20 µl | 22.20 ng/µl | 9.01 µl | 200 ng |
| CD | 20 µl | 22.13 ng/µl | 9.04 µl | 200 ng |
| C- | 20 µl | 19.63 ng/µl | 10.19 µl | 200 ng |

**6. Poly-A tail addition at cDNA 3' end with TdT**

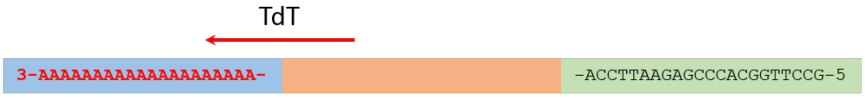


| | | |
|---|---|---|
| 5 pmol cDNA | 200 ng | (<7 pmol) |
| 10X TdT buffer | 5 µl | |
| $CoCl_2$ (2.5 mM) | 5 µl | |
| 10 mM dATP | 1 µl | (10 nmol) |
| TdT enzyme (NEB) | 0.5 µl | |
| $H_2O$ | Fill up to 50 µl | |
| Total | 50 µl | |

Incubate at 37°C for 45 min. Stop the reaction by heating up to 70°C for 10 min
According to NEB manual instructions, 10-30 A will be added.
Use a Sigma spin column to clean the reaction.

**7. EtOH Precipitation**

After centrifugation, resuspend the pellet in 22 µl $H_2O$; take 1 µl to establish the concentration.

| Sample | Volume | Concent. | Sample | Volume | Concent. | Sample | Volume | Concent. |
|---|---|---|---|---|---|---|---|---|
| ADd | 20 µl | 8.04 ng/µl | ADd | 20 µl | 8.34 ng/µl | ADd | 20 µl | 5.73 ng/µl |
| AD | 20 µl | 17.10 ng/µl | AD | 20 µl | 9.32 ng/µl | AD | 20 µl | 6.11 ng/µl |
| A- | 20 µl | 10.29 ng/µl | A- | 20 µl | 16.54 ng/µl | A- | 20 µl | 6.36 ng/µl |

#### 8. PCR preamplification

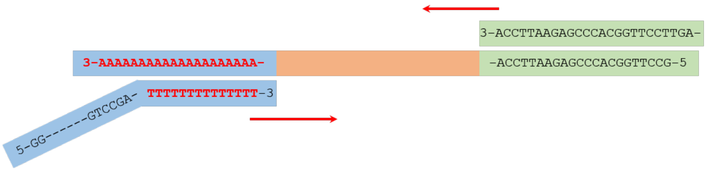


| | |
|---|---|
| $H_2O$ | 27 µl |
| 10X Taq buffer | 5 µl |
| $MgCl_2$ | 4 µl |
| dNTPs (5mM) | 4 µl |
| Illumina-polyT-5N primer (10 µM) | 2 µl |
| Template (50 ng) | ~4 µl |
| Heat up the mix to 94°C, then cool down to r.t. | |
| RT primer +TGG (10 µM) | 2 µl |
| Taq polymerase | 2 µl |
| Total | 50 µl |

| PCR program | | |
|---|---|---|
| 94°C | 2’ | |
| 94°C | 30’’ | 10~18 cycles |
| 55°C | 30’’ | |
| 72°C | 45’’ | |
| 72°C | 2’ | |
| 12°C | ∞ | |

1) Verification of the amplification with gel analysis, 4 % agarose gel. Load 5 µl of each solution.

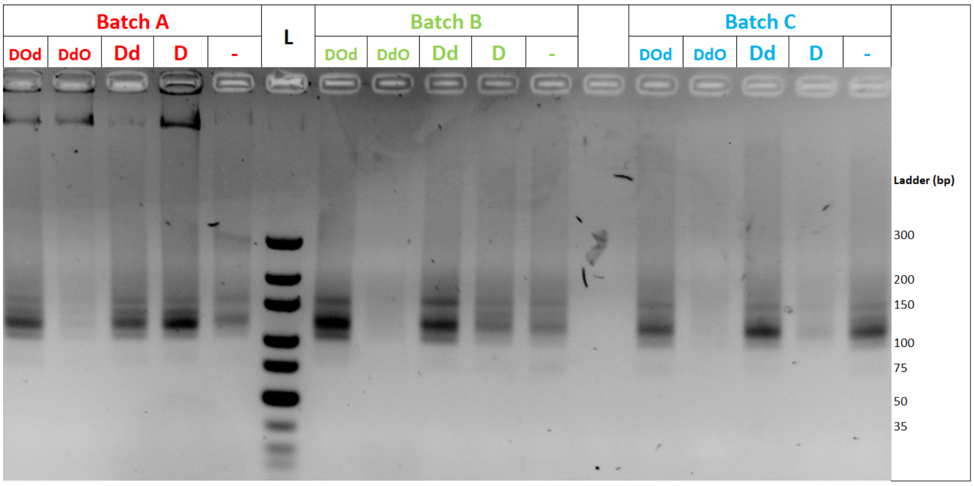


2) Purification of the pre-PCR products
Using MN *NucleoSpin*® column, clean the reaction solutions.
In the last step of purification, use 30 µl fresh elution buffer to wash the column three times.
Add 10ul NaOAc/HOAc buffer + 1 µl glycogen into 90 µl purified PCR product solution and precipitate. Suspend the pellet in 20 µl $H_2O$ and 5 µl TE buffer.

| Sample | Volume | Concent. | Sample | Volume | Concent. | Sample | Volume | Concent. |
|---|---|---|---|---|---|---|---|---|
| Dd | 25 µl | 53.81 ng/µl | Dd | 25 µl | 95.76 ng/µl | Dd | 25 µl | 67.96 ng/µl |
| D | 25 µl | 70.95 ng/µl | D | 25 µl | 55.33 ng/µl | D | 25 µl | 40.12 ng/µl |
| - | 25 µl | 55.82 ng/µl | - | 25 µl | 72.33 ng/µl | - | 25 µl | 68.06 ng/µl |

##### 9. PCR amplification with tag sequence addition

1) Each sample has a different tag (O) to allows multiplexing in Illumina deep-sequencing experiment. Verification of the amplification with gel analysis; 4 % agarose gel. Load 10 µl of each solution.

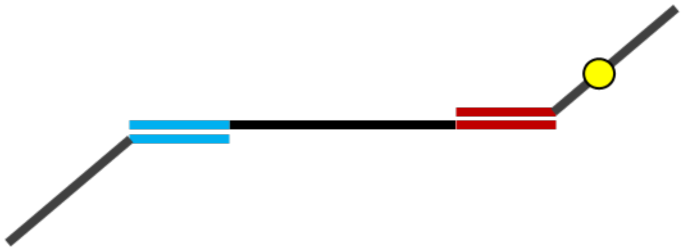

| | |
|---|---|
| dNTPs (5 mM) | 2 µl |
| 10X AccuPrime Buffer I | 5 µl |
| $MgCl_2$ (50 mM) | 0.5 µl |
| Universal Primer (10 µM) | 2 µl |
| Illumina Tag Index (10 µM) | 2 µl |
| Pre-PCR Template | 100 ng |
| AccuPrime Taq | 1 µl |
| $H_2O$ | Full fill to 50 µl |
| Total | 50 µl |

| PCR program | | |
|---|---|---|
| 94°C | 2′ | |
| 94°C | 30′′ | 5~15 cycles |
| 59°C | 30′′ | |
| 68°C | 45′′ | |
| 68°C | 2′ | |
| 12°C | ∞ | |

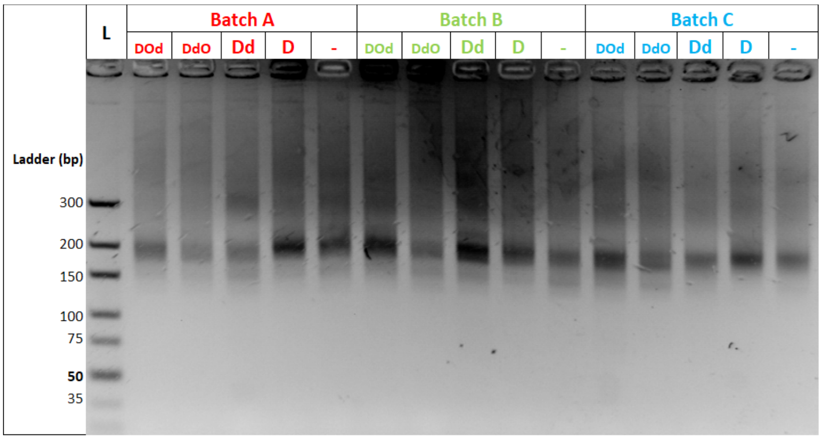


2) PCR products purification
Using MN *NucleoSpin*® column, clean up the reaction solution.
Use 30 µl of fresh elution buffer to wash the column; repeat three times.
Precipitate and suspend the pellet into 25 µl $H_2O$ and 5 µl TE buffer.

| Sample | Volume | Conc. | Sample | Volume | Conc. | Sample | Volume | Conc. |
|---|---|---|---|---|---|---|---|---|
| Dd | 30 µl | 95.20 ng/µL | Dd | 30 µl | 97.75 ng/µL | Dd | 30 µl | 83.01 ng/µL |
| D | 30 µl | 82.71 ng/µL | D | 30 µl | 67.11 ng/µL | D | 30 µl | 77.86 ng/µL |
| - | 30 µl | 100.58 ng/µL | - | 30 µl | 69.22 ng/µL | - | 30 µl | 60.23 ng/µL |

These samples are ready for *Illumina* deep-sequencing analysis. With multiplexing, it is recommended to establish the concentrations very carefully so that a mixture with equal amount of each sample can be prepared. This will ensure the best possible comparison between the signals obtained from each sample.
