## Supplemental File S4 for "Highly reproducible characterization of *Escherichia coli* tRNA epitranscriptome with a simple method of library preparation for deep sequencing"

Termination signal (%) of all tRNA transcripts and three tRNA pseudogenes transcripts (pauD, pawZ, ptwF) from *Escherichia coli MG1655.* This file presents details of the signals shown in Figure 2e. Each bar is calculated from the combined coverage of the 9 samples examined in the present study, while dashes represent sample values. Only batches can be differentiated (by color): A**D**, A**Dd**, A**-**, B**D**, B**Dd**, B**-**, C**D**, C**Dd** and C**-**. Note that because the tRNA coverage in a sample may drop below 100 before the 5' end, individual **ts** signals (established only when the coverage is ≥ 100) may not be present although the combined coverage allows to establish a weighted value of the **ts** signal. Identical tRNA genes are represented only once (copies are listed in brackets).

**alaT (alaU alaV)**

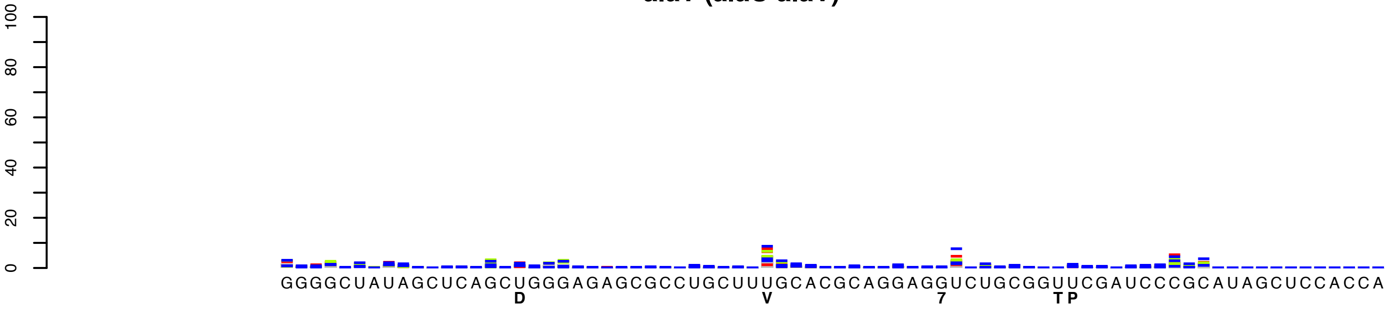


**alaX (alaW)**

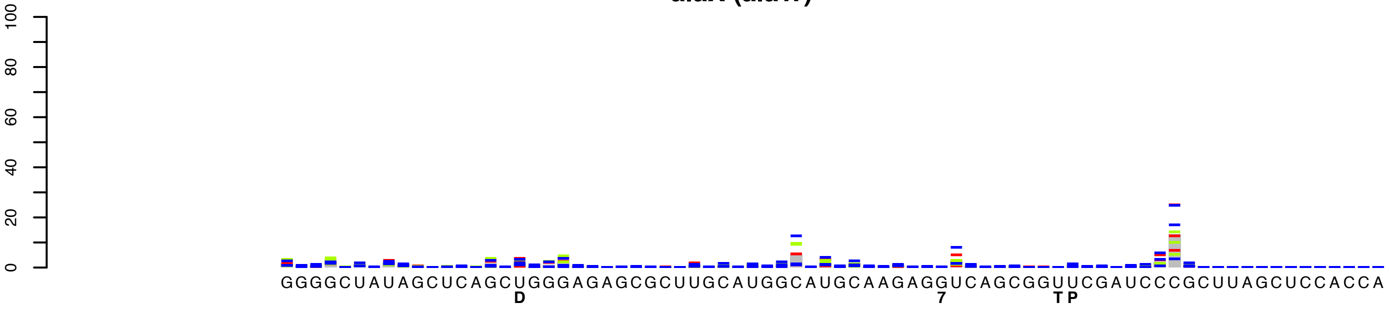

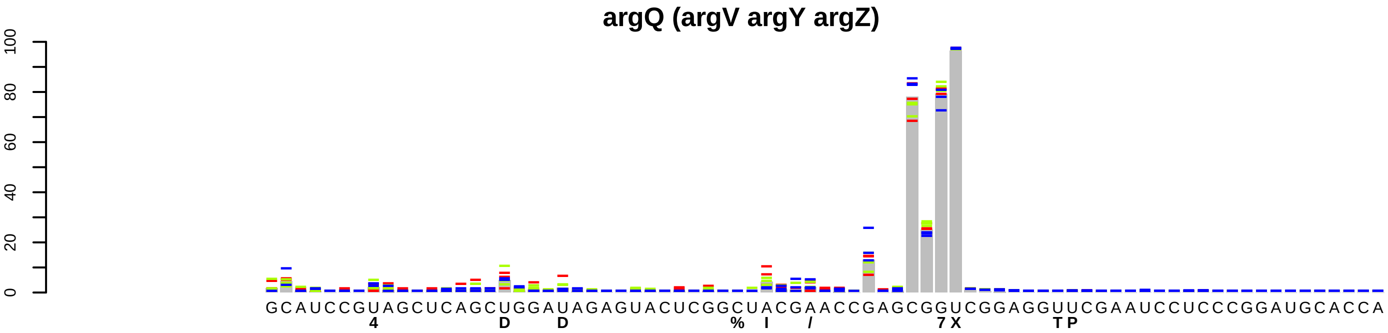
argQ (argV argY argZ)
0 20 40 60 80 100
GCAUCCGUAGCUCAGCUGGAUAGAGUACUCGGCUACGAACCGAGCGGUCGGAGGUUCGAAUCCUCCCGGAUGCACCA
4 D D % I / 7X TP

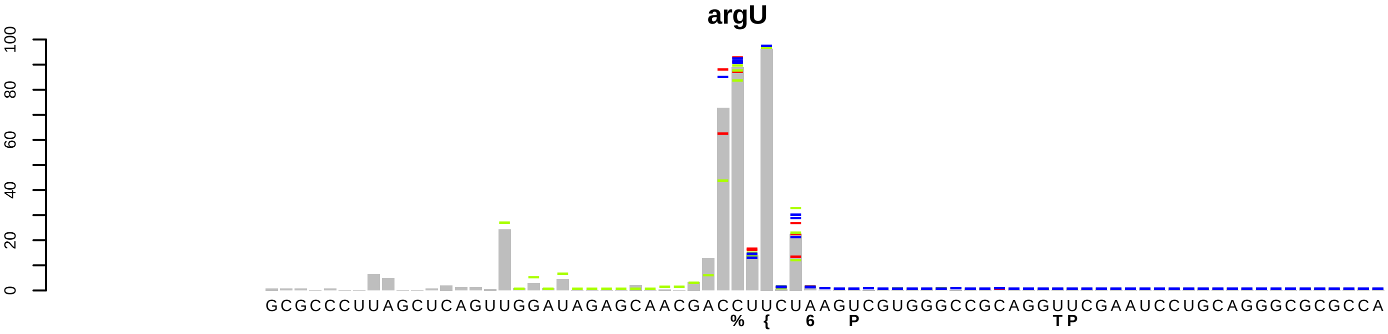
argU
0 20 40 60 80 100
GCGCCCUUAGCUCAGUUGGAUAGAGCAACGACCUUCUAAGUCGUGGGCCGCAGGUUCGAAUCCUGCAGGGCGCGCCA
% { 6 P TP

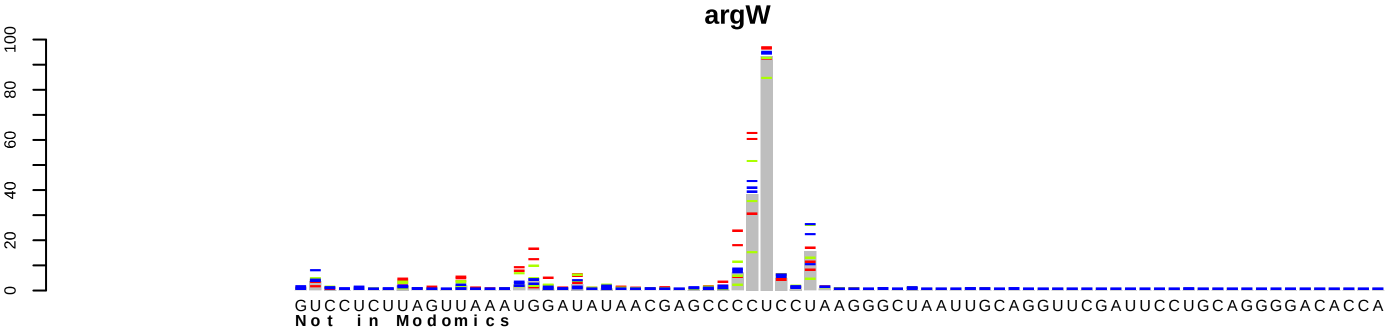
argW
0 20 40 60 80 100
GUCCUCUUAGUUAAAUGGAUAUAACGAGCCCCUCCUAAGGGCUAAUUGCAGGUUCGAUUCCUGCAGGGGACACCA
Not in Modomics

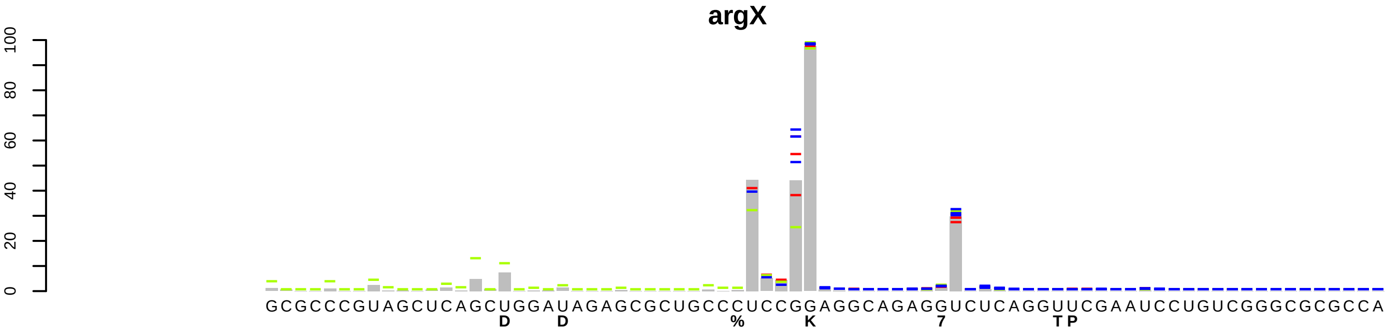
argX
0 20 40 60 80 100
GCGCCCGUAGCUCAGCUGGAUAGAGCGCUGCCCUCCGGAGGCAGAGGUCUCAGGUUCGAAUCCUGUCGGGCGCGCCA
D D % K 7 TP

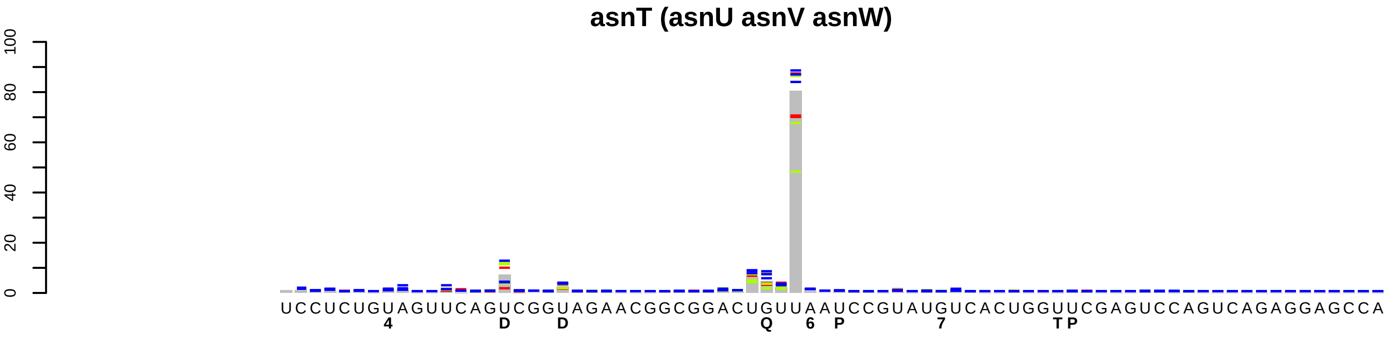
asnT (asnU asnV asnW)
0
20
40
60
80
100
UCCUCUGUAGUUCAGUCGGUAGAACGGCGGACUGUUAAUCCGUAUGUCACUGGUUCGAGUCCAGUCAGAGGAGCCA
4
D
D
Q
6
P
7
T P

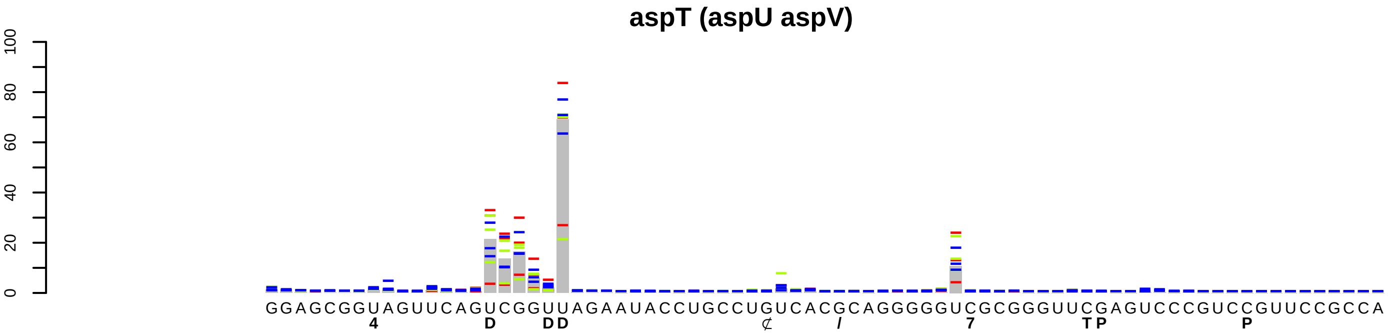

aspT (aspU aspV)
100
80
60
40
20
0
GGAGCGGUAGUUCAGUCGGUUAGAAUACCUGCCUGUCACGCAGGGGGUCGCGGGUUCGAGUCCCGUCCGUUCCGCCA
4
D
DD
7
TP
P

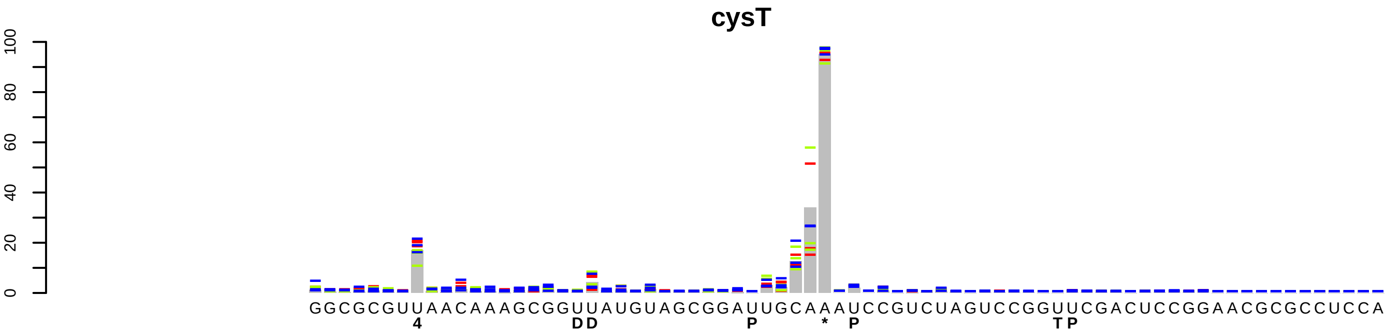
cysT
0
20
40
60
80
100
GGCGCGUUAACAAAGCGGUUAUGUAGCGGAUUGCAAAUCCGUCUAGUCCGGUUCGACUCCGGAACGCGCCUCCA
4
DD
P
*
P
TP

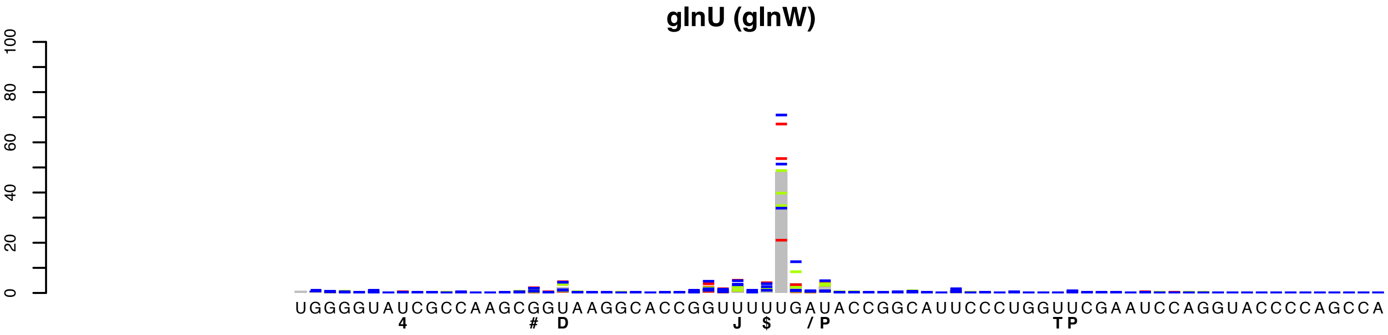
glnU (glnW)
0
20
40
60
80
100
UGGGGUAUCGCCAAGCGGUAAGGCACCGGUUUUUGAUACCGGCAUUCCCUGGUUCGAAUCCAGGUACCCCAGCCA
4
#
D
J
$
/
P
T
P

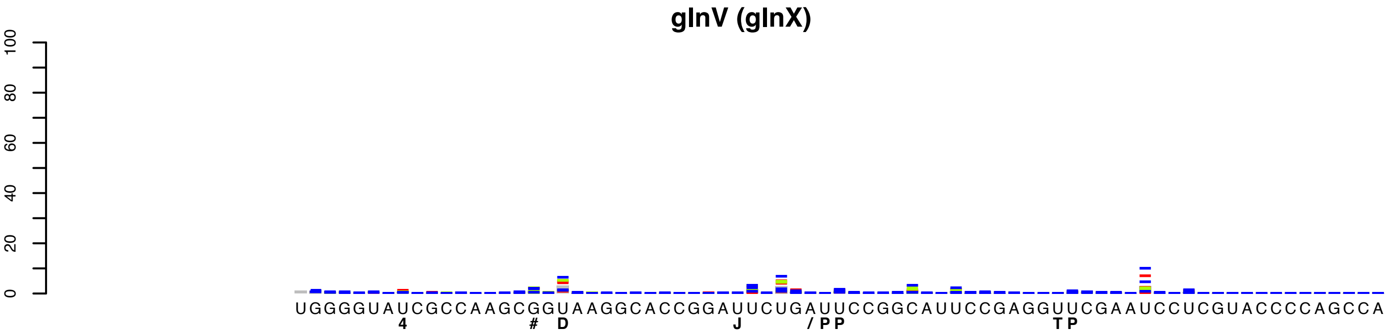
glnV (glnX)
0
20
40
60
80
100
UGGGGUAUCGCCAAGCGGUAAGGCACCGGAUUCUGAUUCCGGCAUUCCGAGGUUCGAAUCCUCGUACCCCAGCCA
4
#
D
J
/
P
P
T
P

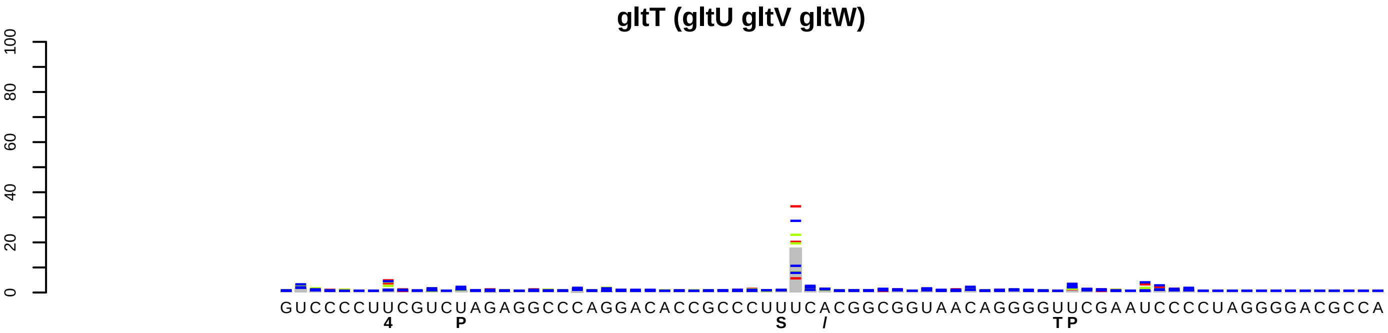
gltT (gltU gltV gltW)
0
20
40
60
80
100
GUCCCCUUCGUCUAGAGGCCCAGGACACCGCCCUUUCACGGCGGUAACAGGGGUUCGAAUCCCCUAGGGGACGCCA
4
P
S
/
T P

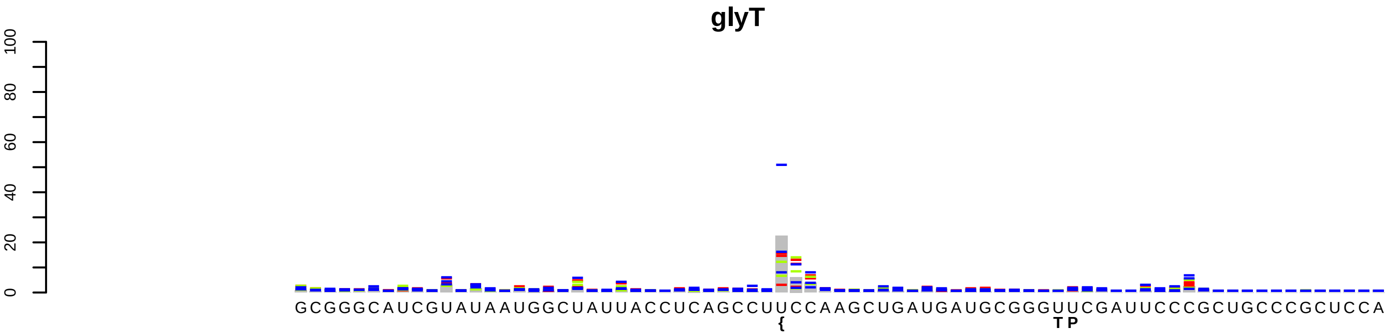
glyT
0 20 40 60 80 100
GCGGGCAUCGUAUAAUGGCUAUUACCUCAGCCUUCCAAGCUGAUGAUGCGGGUUCGAUUCCCGCUGCCCGCUCCA
{
T P

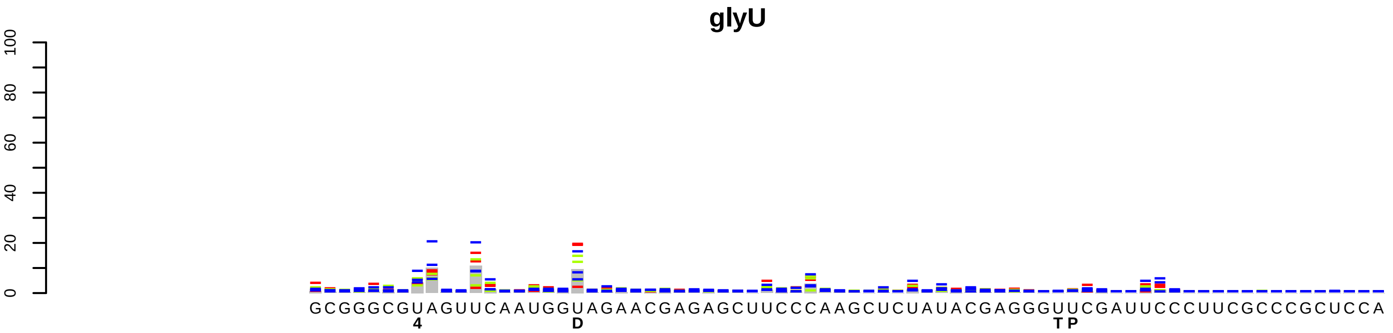
glyU
0 20 40 60 80 100
GCGGGCGUAGUUCAAUGGUAGAACGAGAGCUUCCCAAGCUCUAUACGAGGGUUCGAUUCCCUUCGCCCGCUCCA
4
D
T P

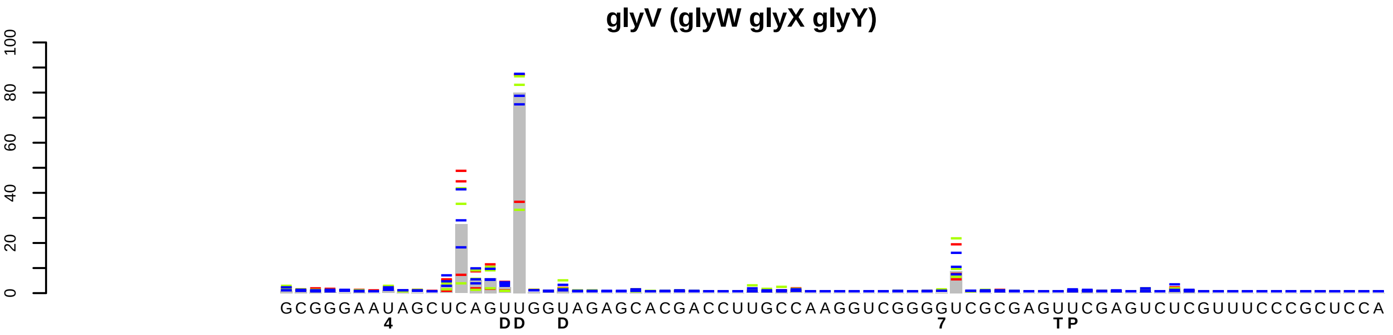
glyV (glyW glyX glyY)
0 20 40 60 80 100
GCGGGAAUAGCUCAGUUGGUAGAGCACGACCUUGCCAAGGUCGGGGUCGCGAGUUCGAGUCUCGUUUCCCGCUCCA
4
DD
D
7
T P

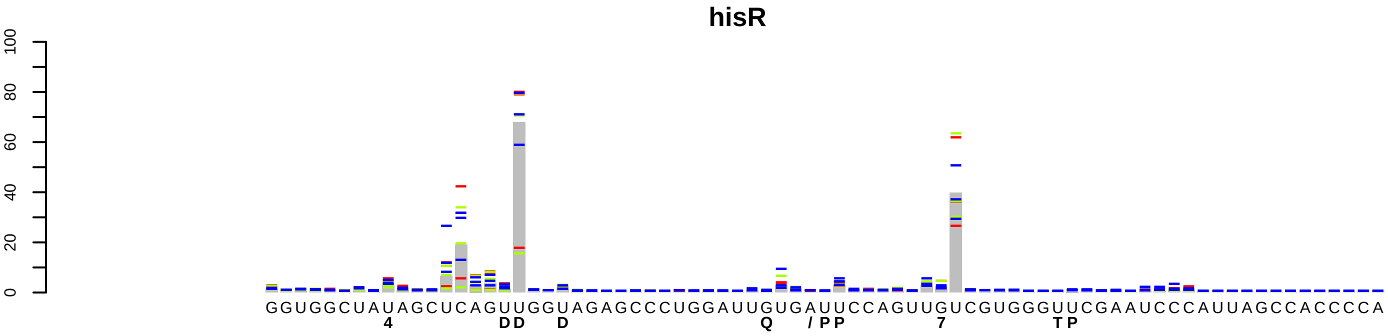
hisR
100
80
60
40
20
0
GGUGGCUAUAGCUCAGUUGGUAGAGCCCUGGAUUGUGAUUCCAGUUGUCGUGGGUUCGAAUCCCAUUAGCCACCCCA
4 DD D Q / PP 7 TP

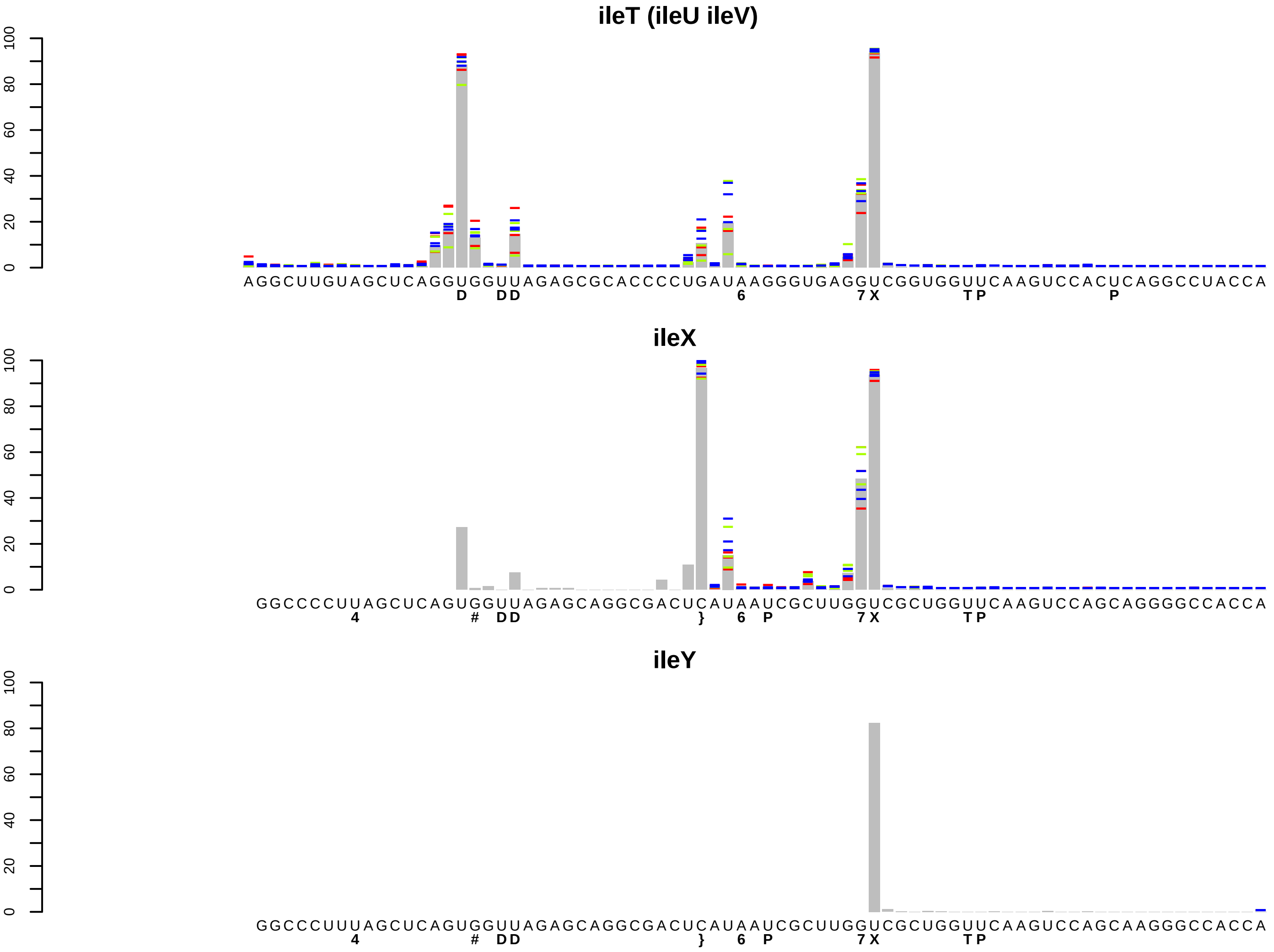
ileT (ileU ileV)
0 20 40 60 80 100
AGGCUUGUAGCUCAGGUGGUUAGAGCGCACCCCUGAUAAGGGUGAGGUCGGUGGUUCAAGUCCACUCAGGCCUACCA
D DD 6 7X TP P
ileX
0 20 40 60 80 100
GGCCCCUUAGCUCAGUGGUUAGAGCAGGCGACUCAUAAUCGCUUGGUCGCUGGUUCAAGUCCAGCAGGGGCCACCA
4 # DD } 6 P 7X TP
ileY
0 20 40 60 80 100
GGCCCUUUAGCUCAGUGGUUAGAGCAGGCGACUCAUAAUCGCUUGGUCGCUGGUUCAAGUCCAGCAAGGCCACCA
4 # DD } 6 P 7X TP

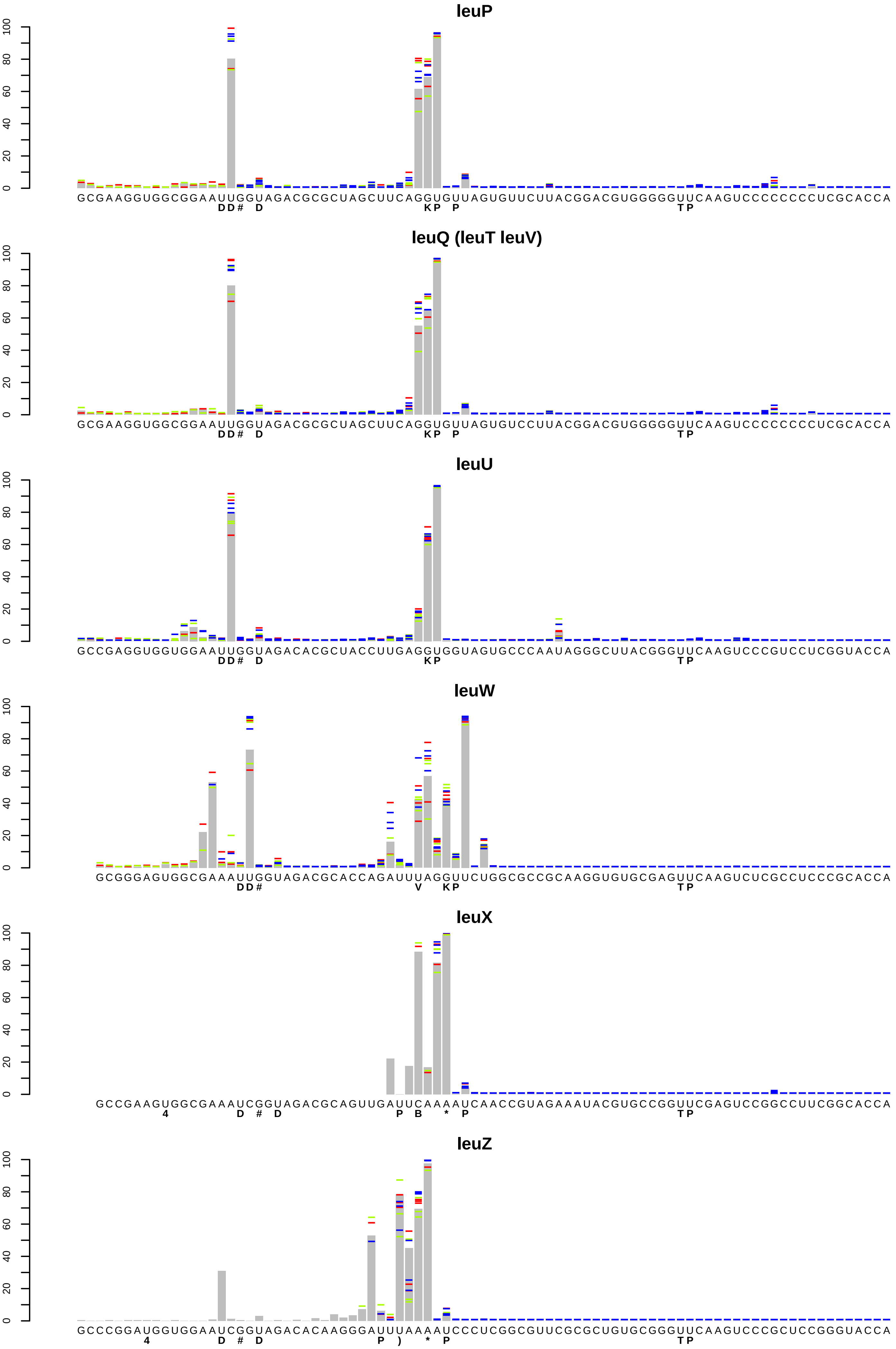

leuP
GCGAAGGUGGCGGAAUUGGUAGACGCGCUAGCUUCAGGUGUUAGUGUUCUUACGGACGUGGGGGUUCAAGUCCCCCCCCUCGCACCA
DD# D KP P TP
leuQ (leuT leuV)
GCGAAGGUGGCGGAAUUGGUAGACGCGCUAGCUUCAGGUGUUAGUGUCCUUACGGACGUGGGGGUUCAAGUCCCCCCCCUCGCACCA
DD# D KP P TP
leuU
GCCGAGGUGGUGGAAUUGGUAGACACGCUACCUUGAGGUGGUAGUGCCCAAUAGGGCUUACGGGUUCAAGUCCCGUCCUCGGUACCA
DD# D KP TP
leuW
GCGGGAGUGGCGAAAUUGGUAGACGCACCAGAUUUAGGUUCUGGCGCCGCAAGGUGUGCGAGUUCAAGUCUCGCCUCCCGCACCA
DD# V KP TP
leuX
GCCGAAGUGGCGAAAUCGGUAGACGCAGUUGAUUCAAAAUCAACCGUAGAAAUACGUGCCGGUUCGAGUCCGGCCUUCGGCACCA
4 D # D P B * P TP
leuZ
GCCCGGAUGGUGGAAUCGGUAGACACAAGGGAUUUAAAAUCCCUCGGCGUUCGCGCUGUGCGGGUUCAAGUCCCGCUCCGGGUACCA
4 D # D P ) * P TP

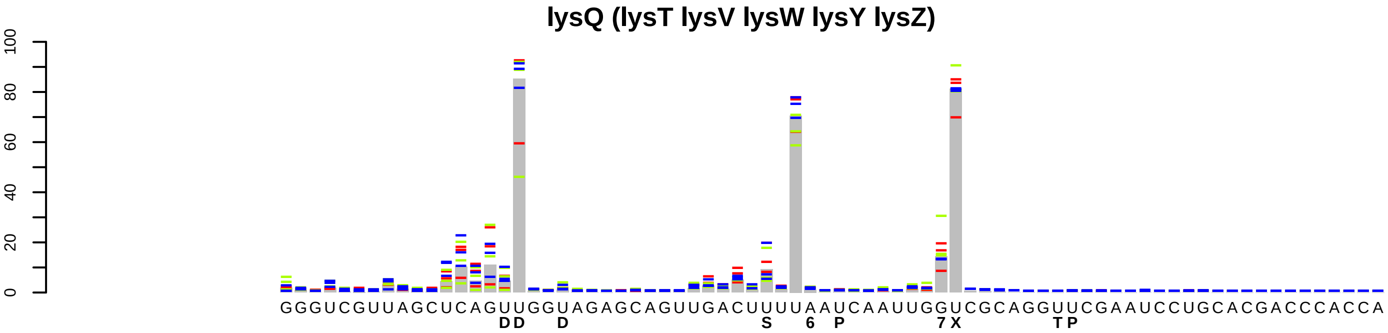
lysQ (lysT lysV lysW lysY lysZ)
0
20
40
60
80
100
GGGUCGUUAGCUCAGUUGGUAGAGCAGUUGACUUUUAAUCAAUUGGUCGCAGGUUCGAAUCCUGCACGACCCACCA
DD D S 6 P 7X TP

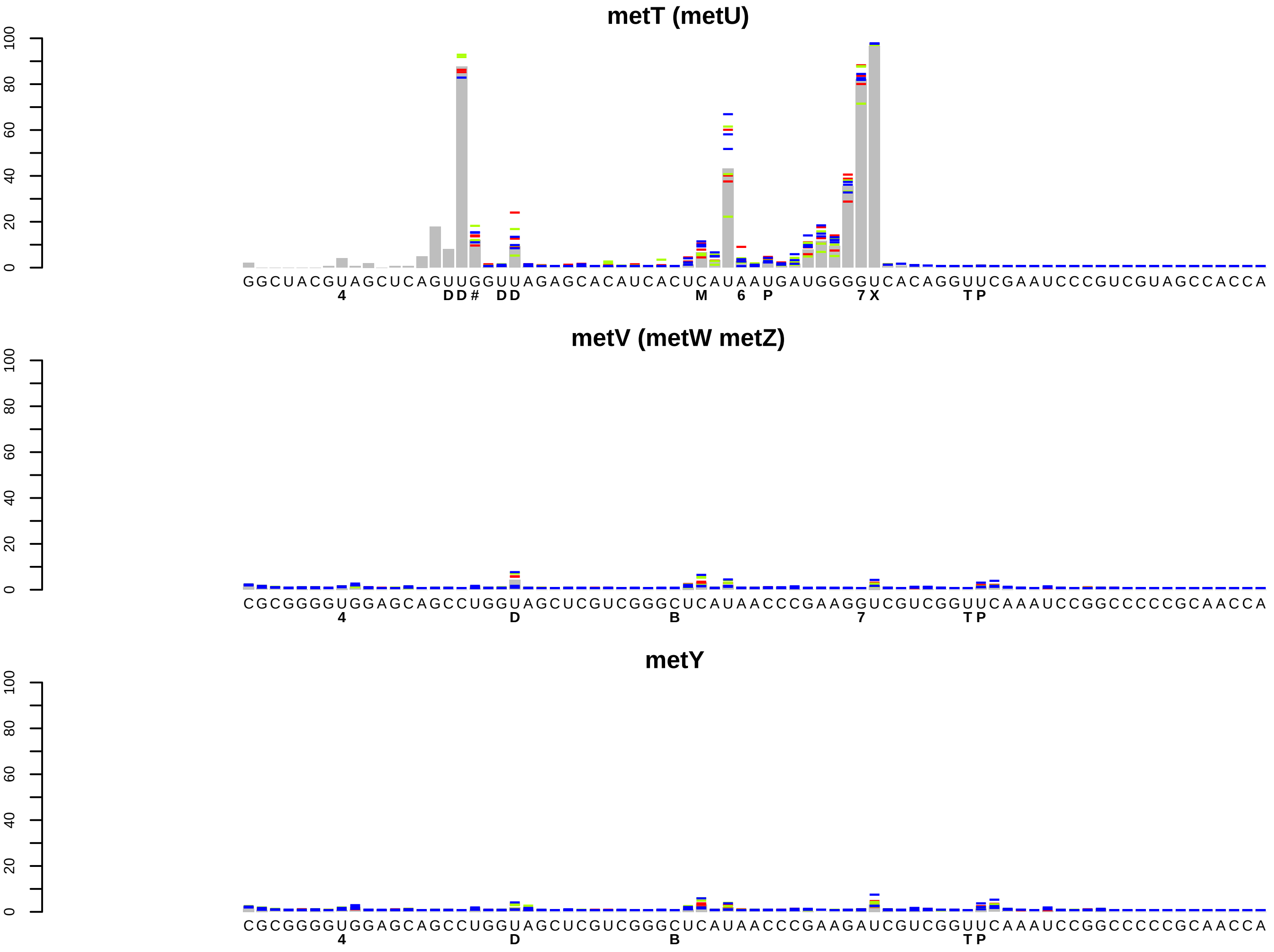
metT (metU)
0
20
40
60
80
100
GGCUACGUAGCUCAGUUGGUUAGAGCACAUCACUCAUAAUGAUGGGGUCACAGGUUCGAAUCCCGUCGUAGCCACCA
4 DD# DD M 6 P 7X TP
metV (metW metZ)
0
20
40
60
80
100
CGCGGGGUGGAGCAGCCUGGUAGCUCGUCGGGCUCAUAACCCGAAGGUCGUCGGUUCAAAUCCGGCCCCCGCAACCA
4 D B 7 TP
metY
0
20
40
60
80
100
CGCGGGGUGGAGCAGCCUGGUAGCUCGUCGGGCUCAUAACCCGAAGAUCGUCGGUUCAAAUCCGGCCCCCGCAACCA
4 D B TP

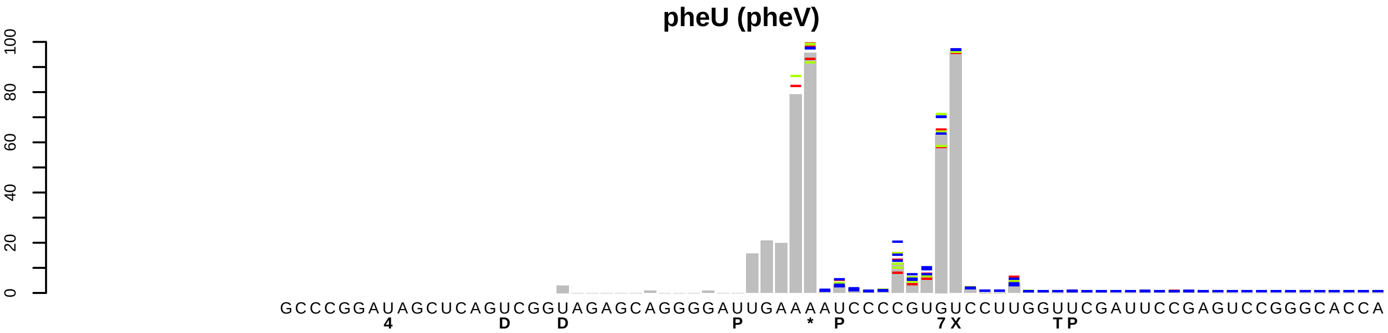
pheU (pheV)
0
20
40
60
80
100
GCCCGGAUAGCUCAGUCGGUAGAGCAGGGGAUUGAAAAUCCCCGUGUCCUUGGUUCGAUUCCGAGUCCGGGCACCA
4 D D P * P 7 X T P

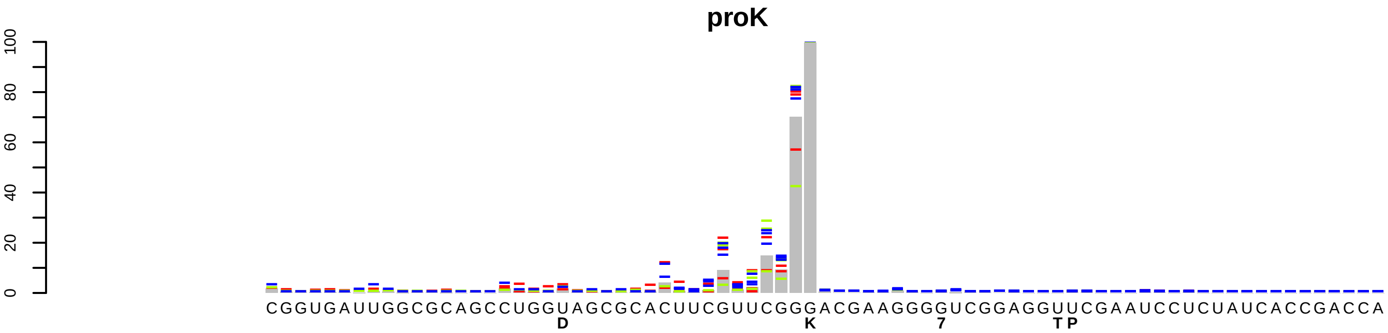
proK
0
20
40
60
80
100
CGGUGAUUGGCGCAGCCUGGUAGCGCACUUCGUUCGGGACGAAGGGGUCGGAGGUUCGAAUCCUCUAUCACCGACCA
D
K
7
TP

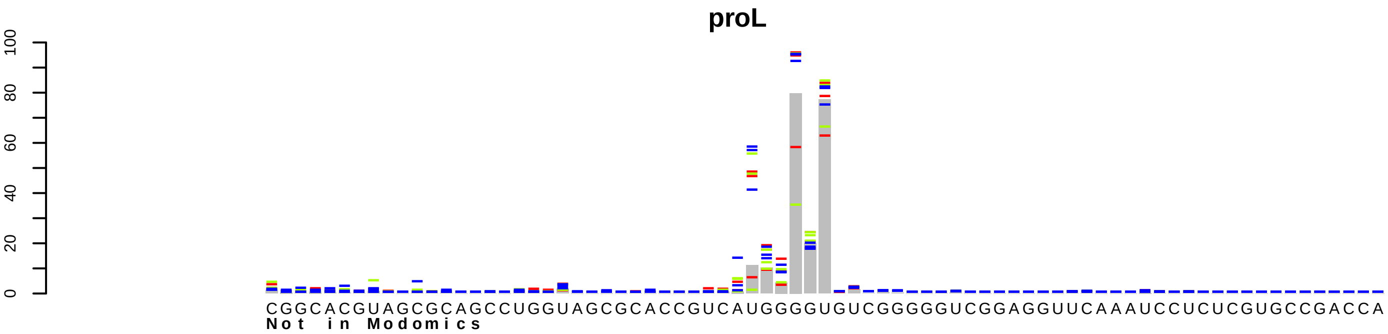
proL
0
20
40
60
80
100
CGGCACGUAGCGCAGCCUGGUAGCGCACCGUCAUGGGGUGUCGGGGGUCGGAGGUUCAAAUCCUCUCGUGCCGACCA
Not in Modomics

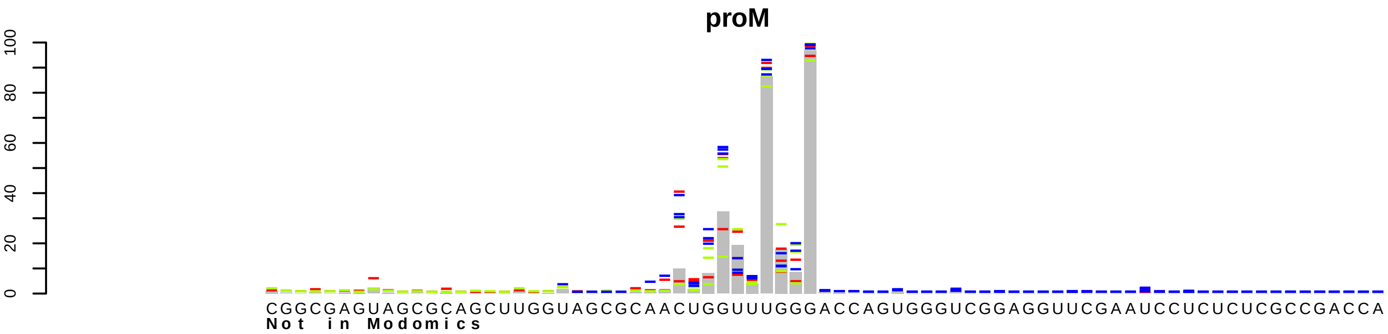
proM
0
20
40
60
80
100
CGGCGAGUAGCGCAGCUUGGUAGCGCAACUGGUUUGGGACCAGUGGGUCGGAGGUUCGAAUCCUCUCUCGCCGACCA
Not in Modomics

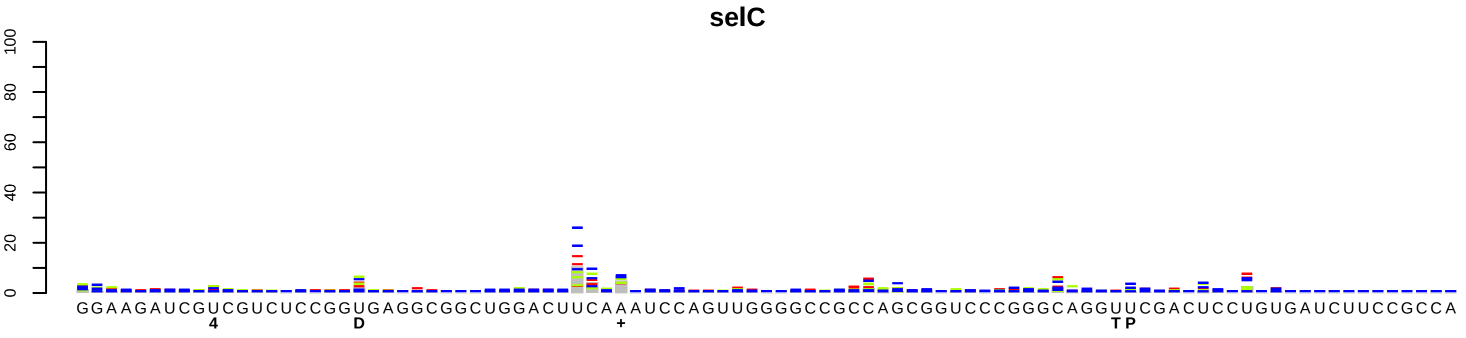
selC
100
80
60
40
20
0
GGAAGAUCGUCGUCUCCGGUGAGGCGGCUGGACUUCAAAUCCAGUUGGGGCCGCCAGCGGUCCCGGGCAGGUUCGACUCCUGUGAUCUUCCGCCA
4
D
+
T P

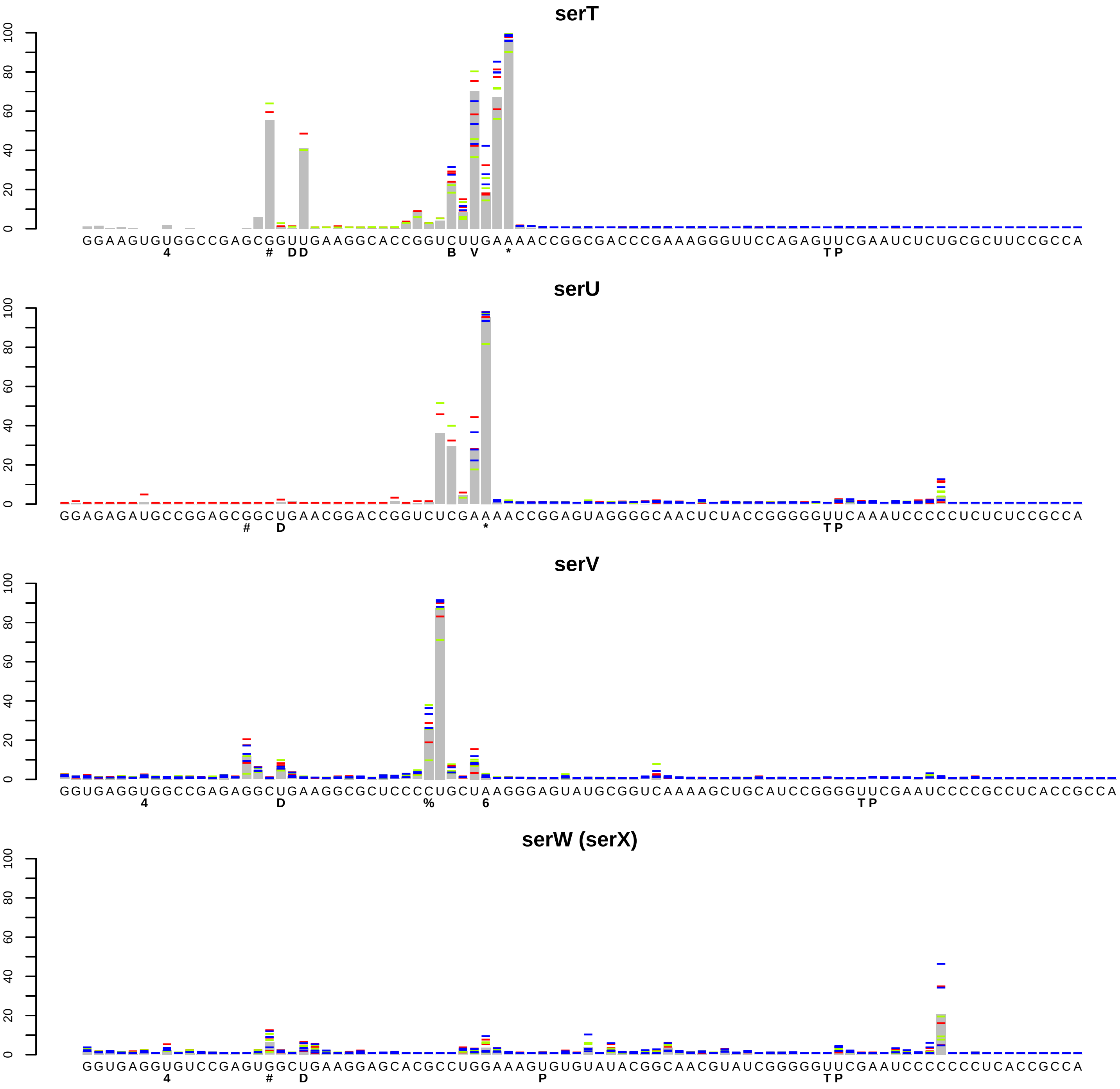
serT
0 20 40 60 80 100
GGAAGUGUGGCCGAGCGGUUGAAGGCACCGGUCUUGAAAACCGGCGACCCGAAAGGGUUCCAGAGUUCGAAUCUCUGCGCUUCCGCCA
4 # D D B V * T P
serU
0 20 40 60 80 100
GGAGAGAUGCCGGAGCGGCUGAACGGACCGGUCUCGAAAACCGGAGUAGGGGCAACUCUACCGGGGGUUCAAAUCCCCCUCUCUCCGCCA
# D * T P
serV
0 20 40 60 80 100
GGUGAGGUGGCCGAGAGGCUGAAGGCGCUCCCCUGCUAAGGGAGUAUGCGGUCAAAAGCUGCAUCCGGGGUUCGAAUCCCCGCCUCACCGCCA
4 D % 6 T P
serW (serX)
0 20 40 60 80 100
GGUGAGGUGUCCGAGUGGCUGAAGGAGCACGCCUGGAAAGUGUGUAUACGGCAACGUAUCGGGGGUUCGAAUCCCCCCCUCACCGCCA
4 # D P T P

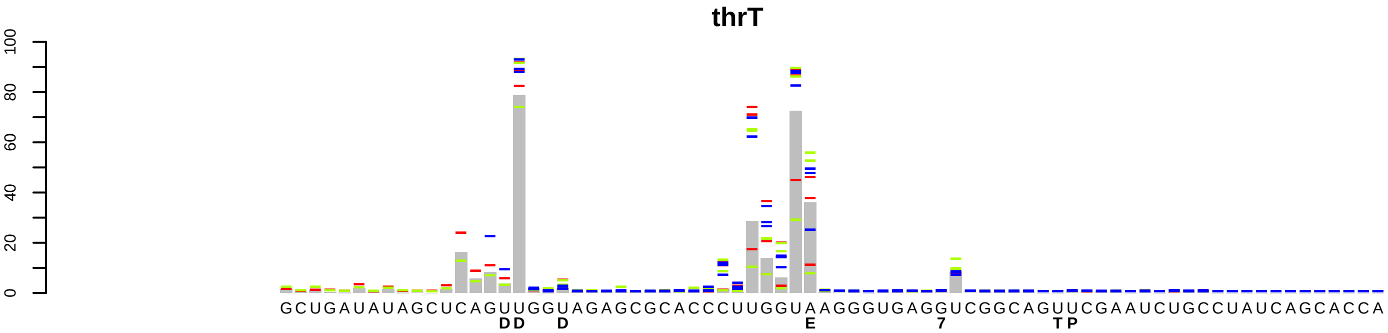
thrT
0
20
40
60
80
100
GCUGAUAUAGCUCAGUUGGUAGAGCGCACCCUUGGUAAGGGUGAGGUCGGCAGUUCGAAUCUGCCUAUCAGCACCA
DD D E 7 TP

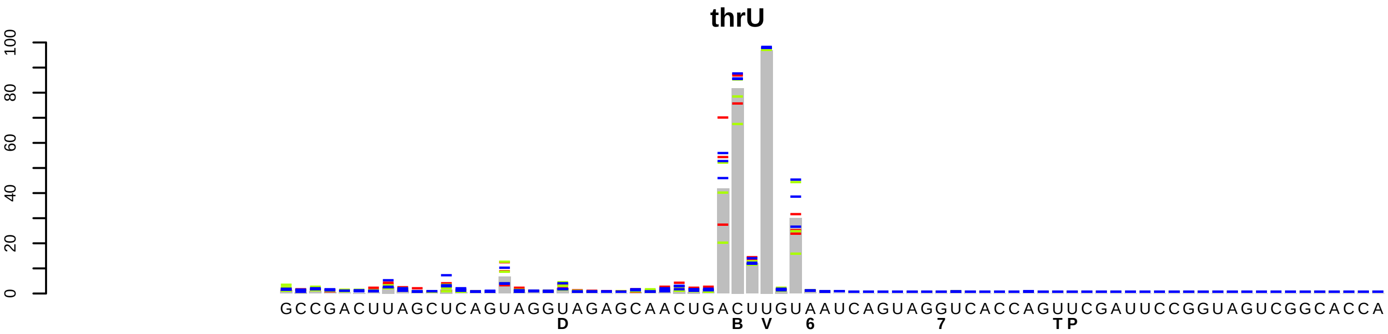
thrU
0
20
40
60
80
100
GCCGACUUAGCUCAGUAGGUAGAGCAACUGACUUGUAAUCAGUAGGUCACCAGUUCGAUUCCGGUAGUCGGCACCA
D B V 6 7 TP

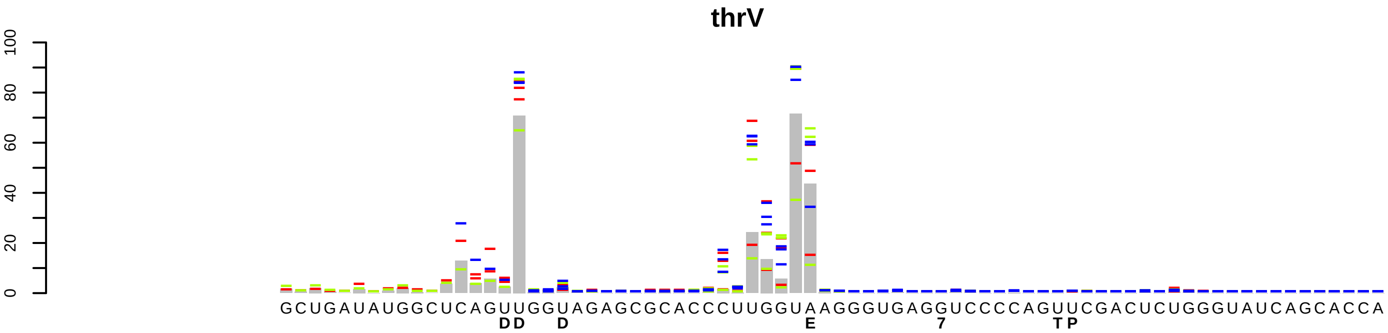
thrV
0
20
40
60
80
100
GCUGAUAUGGCUCAGUUGGUAGAGCGCACCCUUGGUAAGGGUGAGGUCCCCAGUUCGACUCUGGGUAUCAGCACCA
DD D E 7 TP

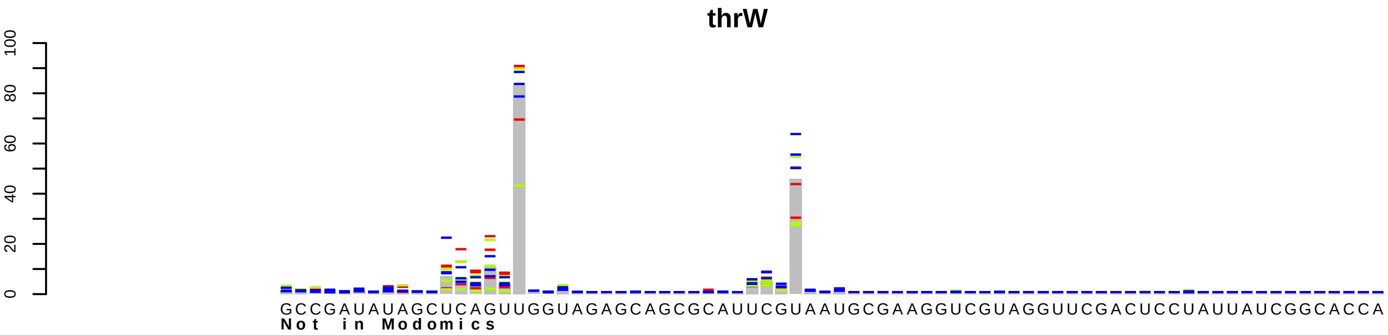
thrW
0
20
40
60
80
100
GCCGAUAUAGCUCAGUUGGUAGAGCAGCGCAUUCGUAAUGCGAAGGUCGUAGGUUCGACUCCUAUUAUCGGCACCA
Not in Modomics

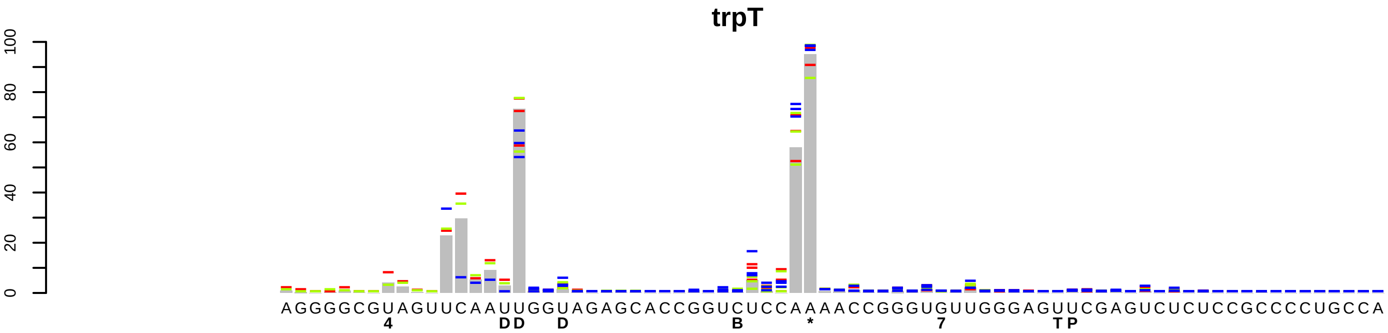
trpT
0
20
40
60
80
100
AGGGGCGUAGUUCAAUUGGUAGAGCACCGGUCUCCAAAACCGGGUGUUGGGAGUUCGAGUCUCUCCGCCCCUGCCA
4
DD
D
B
*
7
TP

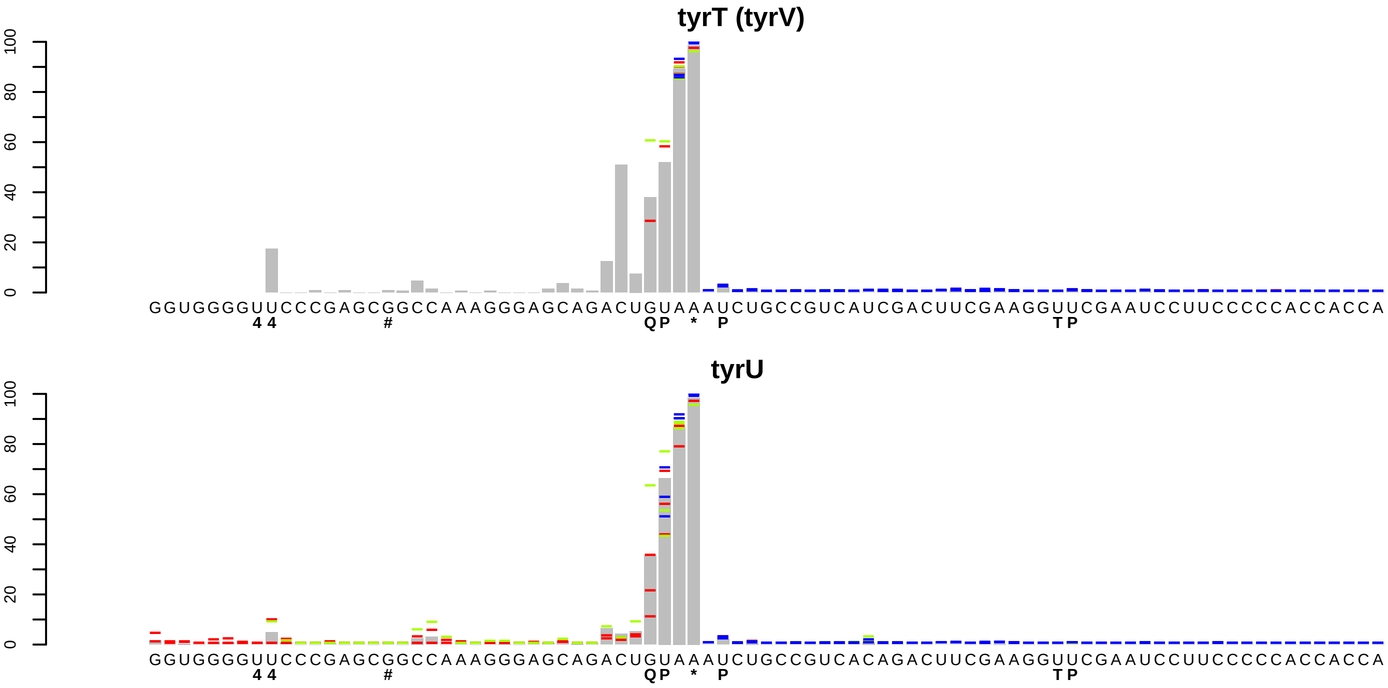
tyrT (tyrV)
0
20
40
60
80
100
GGUGGGGUUCCCGAGCGGCCAAAGGGAGCAGACUGUAAAUCUGCCGUCAUCGACUUCGAAGGUUCGAAUCCUUCCCCCACCACCA
4 4 # Q P * P T P
tyrU
0
20
40
60
80
100
GGUGGGGUUCCCGAGCGGCCAAAGGGAGCAGACUGUAAAUCUGCCGUCACAGACUUCGAAGGUUCGAAUCCUUCCCCCACCACCA
4 4 # Q P * P T P

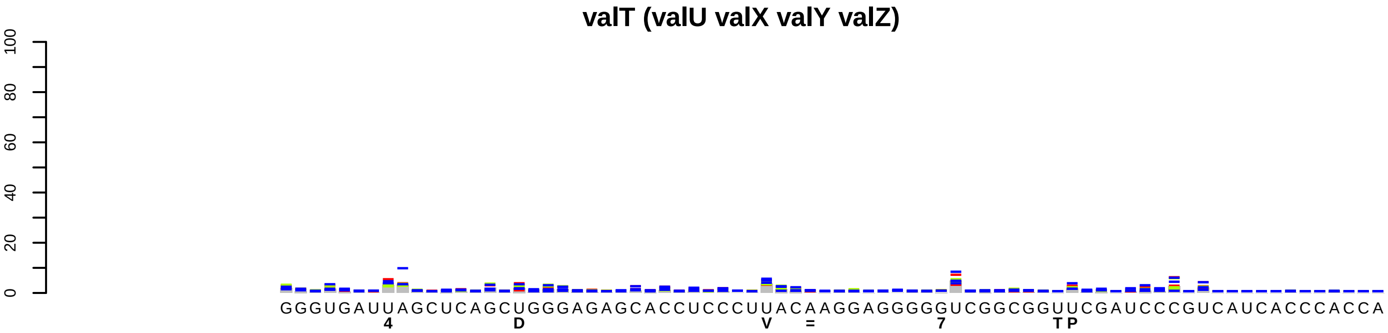
valT (valU valX valY valZ)
0
20
40
60
80
100
GGGUGAUUAGCUCAGCUGGGAGAGCACCUCCCUUACAAGGAGGGGGUCGGCGGUUCGAUCCCGUCAUCACCCACCA
4 D V = 7 TP

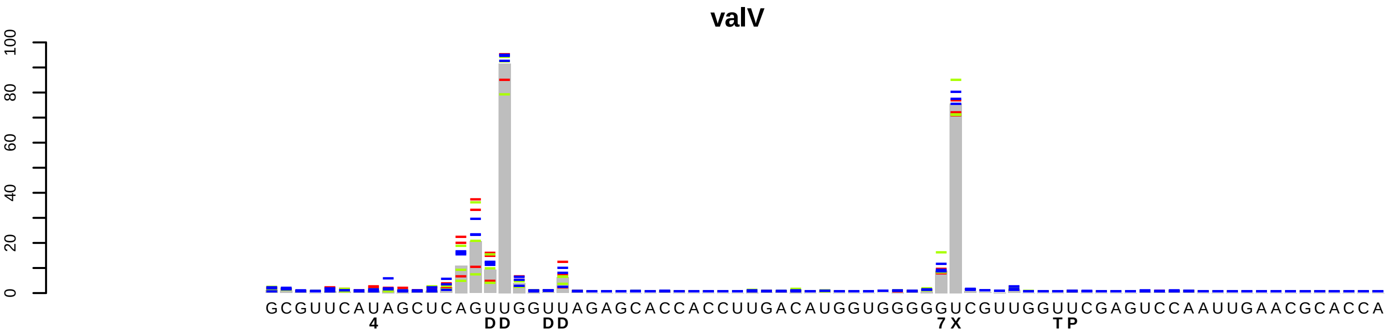
valV
0
20
40
60
80
100
GCGUUCAUAGCUCAGUUGGUUAGAGCACCACCUUGACAUGGUGGGGGUCGUUGGUUCGAGUCCAAUUGAACGCACCA
4 DD DD 7X TP

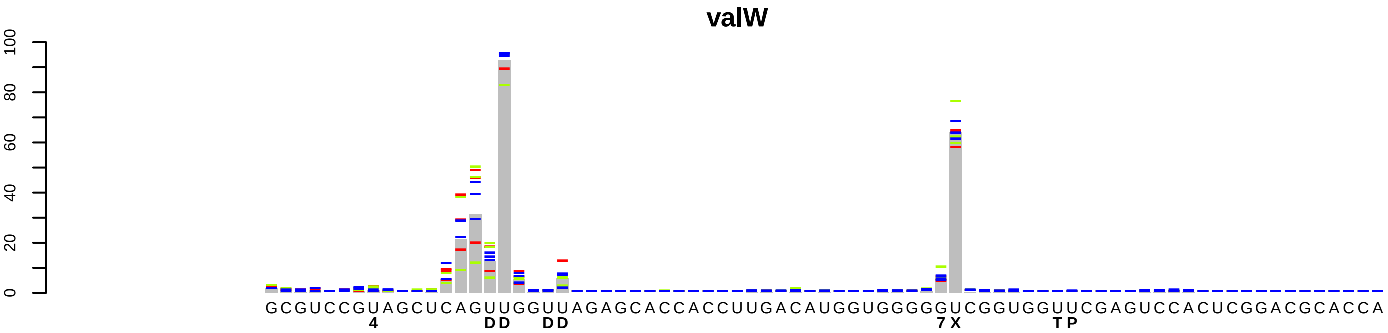
valW
0
20
40
60
80
100
GCGUCCGUAGCUCAGUUGGUUAGAGCACCACCUUGACAUGGUGGGGGUCGGUGGUUCGAGUCCACUCGGACGCACCA
4 DD DD 7X TP

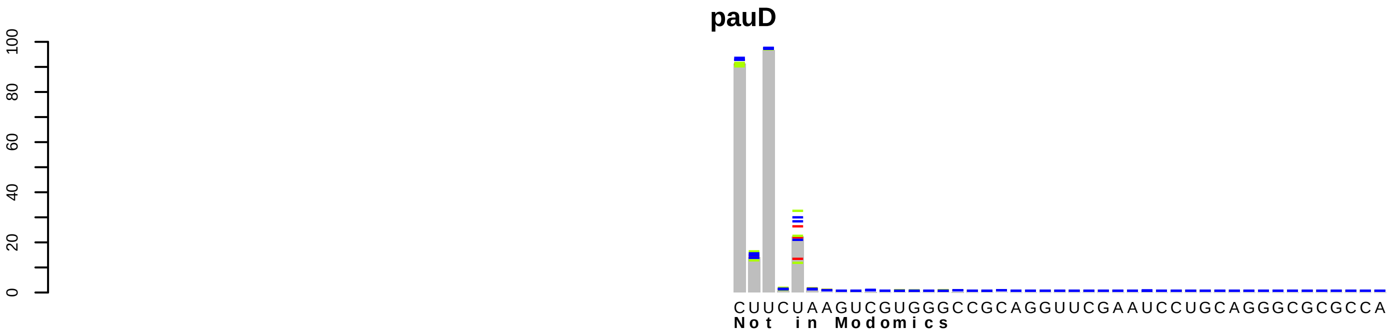
pauD
0
20
40
60
80
100
CUUCUAAGUCGUGGGCCGCAGGUUCGAAUCCUGCAGGGCGCGCCA
Not in Modomics

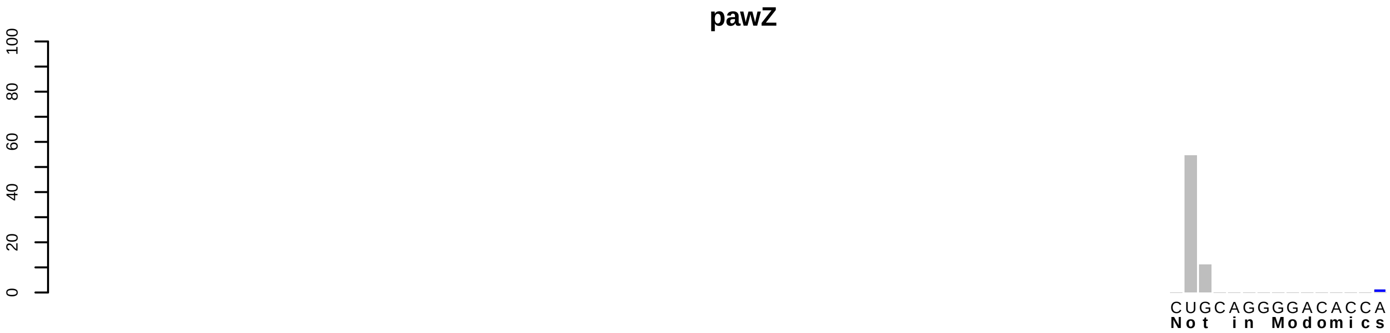
pawZ
0
20
40
60
80
100
CUGCAGGGGACACCA
Not in Modomics

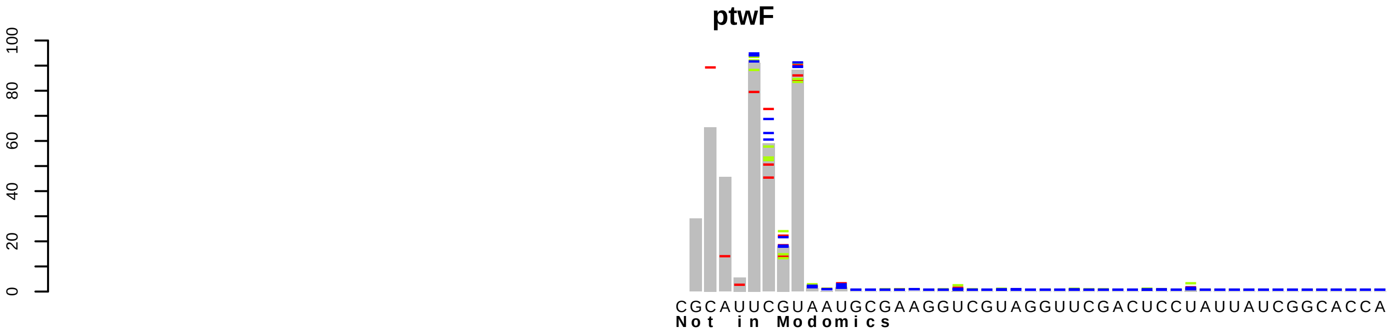
ptwF
0
20
40
60
80
100
CGCAUUCGUAAUGCGAAGGUCGUAGGUUCGACUCCUAUUAUCGGCACCA
Not in Modomics
